# Shaping the structural dynamics of motor learning through cueing during sleep

**DOI:** 10.1101/2024.08.16.608370

**Authors:** Whitney Stee, Antoine Legouhy, Michele Guerreri, Michael-Christopher Foti, Jean-Marc Lina, Hui Zhang, Philippe Peigneux

## Abstract

Enhancing the retention of recent memory traces through sleep reactivation is possible via Targeted Memory Reactivation (TMR), involving cueing learned material during post-training sleep. Evidence indicates detectable short-term microstructural changes in the brain within an hour after motor sequence learning, and post-training sleep is believed to contribute to the consolidation of these motor memories, potentially leading to enduring microstructural changes. In this study, we explored how TMR during post-training sleep affects performance gains and delayed microstructural remodeling, using both standard Diffusion Tensor Imaging (DTI) and advanced Neurite Orientation Dispersion & Density Imaging (NODDI). Sixty healthy young adults participated in a five-day protocol, undergoing five Diffusion-Weighted Imaging (DWI) sessions, pre- and post-two motor sequence training sessions, and after a post-training night of either regular sleep (RS) or TMR. Results demonstrated rapid skill acquisition on Day 1, followed by performance stabilization on Day 2, and improvement on Day 5, in both RS and TMR groups. (Re)training induced widespread microstructural changes in motor-related areas, initially involving the hippocampus, followed by a delayed engagement of the caudate nucleus. Mean Diffusivity (MD) changes were accompanied by increased Neurite Density Index (NDI) in the putamen, suggesting increased neurite density, while Free Water Fraction (FWF) reduction indicated glial reorganization. TMR-related structural differences emerged in the dorsolateral prefrontal cortex (DLPFC) on Day 2 and the right cuneus on Day 5, suggesting unique sleep TMR-related neural reorganization patterns. Persistence of practice-related structural changes, although moderated over time, suggest a lasting neural network reorganization, partially mediated by sleep TMR.

**Statement of Significance:** This study demonstrates how motor practice triggers short-term and long-term microstructural reorganization in (sub)cortical grey matter, with a focus on how sleep Targeted Memory Reactivation (TMR) influences these changes, particularly in motor-related areas. Combining standard Diffusion Tensor Imaging (DTI) with advanced Neurite Orientation Dispersion & Density Imaging (NODDI) imaging techniques, we highlight distinct patterns of neural reorganization, suggesting glial and/or synaptic changes, in response to motor practice, possibly partially mediated by TMR. This research enhances our understanding of motor practice-related structural plasticity mechanisms and highlights the potential of targeted sleep interventions to modulate brain reorganization, suggesting new avenues for optimizing motor skill acquisition.

## 1. Introduction

Upon encountering new experiences, the brain demonstrates a remarkable ability to adjust both its structure and function. These changes result from long-lasting modifications in neural properties, enabling an appropriate response [1], [2], [3]. Noticeable structural brain modifications have been detected following brief periods of motor training [4], [5], [6], using conventional Diffusion Tensor Imaging (DTI) measures. These DTI metrics, i.e., mean diffusivity (MD) and fractional anisotropy (FA), respectively aim at estimating tissue density and fiber organization/directionality. Learning-related MD reductions have been identified in a wide set of cortical and subcortical motor-related regions including the cerebellum [5], [6], hippocampus [4], [5], putamen, caudate nucleus, and thalamus [5], as well as occipital [5], parietal [4], [5], and temporal regions, along with the motor cortex [5], [6]. These findings suggest tissue densification following training in regions involved in motor learning that could, at least partially, be attributable to an increase in neurite density [5], as evidenced using the Neurite Orientation Dispersion & Density Imaging (NODDI; see [7]) biophysical model. Furthermore, changes detected in the precuneus persisted over 24 hours, but a reversal of the hippocampal changes (thus, going from an initial decrease to a significant increase in MD) was observed within the same timeframe [4]. Additionally, both increases and decreases in MD were observed in restricted areas of the parietal and temporal cortices three days following motor training [5], emphasizing the temporal and spatial dynamic nature of motor learning and consolidation-related microstructural modifications.

Besides, sleep has been shown to play a pivotal role in memory consolidation by promoting enduring changes in neural networks [8]. While sleep seems to robustly enhance the consolidation of explicit motor sequence tasks [9], [10], [11], mixed results have been found for more implicit visuomotor tasks: some studies support the hypothesis of sleep-dependent processes [12], [13] while others do not [14] (for review see [15]). One of the mechanisms proposed to underlie the reorganization of neural networks [16] and long-term memory processes [17], [18] is the spontaneous reactivation of recent memory traces during subsequent sleep. In the motor domain, neuronal replay has been observed in rodents during subsequent NREM sleep. Notably, the extent of this reactivation was found to correlate with performance improvement following sleep [19]. Similarly in humans, the reactivation of learning-related neural networks during subsequent sleep has been observed following motor skill acquisition, both in non-rapid eye movement (NREM) [20], [21], [22], [23] and rapid eye movement (REM) sleep [24], [25]. More specifically, the striato-cerebello-cortical network, activated during training, was found reactivated during NREM sleep [20], [21]. Reactivation levels correlated with subsequent performance enhancements on the following day [10]. Furthermore, a reliable relationship has been evidenced between brain structural metrics and sleep oscillation features reflecting brain plasticity processes, such as sleep spindles [26], [27], [28], slow-wave activity [29] or the slope of slow oscillations [30] during NREM sleep. These brain structural measures encompass inter-individual differences in grey matter (GM) volume [26], [27], [29], [31], GM cortical thickness [32] or white matter (WM) diffusion [27], [28]. These findings suggest that, in addition to mirroring the dynamics of neuronal networks, sleep oscillations may, to some extent, reflect and be influenced by the microstructure of their localized brain sources.

Given the strong connection between spontaneous neural reactivation during subsequent sleep and the facilitated consolidation and retrieval of recently acquired memories, it was proposed to further optimize this process using Targeted Memory Reactivation (TMR [33]; for reviews, see [34], [35]). TMR involves presenting sensory cues during sleep that were previously associated with specific information learned while awake. Studies demonstrated that sleep TMR can lead to a distinct improvement in performance for the cued information, compared to information not cued overnight [36]. For instance, in the motor domain, specific performance improvements were observed on the next day after presenting motor learning-related olfactory [37], [38] or auditory [39], [40], [41], [42], [43] cues during subsequent sleep. Various strategies have been adopted for cueing, some exclusively targeting NREM sleep stage 2 (N2) [37], [38], others focusing on NREM sleep stage 3 (N3) [39], [41], [42], or delivering cues during both sleep stages [40], [43]. Also, following training on a Serial Reaction Time Task (SRTT), TMR was shown to enhance the explicit recall of cued sequences over non-cued ones [41], a result that was replicated but found to be rather gender-specific, benefiting men exclusively [44]. Additionally, TMR has been associated with the modulation of neural oscillatory activity, the presentation of learning-related cues leading to a significant increase in sleep spindle activity [37], [43], particularly within the parietal regions [37]. This enhancement in spindle response has been correlated with the behavioral benefits of TMR [37], [39]. Cueing also appears to amplify the occurrence of slow oscillations (SOs), time-locked to spindles [38], [43]. Notwithstanding, the observation of such spindle modulation has not been consistent across studies, even when using similar protocols [38]. At the functional level, increased learning-related activity has been detected in the caudate nucleus and hippocampus for cued sequences, compared to uncued sequences, with this activity being associated with time spent in slow wave sleep (SWS). This was parallelled by heightened functional connectivity between these regions for the cued sequences. In contrast, increased functional TMR-related activity in the DLPFC, as well as in other cortical and cerebellar regions, was related to time spent in REM sleep [42]. These findings suggest that TMR can not only influence the neural oscillations associated with memory reactivation during sleep, but also modify functional activity within motor-related brain regions at subsequent wake. These changes are closely linked to the observed offline performance improvements [42], suggesting a more effective consolidation of motor memories. Furthermore, since transient adaptations in functional connectivity can influence structural connections, which in turn impacts functional connectivity [45], TMR might not only influence functional activation/connectivity and sleep oscillations, but also eventually result in accelerated and/or enhanced learning-related changes in the brain microstructure underlying these functional changes.

In the present study, we aimed to determine to what extent TMR during post-training sleep is associated with immediate (next day) and delayed (3 days later) post-sleep microstructural changes, using a motor sequential Serial Reaction Time Task (SRTT — adapted from [46], [47]). Auditory cues were delivered during stable NREM stage (N2) and stage 3 (N3) in the TMR condition, as compared to regular sleep (RS) where no sounds were delivered. Besides standard DTI, we used the multicompartment analysis method NODDI [7] that offers a more nuanced analysis of microstructural changes, with a delineation of the underlying microstructural properties of the brain tissue. Indeed, NODDI provides three valuable indices: the neurite density index (NDI), reflecting the density of both axons and dendrites; the orientation dispersion index (ODI), characterizing the angular variability of neurites; and the free water fraction (FWF), indicating the proportion of cerebrospinal fluid (CSF) and separating its contribution from the overall signal. Hereby, NODDI offers enhanced specificity compared to the traditional DTI model. We hypothesized that the TMR group will exhibit an accelerated structural reorganization correlating with behavioral performance gains as compared to the RS group, with significant differences in diffusion metrics following the experimental night, that may or not persist 3 days later. At the subcortical level, we focused our analysis on 6 bilateral subcortical regions of interest (ROIs) based on the SRTT literature and motor studies using DTI, i.e., the Cerebellar Cortex [4], [6], [48], [49], Thalamus [48], [49], Hippocampus [4], [50], [51], Caudate, Putamen, and Pallidum [48], [49]. In contrast, for the cortical analysis, we adopted a whole brain approach due to the variability in the literature regarding the exact cortical regions and networks involved.

## 2. Methods

### 2.1. Participants

Sixty young, healthy participants (30 females), aged 18-29 years (mean age ± SD = 21.58 ± 2.17), gave written informed consent before taking part into this study approved by the Liège University Hospital Ethics Committee (approval #2020/138). All participants were without neurological or psychiatric history, had a body mass index (BMI) below 28, and no contra-indication to EEG and MRI. Their chronotype was moderate to neutral (mean Morningness– Eveningness Questionnaire [52] score ± SD = 54.28 ± 7.10; ranging 32 - 68) and they reported a good sleep quality (mean Pittsburgh Sleep Quality Index [53] score ± SD = 3.47 ± 1.40, ranging 0 - 6). Musicians and computer scientists (with potentially superior hand motor dexterity due to practice) as well as smokers and individuals who experienced jetlag in the previous 3 months were excluded from the study. Both right- and left-handers were included (mean Edinburgh Inventory [54] score ± SD = 4.85 ± 6.60, ranging -10 - 10) considering the bimanual nature of the motor task (see below) and the lack of consistent structural asymmetries associated with handedness (see [55], [56] but [57]). Participants were pseudo-randomly assigned to either the Regular Sleep (RS) group, or the TMR sleep (TMR) group, both counting 15 women and 15 men (see Supplemental Information (SI) section 7, Table S1 for details).

### 2.2. Motor Task

In our study, we employed a 6-choice version of the SRTT (see Figure 1, adapted from [46], [47]), coupled with auditory tones, implemented on PsychoPy3 v2020.2.10 (Nottingham, UK). Participants were seated in front of the computer screen and asked to place their 6 middle fingers (index, middle and ring fingers of each hand) on the 6 response keys corresponding to the 6 squares horizontally positioned on the screen. The task required participants to press as fast and as accurately as possible the key associated with the square where a visual cue appeared. Each key press produced a beep tone linked to that specific key/position (1-C, 2-C#, 3-D, 4-D#, 5-E, 6-F), and the subsequent trial appeared 500 milliseconds later. One block consisted of 96 trials, with each block separated by a brief, self-timed rest period. The trials in each block either followed a 12-element sequence (5-3-1-6-2-4-1-5-2-3-6-4) or were a pseudo-random succession of cues (with the rule that the same position could never been displayed twice in a row). The initial learning session included 30 blocks (with block 26 being random), the next morning’s retest comprised 2 sequential blocks (31-32), and the relearning session counted 20 blocks (33-52; with blocks 35 and 48 being random). Random blocks were included to distinguish sequence-dependent learning from sequence-independent learning (or mere motor-related learning), both contributing to performance improvement [58]. Reaction times typically increase, and accuracy decreases, in random blocks due to the unpredictability of the next stimulus, indirectly indicating the learning of regularities in the sequential blocks. Participants were not informed beforehand about the sequential nature of the task. Consequently, we included a generation task at the end of the procedure to assess explicit knowledge. In the first block, participants were asked to generate the sequence from memory. In the second block, they were required to perform the sequence in reverse order (see SI, section 1 for details & results).

**Figure 1.**
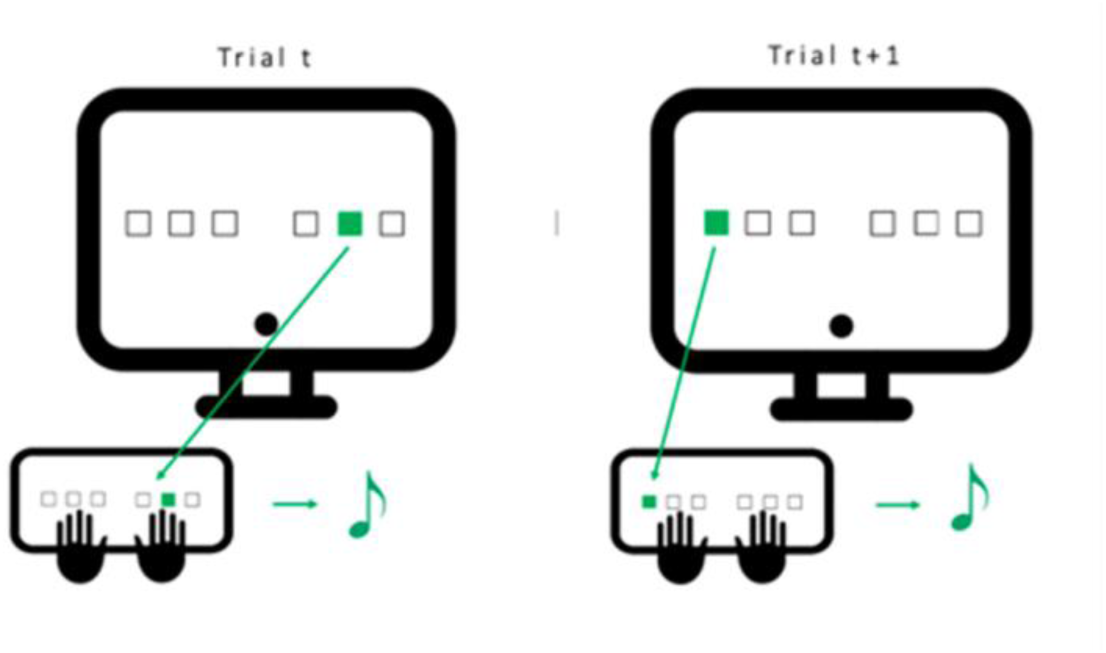
Serial Reaction Time Task (SRTT) Participants are seated in front of the computer screen, with the index, middle, and ring fingers of both hands positioned on the six keys corresponding to the six positions on the screen. They are asked to press the key that matches the cued position on the screen as fast and as accurately as possible. Following a key press, a specific tone associated with the key is played, and the subsequent visual cue appears after a 500-msec delay. A single block is composed of 96 cues, arranged in either a repeated 12-element sequence or in a pseudo-random order.

### 2.3. General Procedure

Participants were instructed to avoid caffeine and other stimulants on testing days, and to keep a consistent sleep-wake pattern throughout the experiment. Sleep schedule regularity was monitored using self-reported sleep logs (St. Mary’s Hospital sleep questionnaire [59]) and visual inspection of actimetric recordings (ActigraphTM wGT3X-BT, Pensacola, FL, USA) from 3 days prior the first testing day until the end of the procedure. To control for hormonal biases on motor performance, sleep, and consolidation, female participants were tested during their luteal phase only [60], [61].

Figure 2 illustrates the experimental design. Three days prior to the initial testing day (Day 1), participants underwent a habituation night of sleep in the lab under 256-channel high-density EEG (hd-EEG). On Day 1, around 16:30, participants underwent their first diffusion-weighted MRI (DWI1). Immediately afterward, they completed 30 SRTT blocks (duration about 1 hour) (see details section 2.2.). Half an hour following the end of the learning episode, a post-training diffusion-weighted MRI (DWI2) was conducted. Participants were then assigned to either the RS or TMR condition for the post-learning night (but were not informed of their group assignment). Around 21:30, both groups were equipped with the hd-EEG cap before going to sleep for the entire night at their regular bedtime in the laboratory (approximately 8-9 hours).

**Figure 2.**
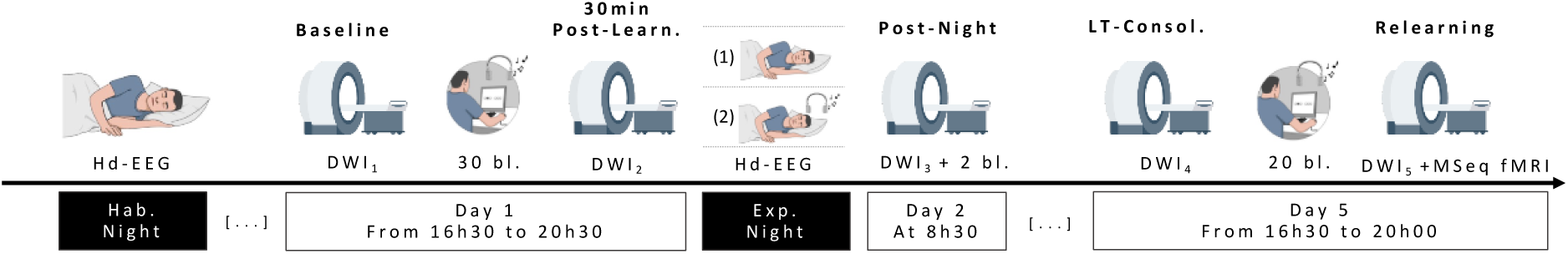
Experimental design. Three days prior the initial testing day, all participants came to the lab for a habituation night under hd-EEG. On Day 1, they underwent a first diffusion MRI session (DWI_1_), followed by approximately 1h (30 blocks) of motor sequence learning on the SRTT, where each response key was associated with a specific auditory tone. Thirty minutes after the end of the learning episode, a second diffusion MRI session (DWI_2_) was conducted. During the subsequent night, naïve subjects either had a regular night of sleep (1; RS) or were exposed to learning-related auditory cues during sleep stage N2 & N3 (2; TMR), under hd-EEG. On the next morning, all participants underwent a third diffusion MRI session (DWI_3_) and a brief behavioural retest (2 blocks). After spending 3 nights of regular sleep at home, participants returned for a fourth diffusion MRI session (DWI_4_), followed by a 40min retraining session on the SRTT (20 blocks), and then a final diffusion MR session (DWI_5_).

Regarding the TMR procedure, we first calibrated the volume of the sounds to align to each participant’s sensitivity prior bedtime. To this end, we played the sounds in a random order and asked the participant to indicate the maximal volume at which they would continue to sleep undisturbed, should the sounds be played during the night. Then, in the TMR condition, once the participant was asleep, auditory cues were delivered during stable N3 for the first sleep cycle and during both N2 and N3 for the second cycle. As each response key of the 12-element SRTT was associated with a distinct auditory tone (see section 2.2.), we utilized the "melody" heard during practice as overnight cue. Hence, we replayed the 12 tones, each lasting 500 msec, to create a 6-second auditory sequence. Each 6-sec stimulation was followed by a 6-sec silent period before the next set of auditory cues was delivered. Presentation of the learning-related cues was interrupted whenever signs of arousal or a transition to a non-targeted sleep state were detected in the polysomnographic recording. Stimulation was then resumed once the participant returned to stable N2 or N3.

On Day 2 at around 08:30, both groups underwent a third diffusion-weighted MRI (DWI3) before performing a short SRTT retest (2 blocks) outside of the scanner. They were then sent home and instructed to maintain a regular sleep pattern. On Day 5 around 16:30 (i.e., at the same time than on Day 1 to control for circadian influences [62], [63]), a fourth diffusion-weighted MRI (DWI4) was acquired. Participants were then retrained on the SRTT for 20 blocks (approximately 40 min), followed by a fifth MRI (DWI5). A 5-minute version of the Psychomotor Vigilance Test -PVT [64] was administered before each SRTT session to track possible changes in behavioural alertness. The RS dataset has been previously used in a prior publication, where the RS condition was compared to an experimental night involving sleep deprivation (see [5]).

### 2.4. Behavioural data analyses

At the behavioural level, SRTT performance was assessed for each block computing the Global Performance Index (GPI) [37] (see SI, section 2 for detailed formula). The GPI (ranging 0 to 1, 1 being the ideal performance) accounts for individual strategies by considering both accuracy and speed. Indeed, some individuals may focus on speed, sacrificing accuracy, while others may prioritize error-free sequences. Such strategic variations can lead to minor differences on an individual basis, rendering either speed or accuracy individually less reliable for assessing online performance and offline consolidation [37]. For the sake of completeness, detailed separate analyses on mean reaction time and accuracy can be found as supplemental information (SI, section 3).

Frequentist statistics were computed using JASP version 0.15 (JASP Team (2021)). Welch t-tests and Welch ANOVAs were always preferred to Student t-tests and classical One-way ANOVAs considering their increased power in case of heterogeneity of variance, that Levene’s test for equality of variances often fails to detect [65], [66]. When normality was violated, Mann–Whitney U-tests were performed. Degrees of freedom were corrected with Greenhouse–Geisser sphericity correction in case Mauchly’s sphericity test indicated violated assumption. Bonferroni correction for multiple comparison was applied when post-hoc tests were conducted. All tests are based on a two-sided significance level set at *p* < 0.05.

### 2.5. MRI Data Acquisition & Processing

The MRI acquisition parameters and the processing pipeline were overtaken from [5].

#### 2.5.1. MRI Data Acquisition

MR data were acquired on a Siemens Magnetom Prisma 3T (software: Syngo MR E11) scanner. High resolution structural images were acquired for anatomical reference. Parameters for the 3D T1-weighted magnetization-prepared rapid gradient echo (MPRAGE) were acquisition time = 4 min 10 s, echo time (TE) = 2.19 ms, repetition time (TR) = 1900 ms, inversion time (TI) = 900 ms, flip angle = 9°, voxel size = 1 × 1 × 1 mm^3^, and matrix dimensions = 224 × 240 × 256 (sagittal, coronal, axial). For the 3D T2-weighted spin-echo, acquisition time was 8 min 27 s, TE = 5.66 ms, TR = 3200 ms, flip angle = 120°, voxel size = 0.7 × 0.7 × 0.7 mm^3^, and matrix dimensions = 256 × 320 × 303 (sagittal, coronal, axial). Multi-shell diffusion acquisitions were composed of 13 b = 0 and diffusion-weighted images with b-values 650, 1000 and 2000 s.mm^-2^, respective number of directions = 15, 30, 60. For distortion correction purpose, two sets of DWI acquisitions were acquired with the same settings except for the phase encoding direction (PED) that was reversed - antero-posterior (AP) and postero-anterior (PA). For the 2D axial spin-echo echo-planar imaging used for DWI, acquisition time (for one set of DWIs) was = 8 min 12 s, TE = 70.2 ms, TR = 4070 ms, flip angle = 90°, voxel size = 2 × 2 mm^2^, slice thickness = 2 mm, slice dimensions = 96 × 96 (sagittal, coronal), number of slices = 70. Lastly, for the task-based fMRI, multi-slice T2*-weighted functional images using axial slice orientation and covering the whole brain were acquired with gradient-echo echo-planar imaging (EPI), TE = 30 ms, TR = 2260 ms, flip angle = 90°, voxel size = 3 × 3 × 3 mm^3^, 25% interslice gap, number of slices = 36, matrix dimension = 72 × 72 × 36.

#### 2.5.2. MRI Data Processing

##### 2.5.2.1. Anatomical processing

Raw T1-weighted images were corrected for bias field signal using the BiasFieldCorrection_sqrtT1wXT2w script from https://github.com/Washington-University/HCPpipelines/tree/master, as described in the minimal preprocessing pipelines for the Human Connectome Project [67]. Segmentation was performed on the T1-weighted images using FastSurferCNN [68], an advanced deep learning model trained to replicate Freesurfer DKT’s segmentation. FastSurferCNN segments the brain into 95 cortical and subcortical regions following the Desikan-Killiany-Tourville protocol [69], [70]. Cortical surface reconstruction was performed on T1-weighted images using FastSurfer [68] which is an extensively validated pipeline to efficiently mimic Freesurfer recon-all [71], [72], [73] by leveraging FastSurferCNN output.

##### 2.5.2.2. Diffusion preprocessing

The susceptibility distortion field was estimated through registration of the raw AP and PA reversed phased encoded b=0 volumes using FSL TOPUP [74]. Eddy-current distortion and head motion parameters have been estimated using FSL EDDY [75]. The reconstruction of the undistorted DWI volumes from all the raw AP and PA reversed phase encoded images was also handled by FSL Eddy following the least-squares restoration approach from [74]. By feeding TOPUP outputs to EDDY, all the distortion and movement parameters were composed to be applied all at once, thus avoiding unnecessary resampling.

##### 2.5.2.3. Diffusion model fitting

The DTI model was fitted through linear least squares using FSL DTIFIT. To limit the effect of non-Gaussian diffusivity which gets stronger with high b-values [76], only the pre-preprocessed DWI volumes with b-values 0, 650 and 1000 s.mm^-2^ have been used for the fitting. The revised version [77], [78] of the original NODDI model [7] was fitted using the NODDI matlab toolbox (http://mig.cs.ucl.ac.uk/index.php?n=Tutorial.NODDImatlab). All the preprocessed DWI volumes were used for the fitting.

##### 2.5.2.4. Diffusion to anatomical mapping

The diffusion maps in subject native diffusion space were mapped to the high-resolution subject native anatomical space through rigid boundary-based registration [79] of the estimated b=0 image onto the T1-weighted image using FSL EPI_REG script. Diffusion metric statistics were then extracted in this native anatomical space.

##### 2.5.2.5. ROI-wise diffusion metrics extraction

To reduce the bias associated with cerebrospinal fluid (CSF) partial volume contamination when using conventional mean, we instead used the FWF estimated from NODDI to compute a tissue-weighted (“tw”) mean [80] for each ROI as summary statistic.

##### 2.5.2.6. Surface-wise diffusion metrics extraction

The following processing was performed using the FreeSurfer suite and outputs from the cortical surface reconstruction. Using mri_vol2surf, the diffusion metrics volumes were projected onto the mid-cortical surface, halfway through the white-grey matter border and the pial surface. Then, a smoothing kernel of FWHM 6 mm was applied along the mid-cortical surface, thus properly following the gyri and sulci circumvolutions, which usual volumetric smoothing does not allow. Surfaces of all subjects were then aligned onto a common surface template using mris_preproc.

##### 2.5.2.7. dMRI data analyses

For the ROI-based statistical analysis, we performed multivariate analysis of variances (MANOVA) separately on DTI (twMD, twFA) and NODDI (twNDI, twODI, FWF) parameters using SPSS version 28.0.0.0. Significance level was set at 0.008 to correct for multiple comparisons (0.05/6 ROIs). Post-hoc univariate ANOVAs were performed when necessary (significance level set at 0.05). Correlations between changes in DWI metrics for each ROI between two timepoints and behavioural parameters or sleep parameters were also computed (p < 0.05 threshold; Pearson’s *r*, or Spearman’s *ρ* when normality assumptions are violated). For behavioural scores, we calculated proportional online gains for all participants (computed as the difference in RT between the mean of the 2 last blocks of the learning/relearning session minus the mean of the 2 first blocks of the learning/relearning session, divided by the mean of the 2 first blocks of the learning/relearning session and multiplied by 100). Complementary post-hoc MANOVAs on the hippocampus and the caudate were performed with a significance level set at 0.025 (0.05/2 ROIs). For sleep parameters, we used normalized sleep stage duration (NREM2, NREM3, REM) during the experimental night.

The surface-based statistical analysis was also conducted using the FreeSurfer suite. For each chosen contrast, a general linear model (GLM) was fitted on the mris_preproc outputs using mri_glmfit (different onset, different slope), and two-tailed significance for t-statistic was computed for the estimated parameters at each vertex. We performed a one-sample t-test across the entire sample to assess the significance of the mean difference in measurements between 2 distinct timepoints (*Learning/Day/Relearning* effects). We then compared between groups the mean differences between timepoints (*Learning/Day/Relearning* effects by *Conditions*), using an unpaired t-test. To account for multiple comparisons, a cluster-wise correction based on permutations [81] was performed using mri_glmfit-sim. We set 1000 permutations, a vertex-wise cluster-forming p-value (*p*) threshold at *p* < 0.001, and a cluster-wise p-value (CWP) threshold at CWP < 0.05. We also computed Pearson’s correlation maps between changes in cortical DWI metrics (at each vertex) observed between specific sessions and behavioural or sleep parameters (p < 0.05 threshold), calculated as described hereabove. We did not include FA in the cortical ribbon analysis as it is not suited for GM exploration.

### 2.6. Hd-EEG Data Acquisition & Processing

EEG data were recorded using a 256-channel high-density EEG system (Geodesic EEG System, Electrical Geodesics, Inc.). The electrode net was composed of microcel sensors that conformed to the participant’s head to ensure optimal scalp coverage. The EEG sampling rate was 250 Hz, with a band-pass filter set from 0.1 to 100 Hz. Electrode impedances were all below 70 kΩ at the beginning of the night recording. EEG data were recorded throughout the entire night, and the presentation of the acoustic stimulations was marked on the EEG recording. Sleep scoring according to the AASM guidelines [82] and artifact rejection was performed using visual inspection on a subset of electrodes located at positions corresponding to F3, F4, C3, C4, O1, alongside 2 EOG and 1 EMG channels, ensuring that data contaminated by arousals were excluded from further analysis. In this paper, EEG data were used to monitor sleep quality and efficiency, as well as to compute basic sleep parameters. A more detailed analysis of hd-EEG data will be presented in a separate publication.

## 3. Results

### 3.1. Demographic Data

Welch ANOVAs performed separately on age, laterality, sleep quality and chronotype (see SI, section 7, Table S1) with between-subject factor *Condition* (RS vs. TMR) did not reveal any significant differences between the RS and TMR groups (all *p*s > 0.353).

### 3.2. Sleep Data

Mixed ANOVAs investigating the time spent in each sleep stage - N1, N2, SWS, REM - with within-subject factor *Night* (habituation or experimental) and between-subject factor *Condition* (RS vs. TMR) revealed a significant main *Night* effect for time spent in N1 (*p* < 0.001) and REM (*p* < 0.001), but not for N2 (*p* = 0.648) or SWS (*p* = 0.907). Participants exhibited less N1 but more REM sleep during the second, experimental night as compared to the first, habituation night. A main *Condition* effect was evidenced for SWS only (*p* = 0.018; all other *p*s > 0.207), with an overall longer time spent in SWS for the RS group as compared to the TMR group. However, no *Night*\**Condition* interaction emerged (all *p*s > 0.113). See SI, section 7, Table S2 for details on the time duration spent in each stage. Additional analyses on spindle and slow oscillation density are available in SI, section 4.

### 3.3. Behavioural Data

Detailed results on GPI, mean RT and accuracy are available as supplementary information (SI section 3). On Day 1, GPI progressively increased with task practice (*p* < 0.001; see Figure 3, blocks 1-30). At pseudo-random block 26, a drop in performance was observed (*p_corr_* < 0.001), indicating that participants started anticipating the sequence, demonstrating performance improvement was not merely due to motor practice. As expected, there was no significant effect of *Condition* (*p* = 0.336) or *Block*\**Condition* interaction (*p* = 0.410) at this stage, as the experimental manipulation (TMR vs regular sleep) had not happened yet.

**Figure 3:**
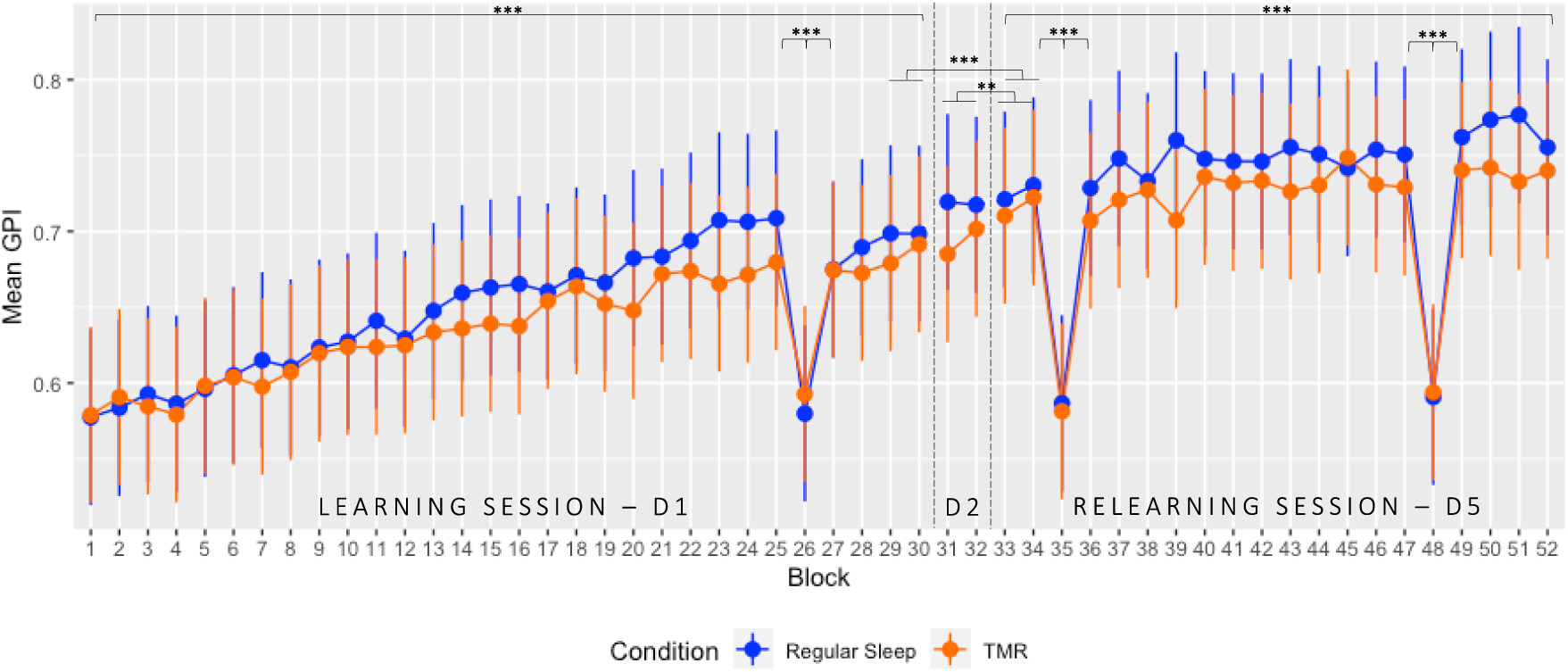
GPI evolution over the entire protocol. Global Performance Index (GPI) ± standard deviation plotted for all blocks executed over the 3 different testing days. On day 1, participants completed 30 blocks in a learning session (D1; block 26 pseudo-random). Following the experimental night taking place between Day 1 & Day 2, performance was assessed in the morning on two blocks (D2; blocks 31-32). After spending three nights of sleep at home, participants performed the task again for 20 blocks in the relearning session (D5; block 35 and 48 pseudo-random). *** p < 0.001; ** < 0.01. The results of the RS dataset have been previously reported [see 5] in comparison with a sleep deprivation condition.

Regarding delayed offline gains, no significant changes in GPI were found between D1 and D2 (*p_corr_* = 0.070), but a significant increase was observed on D5 as compared to D1 or D2 (all *p_corr_s* < 0.033). *Condition* (*p* = 0.375) and *Day*Condition* interaction effects were non-significant (*p* = 0.341), suggesting both groups showed similar delayed improvements on D5, regardless of the experimental TMR manipulation (see Figure 3, blocks 29-34).

Finally, on Day 5, GPI continued to increase over sequential blocks, as reflected by a significant *Block* effect (*p* < 0.001). Pseudo-random blocks 35 and 48 disclosed lower GPI than the surrounding sequential blocks (all *p_corr_*s < 0.001), indicating acquisition of the sequential regularities. *Condition* (*p* = 0.316) and *Block*Condition* interaction (*p* = 0.140) effects were non-significant, suggesting that the presence of auditory cues during the post-learning night (TMR) did not modulate the online gains developing during a relearning session on the same material (see Figure 3, blocks 33-52).

Additional control analyses on alertness and sleep data collected via questionnaires during the protocol are provided in SI, section 5.

### 3.4. Diffusion Weighted Imaging (DWI) Data

#### 3.4.1. Motor training-related short-term structural changes (Day 1; DWI1 vs. DWI2)

First, we investigated the presence of immediate post-initial training changes in brain microstructure by comparing post-(DWI2) and pre- (DWI1) training structural acquisitions, both in the cortical ribbon and specific ROIs on Day 1, prior to the TMR manipulation during subsequent sleep.

##### 3.4.1.1. Cortical Ribbon

*Learning* effects common to both *Conditions* and their correlations are provided in SI, section 6.1.1. As a control analysis, we confirmed that *Learning* effects did not significantly differ between *Conditions* (RS vs. TMR) across all four tested metrics (MD, NDI, FWF, ODI). As expected at this pre-manipulation stage, no differences emerged.

##### 3.4.1.2. Subcortical ROIs

MANOVAs with within-subject factor *Learning* (Pre DWI_1_ vs. Post DWI_2_) and between-subject factor *Condition* (TMR vs. RS) were conducted on DTI parameters for each ROI. Main *Condition* effects were non-significant (all *p_corr_* > 0.033), but there was a significant *Learning*\**Condition* interaction effect in the right caudate (*p_corr_* < 0.001; all *p_corr_* > 0.013 in other ROIs). Post-hoc analyses evidenced decreased MD and increased FA over learning in the RS group, but increased MD and decreased FA in the TMR group (all *ps* ≥ 0.001).

Similar MANOVAs were performed on NODDI parameters in every ROI. Unexpectedly, a main *Condition* effect was found in the right pallidum (*p_corr_* = 0.002) with a significantly lower ODI in the RS group compared to the TMR group (*p* = 0.002) and a significantly lower FWF in the RS group compared to the TMR group (*p* = 0.008). Also, a significant *Learning*\**Condition* interaction emerged in the right caudate (*p_corr_* = 0.002) with a decrease in ODI (*p* = 0.018) and FWF (*p* < 0.001) for the RS group, but an increase for both metrics in the TMR group.

The main *Learning* effects and the correlations are presented in SI, section 6.1.2.

#### 3.4.2. Post-night structural changes and sleep-related effects in Pre-learning vs. Morning scan (DWI1 vs. DWI3 x RS vs. TMR)

Next, we examined whether the TMR manipulation during the post-training night had a differential impact on overnight microstructural changes, as measured on the next morning (DWI3).

##### 3.4.2.1. Cortical Ribbon

The surface-based statistical analysis on DTI parameters for the total sample disclosed a *Day* (D1 DWI_1_ vs. D2 DWI_3_) effect (CWP < 0.05). Specifically, decreased MD was observed bilaterally in the superior and inferior parietal gyri, the precuneus, the lateral occipital, supramarginal and superior frontal gyri, the cuneus, the isthmus cingulate, precentral, fusiform, postcentral, and caudal anterior cingulate gyri. Also, small clusters emerged in the left posterior cingulate, lingual, pericalcarine and middle temporal gyri and in the right inferior temporal, paracentral, and rostral middle frontal gyri, the right pars opercularis and orbitalis, the banks of the right superior temporal sulcus, the right superior temporal, and lateral orbitofrontal gyri (see Figure 4; for detailed results and cluster size see SI, section 7, Table S4). Overall, MD decreases were still observed on the morning following the experimental night in frontal, parietal and temporal regions, as well as in the occipital cortex but to a lesser extent.

**Figure 4:**
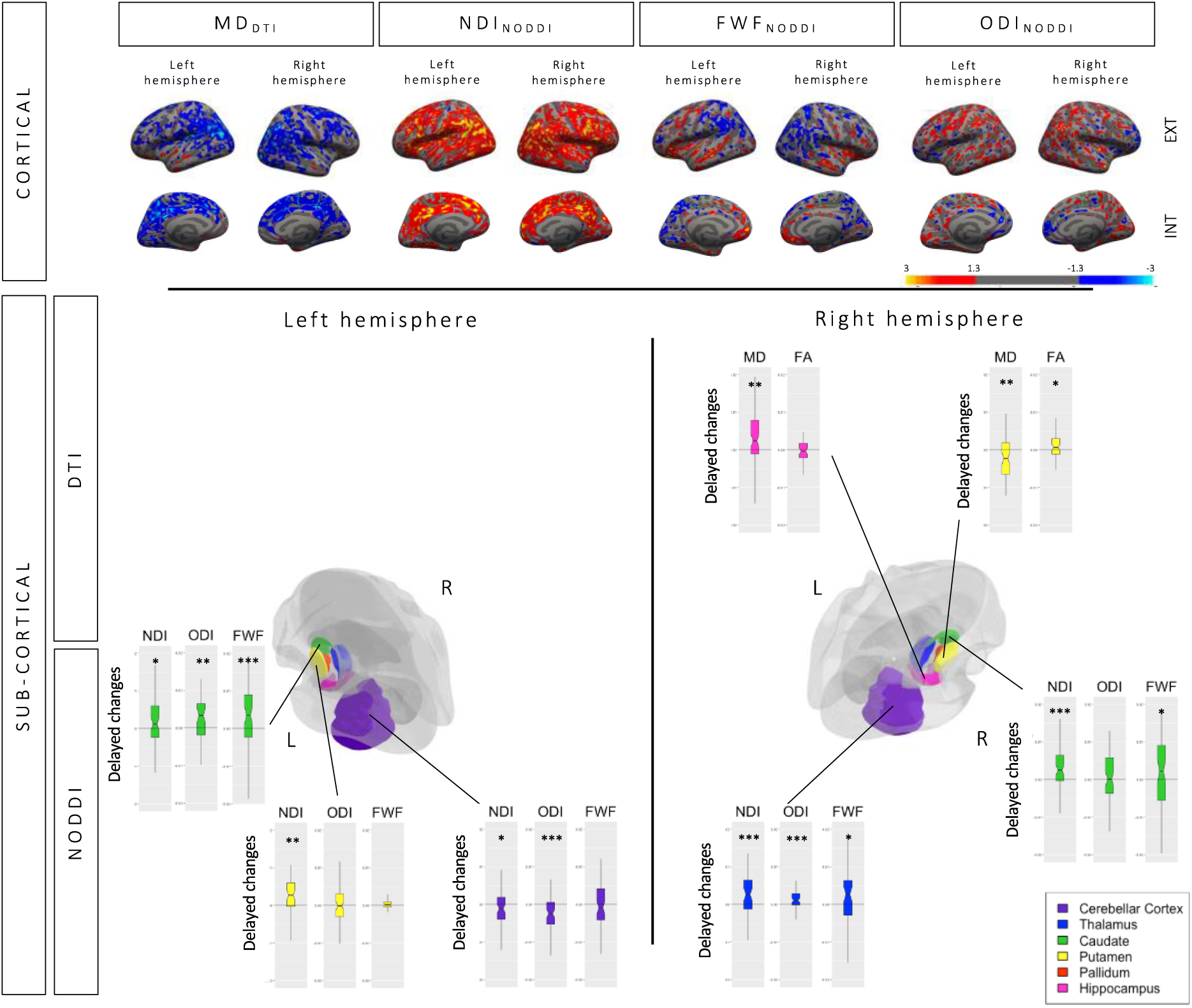
Cortical and subcortical changes following the experimental night (DWI_3_-DWI_1_) On the morning following the experimental night, both groups demonstrated a partial persistence of the changes that emerged following the learning session at the cortical and subcortical level, with reduced magnitude. Cortical *Condition* differences are imaged in Figure 5. Also, the right caudate did show a *Condition*Day* interaction that is detailed in section 3.4.2.2 but not illustrated here. DTI = Diffusion Tensor Imaging, MD = Mean Diffusivity, FA = Fractional Anisotropy, NODDI = Neurite Orientation Dispersion and Density Imaging, NDI = Neurite Density Index, FWF = Free Water Fraction, ODI = Orientation Dispersion Index. Colour-coded cortical images depict Z-scores (uncorrected threshold p < 0.05).

When running the analysis on NODDI parameters, a *Day* (D1 DWI_1_ vs. D2 DWI_3_) effect was evidenced (CWP < 0.05) with NDI increases observed bilaterally in the inferior parietal, superior frontal, superior temporal, precentral, lateral occipital, isthmus cingulate, supramarginal, superior parietal, caudal middle frontal, and the posterior cingulate gyri, the precuneus, the pars opercularis, the insula, the rostral middle frontal gyrus, the banks of the superior temporal sulcus, the pars triangularis, the postcentral gyrus, the cuneus and the pericalcarine gyrus. Other smaller but significant clusters were found in the right caudal anterior cingulate, medial orbitofrontal, and paracentral gyri, the pars orbitalis, and the transverse temporal gyrus as well as in the left inferior temporal, lingual, fusiform, rostral anterior cingulate, middle temporal, and para hippocampal gyri. Interestingly, FWF increases were observed bilaterally in the lateral-and medial orbitofrontal, the middle-and superior temporal gyri and the insula. Also, the right inferior temporal gyrus and the left pars triangularis, superior frontal gyrus, the banks of the left superior temporal sulcus and the parahippocampal gyrus showed similar FWF increase, suggesting an increase in free water inside of the voxels. However, bilateral superior parietal gyrus, the right cuneus, the left lateral occipital, postcentral, lingual, and paracentral gyri, the left precuneus and inferior parietal gyri showed FWF decrease after the experimental night. Lastly, small clusters revealed ODI changes (both increase and decrease depending on the area) in the right superior temporal, inferior parietal, and lingual gyri, the insula, and the para hippocampal gyrus. Also, bilateral changes were observed in the fusiform, lateral orbitofrontal, superior frontal, lateral occipital, inferior temporal, supramarginal, superior parietal, caudal middle frontal, rostral middle frontal, and precentral gyri. Additionally, small clusters were found on the left side only in the precuneus, the rostral anterior cingulate gyrus, the pars triangularis and the postcentral gyrus. To sum up, increased NDI persisted on the next morning even though to a lesser extent. Also, FWF and ODI showed both decreases and increases, suggesting tissue and fiber orientation reorganized during the night as compared to baseline.

Additionally, *Day* effects differed between *Conditions* (RS vs. TMR) in a rostral middle frontal gyrus cluster. The RS group exhibited a strong decrease in MD while the TMR group displayed a trend to increase MD in that area. Regarding FWF, both groups showed an opposite pattern as well, with a light decrease in free water after the night for the RS group but a trend to increase in the TMR group (see Figure 5; for detailed results and cluster size see SI, section 7, Table S5).

**Figure 5:**
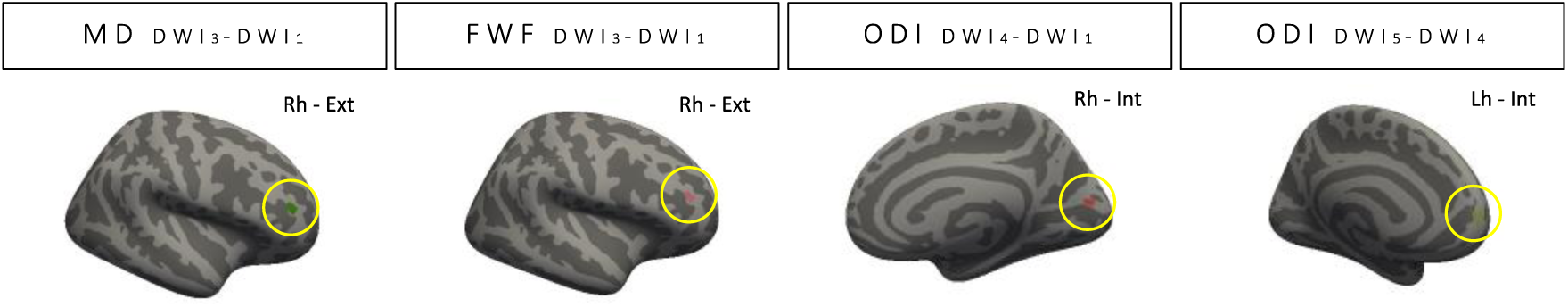
Cortical clusters exhibiting between-group differences. In the right rostral middle frontal gyrus, a region within the prefrontal cortex, we observed on the morning following TMR manipulation (DWI_3_-DWI_1_) a notable decrease in MD and FWF for the RS group, whereas the TMR group exhibited a trend towards an increase in these measures. Additionally, changes in ODI were detected in the right cuneus, part of the occipital visual cortex 3 days later (DWI_4_-DWI_1_). Here, the RS group exhibited a significant decrease, while the TMR group demonstrated a trend for an increase. Furthermore, at the end of the relearning session (DWI_5_-DWI_4_), differences in ODI were also apparent in the superior frontal area, with the RS group showing a trend towards an increase and the TMR group exhibiting a significant decrease.

Finally, there was no significant correlation between clusters exhibiting microstructural changes after the experimental night and the duration of the different sleep stages.

##### 3.4.2.2. Subcortical ROIs

The MANOVAs computed on DTI parameters with within-subject factor *Day* (D1 DWI_1_ vs. D2 DWI_3_) and between-subject factor *Condition* (RS vs. TMR) revealed a significant main effect of *Day* in the right putamen (*p_corr_*= 0.006) and the right hippocampus (*p_corr_* = 0.006; all other *p_corr_*s > 0.017). The post-hoc analysis revealed MD decrease (*p* = 0.008) and FA increase (*p* = 0.020) in the right putamen while the right hippocampus exhibited increased MD (*p* = 0.002; see Figure 4). Also, a significant *Day*\**Condition* interaction emerged in the right caudate (*p_corr_* = 0.002; all other *p_corr_*s > 0.014), with a decrease in MD (*p* < 0.001) and increase in FA (*p* = 0.032) for the RS group as compared to the TMR group that showed the opposite pattern after the experimental night. No main effects of *Condition* emerged (all *p_corr_*s > 0.012).

Likewise, the MANOVAs computed on NODDI parameters disclosed main effects of *Day* in the left cerebellar cortex (*p_corr_* = 0.003), the left putamen (*p_corr_* = 0.005), the left (*p_corr_* < 0.001) and right caudate (*p_corr_* = 0.002), and the right thalamus (*p_corr_* < 0.001). While the cerebellar cortex showed decreased NDI (*p* = 0.018) and decreased ODI (*p* < 0.001), all the other regions showed an increase in NDI (all *p*s < 0.042). Additionally, the left caudate and right thalamus displayed an increase in ODI (all *p*s < 0.007) and increased FWF was observed in bilateral caudate and the right thalamus (all *p*s < 0.031). Significant main *Condition* effects emerged in the right pallidum (*p_corr_* < 0.001) revealing a lower ODI (*p* < 0.001) and FWF (*p* = 0.004) in the RS group compared to the TMR group. Also, a significant *Day*\**Condition* interaction effect in the right caudate (*p_corr_* = 0.002) showed decreased FWF over the night for the RS group but an increase was observed for the TMR group (*p* = 0.001).

Changes in NDI in the right thalamus positively correlated with NREM3 duration (r = 0.379; *p* = 0.003), while changes in FA in the right putamen correlated with REM duration (r = 0.319; *p* = 0.015). Other ROIs exhibiting significant changes after the experimental night did not significantly correlate with the duration of the different sleep stages (all *p*s > 0.076).

#### 3.4.3. Delayed structural changes and sleep-related effects in Pre-learning vs. Pre-relearning (DWI1 vs. DWI4 x RS vs. TMR)

We then investigated for delayed (at Day 5) sleep TMR-related microstructural changes by comparing pre-training DWI at Day5 (DWI4) vs. at Day 1 (DWI1).

##### 3.4.3.1. Cortical Ribbon

The surface-based statistical analysis for the whole sample revealed a *Day* (D1 DWI_1_ vs. D5 DWI_4_) effect (CWP < 0.05) with decreased MD in small clusters located in the left lateral occipital, caudal anterior cingulate and rostral anterior cingulate gyri and in the right fusiform and precentral gyri. Interestingly, a larger cluster located in the inferior temporal gyrus revealed increased MD (see Figure 6; for detailed results and cluster size see SI, section 7, Table S6), suggesting a reorganization of the tissue density by Day 5 as initially, only MD decreases were observed.

**Figure 6:**
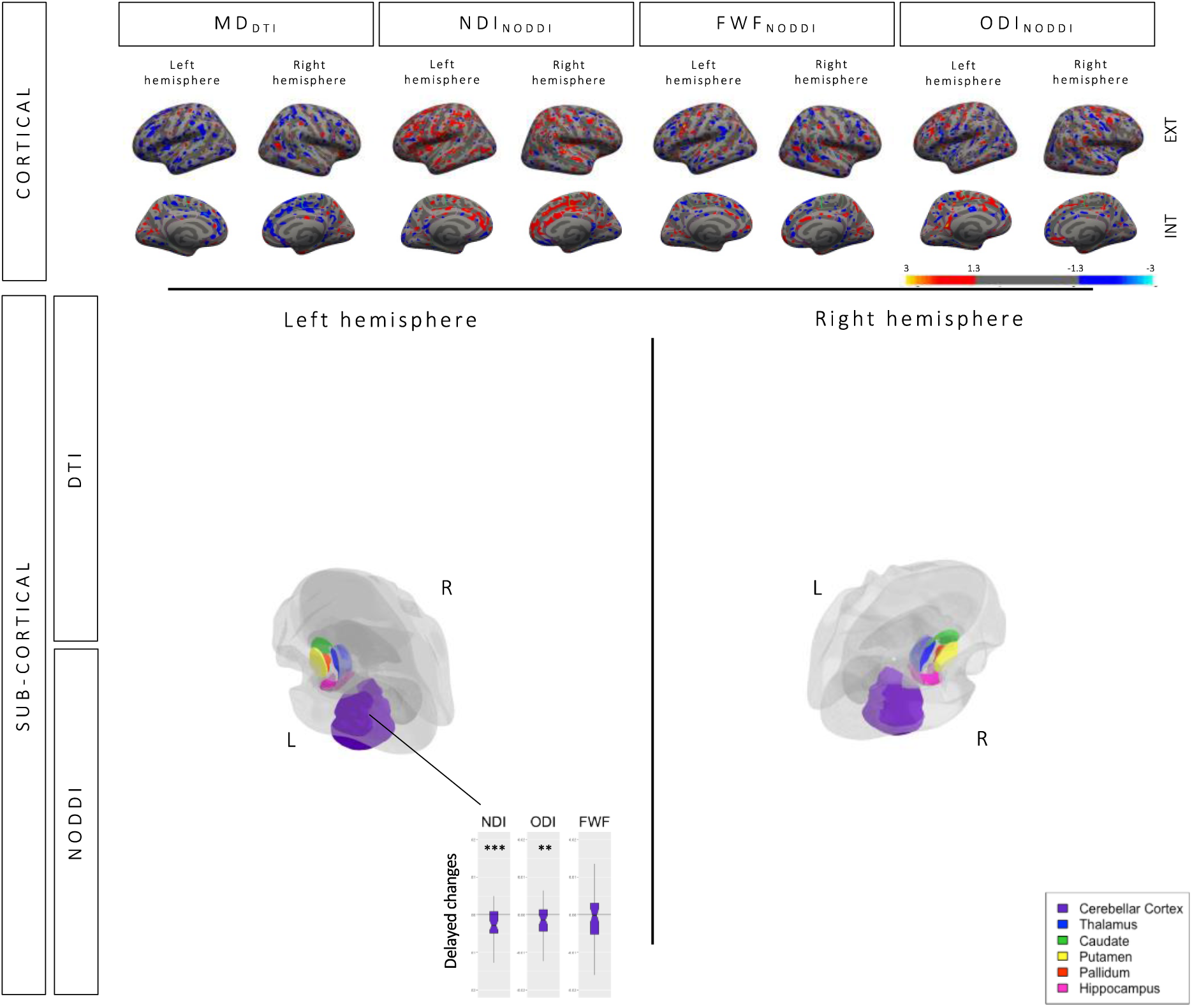
Delayed changes in the cortex and ROIs (DWI_4_-DWI_1_) Limited persistence of learning-related changes at the cortical level and within predefined ROIs across the entire sample when comparing the first delayed scan (DWI_4_), conducted three days after learning, and the baseline scan. Cortical *Condition* differences are imaged in Figure 5. No *Day*Condition* interaction emerged in our ROIs. DTI = Diffusion Tensor Imaging, MD = Mean Diffusivity, FA = Fractional Anisotropy, NODDI = Neurite Orientation Dispersion and Density Imaging, NDI = Neurite Density Index, FWF = Free Water Fraction, ODI = Orientation Dispersion Index. Colour-coded cortical images depict Z-scores representation (uncorrected threshold p < 0.05).

When conducting the same analysis on NODDI metrics, a *Day* (D1 DWI_1_ vs. D5 DWI_4_) effect (CWP < 0.05) revealed decreased NDI in the right isthmus cingulate gyrus only, while the bilateral supramarginal and superior temporal gyri, and the insula as well as the right paracentral and superior frontal gyri and the left postcentral, precentral, and superior parietal gyri showed increased NDI. Also, FWF showed mixed direction changes with decreased FWF in bilateral lateral occipital gyrus, the left superior temporal gyrus, the left insula, the right precentral gyrus. On the other hand, the left caudal middle frontal gyrus, the right insula, and the right superior frontal gyrus showed FWF increase. Lastly, ODI showed changes going in both directions in the bilateral inferior parietal gyrus, the right lateral orbitofrontal and supramarginal gyri and in the left isthmus cingulate, lateral occipital gyri, the left pars opercularis and the left posterior cingulate and paracentral gyri. Altogether, increased NDI persisted in a few clusters, and FWF/ODI both decreased and increased depending on the area, suggesting tissue and fiber orientation reorganization over time.

When comparing *Day* effects between *Conditions* (RS vs. TMR), there was a different ODI pattern between groups in the right cuneus, - with a strong decrease in the RS group but a trend to increase in the TMR group (see Figure 5; for detailed results and cluster size see SI, section 7, Table S7).

Microstructural changes that persisted on Day 5 did not significantly correlate with the duration of the different sleep stages during the experimental night.

##### 3.4.3.2. Subcortical ROIs

The MANOVAs computed on DTI parameters with within-subject factor *Day* (D1 DWI_1_ vs. D5 DWI_4_) and between-subject factor *Condition* (RS vs. TMR) did not reveal any significant main effect of *Day* (all *p_corr_* > 0.036), *Condition* (all *p_corr_* > 0.017), or *Day*Condition* interaction (all *p_corr_* > 0.180) for any of our ROIs (see Figure 6).

Likewise, the MANOVAs computed on NODDI parameters did not disclose any *Day*Condition* interaction effect (all *p_corr_* > 0.175) but a main effect of *Day* in the left cerebellar cortex (*p_corr_* < 0.001) with a significant decrease in NDI (*p* < 0.001) and ODI (*p* = 0.003). Also, a main effect of *Condition* emerged in the right pallidum (*p_corr_* < 0.001) with a lower ODI (*p* < 0.001) and FWF (*p* = 0.005) in the RS group compared to the TMR group. These results suggest a partial persistence at the beginning of Day 5 of the changes already observed right after the experimental night in subcortical ROIs.

ODI changes in the left cerebellum positively correlated with REM sleep duration (r = 0.275; *p* = 0.035). All other correlations were non-significant (all *p*s > 0.139).

#### 3.4.4. Motor retraining-related structural changes and sleep-related effects (Day 5; DWI4 vs. DWI5 X RS vs. TMR)

Finally, we tested for repeated practice-related effects on microstructural changes comparing pre- (DWI4) and post-relearning session (DWI5) acquisitions, possibly modulated by the TMR manipulation conducted during the night after Day1.

##### 3.4.4.1. Cortical Ribbon

Correlations and *Learning* effects common to both *Conditions* are reported in SI, section 6.2.1. *Relearning* effects revealed to be different between *Conditions* (RS vs. TMR) with ODI differences in the superior frontal area with a trend to increase for the RS group but a significant decrease in the TMR group (see Figure 5; for detailed results and cluster size see SI, section 7, Table S9).

##### 3.4.4.2. Subcortical ROIs

The MANOVA on DTI parameters with within-subject factor *ReLearning* (Pre DWI_4_ vs. Post DWI_5_) and between factor *Condition* (RS vs. TMR), revealed no significant main effect of *Condition* (all *p_corr_* > 0.050) or *ReLearning*Condition* interaction (all *p_corr_* > 0.051) effects, suggesting that the post-learning TMR manipulation did not modulate the relearning of previously studied material.

The MANOVA on NODDI parameters disclosed a main *Condition* effect in the right pallidum (*p_corr_* = 0.002; all other *p_corr_*s > 0.077), but no *ReLearning*Condition* interaction (all *p_corr_*s > 0.185) effects.

The main *Learning* effects and correlations are provided in SI, section 6.2.2.

#### 3.4.5. Post-hoc analyses on the hippocampus and caudate nucleus

Separate MANOVAs with within-subject factors *Session* (Learning D1 vs. Relearning D5) and *Moment* (Pre vs. Post) were conducted on the bilateral hippocampus and caudate nucleus for both groups combined, allowing for a direct comparison of diffusion changes in these regions during the learning (DWI_2_-DWI_1_) and relearning sessions (DWI_5_-DWI_4_).

For DTI parameters, a significant *Session* effect was observed in the right caudate (*p_corr_* < 0.001), driven by a significant increase in FA between the learning and relearning sessions (*p* = 0.010). No significant *Session* effect was found in the left caudate (*p_corr_* = 0.030) or in either part of the hippocampus (both *p_corr_*s > 0.506). Significant *Moment* effects emerged in the left caudate (*p_corr_* = 0.004) and in both parts of the hippocampus (both *p_corr_*s < 0.001), but not in the right caudate (*p_corr_* = 0.033). These changes were driven by decreases in MD in both parts of the hippocampus (*p*s < 0.001). No significant interaction effects were found (all *p_corr_* > 0.140).

For NODDI parameters, a similar pattern was observed. A significant *Session* effect was present in both parts of the caudate (both *p_corr_*s < 0.002), driven by changes in ODI in the right caudate (*p* < 0.001) and changes in FWF in the left caudate (*p* < 0.001). However, no significant *Session* effects were detected in either part of the hippocampus (both *p_corr_*s > 0.377). Significant *Moment* effects emerged in the left caudate (*p_corr_* < 0.001) and in both parts of the hippocampus (both *p_corr_*s < 0.002), but not in the right caudate (*p_corr_*= 0.045). These changes were driven by increases in ODI in these 3 subregions (all *p*s < 0.023), increases in NDI in the left caudate (*p* = 0.004) and left hippocampus (*p* = 0.028), and decreases in FWF in the left (*p* = 0.004) and right hippocampus (*p* = 0.005). No significant interaction effects were observed (all *p_corr_* > 0.140).

## 4. Discussion

In the present study, we investigated rapid training-related brain reorganization and to what extent TMR during post-learning sleep is associated with both immediate (on the next morning; Day 2) and more delayed (at Day 5) microstructural changes in the brain. Besides conventional DTI, we used NODDI for a more specific approach regarding the components involved in microstructural remodeling. We predicted that the TMR group would show accelerated structural reorganization, which would be reflected in behavioral performance improvements in comparison to the RS group. Therefore, we also expected noticeable differences in diffusion metrics after the experimental night, that may or may not persist after three days. Results showed a rapid learning curve on day 1 followed by a stabilization on day 2 and an improvement in performance on day 5 without further practice, but not modulated by the TMR manipulation. Rapid (re)training-related microstructural changes were observed after 40 or 60 min of motor practice, and these changes partially persisted and evolved over time. Besides widespread rapid training-related changes in brain microstructure immediately after task practice, the post-learning session TMR manipulation during sleep after Day 1 resulted in differential changes in a limited subset of clusters.

At the behavioral level, participants exhibited a significant and similar improvement in GPI over time, independently of their experimental condition (RS or TMR group), suggesting that motor sequence cueing during post-learning session sleep did not lead to additional performance gains. After a significant improvement during the learning session (Day 1), performance stabilized on the morning retest (Day 2), suggesting that the first night after initial training did not lead to significant offline improvements. Nevertheless, participants demonstrated significant offline improvements between the morning retest on Day 2 and the beginning of the retraining session (Day 5). On day 5, participants exhibited significant online gains during retraining. Also, significant drops in performance were observed on pseudo-random blocks, demonstrating sequence-dependent learning rather than mere sequence-independent (or motor) learning. Hence, while all our groups displayed significant online performance gains and offline consolidation, the effect of auditory cueing during sleep (TMR) was not evidenced at the behavioral level.

Previous studies have shown that auditory cues presented during both N2 and SWS stages can enhance memory, leading to noticeable improvements in accuracy and/or reaction times the following day [39], [40], [41], [42]. In our study, auditory stimulation during N2 and N3 sleep stages did not yield significant behavioral effects on the next day and 3 days later. However, it cannot be excluded that the TMR manipulation in stages N2 and N3 of the first two sleep cycles enhanced memory processing during the early part of the night, eventually reaching a saturation point that was attained by the regular sleep (RS) group but after a longer sleep duration. This aligns with the findings of a study [40] which observed TMR benefits after 2 hours of cueing within a 3-hour nocturnal sleep period, but not after 3 hours without cueing. Interestingly, the TMR-related gains in this latter study were comparable to those resulting from a full night of regular nocturnal sleep (8 hours). Similar results were obtained in the spatial domain with equivalent performance gains after 40min of odor cueing or 90min of regular sleep [83]. Also, the timing of performance assessment following TMR appears to be a crucial factor influencing the outcomes. A longitudinal study [43] revealed that the benefits of TMR may not be immediate, instead emerging only gradually over time. In this study, the authors found that while TMR had no immediate effect on motor performance, its positive impacts became evident 10 days after the experiment, then disappeared at six weeks [43]. Considering that our performance assessments were conducted on the morning following a full night’s sleep with TMR manipulation and again three days later, it is possible that rapid gains developed overnight in the TMR group, which might have been subsequently matched by the regular sleep group. Also, we cannot rule out potential delayed performance improvements in our participants, which may have become evident over a more extended period beyond our assessment timeline.

Whereas many studies demonstrated a generally positive (although moderate) effect of TMR on skill learning [37], [38], [39], [40], [41], [42] (see [36] for a review; also see SI, section 7, Table S10 for a summary), the efficacy of TMR appears to be influenced by several factors. For instance, research suggests that TMR might preferentially consolidate weaker memory traces. This is supported by findings showing that SWS cueing primarily benefits the non-dominant hand, which typically shows weaker performance, as compared to the dominant hand where TMR-related improvements are not as pronounced [84]. Hence, it is possible that the already strong memory trace formed after 1 hour of SRTT practice (30 blocks) in our study left not enough room for further significant gains.

Considering different sensory modalities and sleep stages, olfactory cues presented during SWS did not result in improved performance [44], [85], despite the direct pathway for odor information from the olfactory bulb to critical memory-related areas such as the hippocampus [86]. In contrast, olfactory cueing during N2 sleep was found to improve performance in other studies [37], [38]. This TMR-related improvement was linked to increased spindle frequency activity in the parietal brain regions [37]. Moreover, TMR attempts using tactile stimuli have not been successful [87], [88]. Such attempts appeared to disrupt the coupling of SOs and sleep spindles [87], a process believed to be fundamental for motor memory consolidation [89], [90], [91]. Indeed, strong evidence suggests a correlation between more effective sleep-related memory enhancements and improved SOs/spindles coupling in motor memory [92], [93]. Effective memory recall of the content associated with cues may also significantly depend on the precise timing of their delivery. To be effective, this timing should align with the natural rhythms of sleep spindle activity [94], [95] and SOs during sleep [89], while avoiding excessive interference with these processes. For instance, cue effectiveness reduces when cues are presented too close to each other (less than ∼1500msec), leading to the suppression of spindle activity [96], [97]. Antony et al. [94] further demonstrated this effect by tracking sleep spindles in real-time and manipulating the interval between a spindle and a subsequent sensory cue. Most effective memory enhancement was observed when cues were delivered 3 to 6 seconds after the end of a spindle [94]. Concerning SOs, cueing during the upstate of an SO showed significant positive effects [98], [99]. However, mixed results were obtained when it comes to cueing during the downstate of SOs – some studies reported benefits [100], while others produced inconclusive findings [101]. Noticeably, the synchronization of cues with SO-spindle coupling periods seems particularly beneficial [99], supporting the notion that these coupling periods are critical for memory reactivation and consolidation. In our study, while spindle density remained stable across both nights and groups, we observed a specific reduction in SO density during the first two sleep cycles of the experimental night in the TMR group. This suggests that the application of 6-second auditory cues in our research, needed to initiate the replay of the learned sequence, may not have been ideally timed to align with the spontaneous neural oscillations, potentially leading to a disruption rather than facilitation of these oscillatory patterns, and possibly resulting in the absence of additional behavioral improvements.

At the microstructural level, decreased MD within extensive regions of the occipito-parietal, temporal, and frontal cortices were observed following the learning session. This reduction in diffusivity overlapped with increased NDI, indicating an increase in neurite density within these cortical areas as a result of task practice. Furthermore, these changes were accompanied by a reduction in FWF, which confirm increased tissue fraction. At the subcortical level, bilaterally decreased MD was found in the putamen, hippocampus, and cerebellar cortex, along with the left thalamus. Notably, NODDI highlighted that MD changes in the putamen were underpinned by increased NDI, suggesting an increase in neurite density in this subsection of the striatum. The observed increase in neurite density within the putamen aligns with the incremental activation of the striato-cortical network in response to task practice [102], highlighting the role of this network in the learning process. In contrast, modulations in the other subcortical areas were associated with decreased FWF but no NDI changes, implying that training may mainly induce glial reorganization rather than neurite creation in these regions, as tissue fraction increased (i.e., decreased FWF) more relatively to neurites (stable NDI). Interestingly, before TMR manipulation was applied, the right caudate already exhibited distinct patterns between the two conditions. The RS group exhibited reductions in MD, FWF and ODI, whereas the TMR group demonstrated changes in the opposite direction. Given that these differences emerged at the pre-experimental manipulation stage of our procedure (i.e., during the learning phase), they might possibly be due to the distinct strategies adopted by each group. Indeed, separate analyses conducted on speed and accuracy (see SI, section 3) showed that the RS group appeared to focus more on speed, whereas the TMR group prioritized accuracy.

On the day following the experimental night, changes persisted within the cortical regions for both groups, even if to a lesser extent. In the subcortical regions, modulations were still observed in the putamen, hippocampus, cerebellar cortex, and the left thalamus. The interaction observed in the right caudate over the learning session persisted. Moreover, new modifications (main effects) became apparent bilaterally in the caudate nucleus, a part of the striatum, for both groups, which may indicate enhanced post-sleep integration within the cortico-striatal network, as previously reported at the functional level [103]. These changes included both increased NDI, ODI and FWF, suggesting an optimization of the internal microstructure of the caudate nucleus, as neurite density and fiber dispersion increased while tissue fraction decreased. Additionally, a differential effect of the experimental manipulation was observed in the rostral middle frontal gyrus. The RS group exhibited a marked decrease in MD/FWF, reflecting densification of brain tissue, whereas the TMR group showed a trend towards an increase in MD/FWF in the same region, indicating an increase in water fraction.

Increased functional TMR-related activity was observed in the DLPFC, along with other cortical and cerebellar regions, in response to SWS cueing. This heightened activation correlated with the duration of REM sleep. Additionally, increased learning-related activity in the caudate nucleus and hippocampus was associated with the time spent in SWS [42]. It has been proposed that SWS promotes consolidation in subcortical regions, such as the striatum and hippocampus, which are involved in sequence learning. In contrast, REM sleep might be implicated in consolidating motor-related learning, involving cortical and cerebellar networks [42]. Our findings partially support previous research, showing correlations between diffusion metrics in subcortical regions (right thalamus) and SWS duration. We also observed correlations between diffusion metrics in the right putamen and REM duration on Day 2. Furthermore, a correlation between ODI in the left cerebellar cortex on Day 5 and REM aligns with earlier findings, although cortical correlations were not observed.

The increase in REM sleep duration that we observed on the experimental night is in line with prior reports [104], [105], [106]. While increased REM has been hypothesized to represent a state of motor memory stabilization and integration at the behavioral level [104], it might also play a role in stabilizing and integrating memory traces within motor sequence-related brain structures.

Three days after the experimental night, decreased MD persisted in a few small clusters and a significantly increased MD appeared within a larger cluster in the inferior temporal gyrus. Increased NDI was sustained in several clusters, yet a decrease was noted in the right isthmus of the cingulate gyrus. FWF and ODI exhibited mixed directional changes. Taken together, these observations indicate possible neural reorganization over this period. When comparing both groups, the RS group exhibited a pronounced decrease in ODI in the right cuneus, while the TMR group exhibited a trend toward an increase, suggesting different neural adaptation patterns. At the subcortical level, a reduction in NDI/ODI was found in the left cerebellar cortex. A noticeable between-group difference emerged in the right pallidum, with the RS group displaying lower ODI and FWF compared to the TMR group. This may suggest a differential reorganization, by Day 5, in motor-related networks.

It is worth mentioning that it is possible the microstructural differences observed after TMR manipulation may also stem from different learning strategies adopted by the RS and TMR groups, rather than TMR itself, as already mentioned earlier. For example, in a previous dMRI study, participants demonstrated MD changes within the temporal, motor, and cerebellar regions — including the premotor cortex, supplementary motor area, and middle temporal gyrus — after an accuracy-focused session. In contrast, structural changes were detected in the occipitotemporal areas, including the lingual gyrus and inferior temporal gyrus, as well as in frontal areas, specifically the inferior frontal gyrus, when participants were instructed to prioritize speed [6].

Lastly, the relearning session induced changes similar to those of the initial learning session but less pronounced, suggesting that cortical networks undergo reorganization and optimization during the 3-day interval separating both practice sessions. Subcortical changes were found in the caudate nucleus, putamen, thalamus, and cerebellar cortex, which differed from the initial learning session that similarly affected the putamen, thalamus, and the cerebellar cortex, but also the hippocampus. This pattern supports findings showing that the initial encoding stages heavily involve the cortico-hippocampal networks, whose activity decreases over the first learning episode. Meanwhile, the sensorimotor cortex, along with the cortico-striatal and cortico-cerebellar networks, exhibits increased activation that aligns with task practice [102]. Indeed, the basal ganglia and more specifically the striatum is thought to be involved in the automatization of motor skills [94]. Additionally, sleep is thought to enhance the integration within the cortico-striatal network [103], which could explain the more pronounced involvement of the caudate nucleus during the relearning session as opposed to the initial learning phase. Finally, we identified group differences that exhibited opposing patterns in ODI within the superior frontal area, indicating specific patterns of neural reorganization.

This study also contains certain limitations. We were unable to include a control task, which would have provided a clearer distinction between motor-specific and sequence-specific remodeling. However, the inclusion of random blocks within the training sessions allowed us to differentiate, at least behaviorally, between improvements attributable to motor-specific learning and those related to sequence-specific learning. Despite this, the imaging of dMRI data presents challenges due to its cumulative and long-lasting nature as compared to fMRI.

As a result, while the random blocks confirmed efficient sequential learning among our participants, the observed dMRI modifications likely reflect a combination of both motor-and sequence-related learning effects. Also, the SRTT is supposed to capture implicit motor learning. However, participants may become aware, at least partially, of the repetitive pattern over time [46], [107], [108]. In our study, associating sounds with specific positions or keypresses made it easier for participants to identify the sequence, thereby contributing to the enhancement of their explicit knowledge. Additionally, TMR has been shown to enhance explicit recall of cued sequences compared to non-cued ones [41], an effect that has been replicated but found to be gender-specific, favoring men mostly [44]. Although all participants in our study reported recognizing a repetitive pattern by the end of the protocol (see SI, section 1 for detailed analysis), the precise timing they realized its presence (i.e., immediately after learning, the following morning, or during relearning) remains uncertain and could have differentially influenced consolidation processes (for review, see [15]). However, since we chose not to inform participants about the sequence’s presence, collecting explicit knowledge measures during the protocol to track its evolution along performance was not possible.

In the present study, we offer evidence for rapid motor training-related changes in the brain microstructure associated with an initial learning phase and a subsequent relearning session. Initial acquisition engaged a widespread cortical network and motor-related subcortical regions, including the hippocampus. In contrast, the relearning session recruited similar cortical networks, even if to a reduced extent, along with the caudate nucleus. Using NODDI, we provide more specific information concerning the mechanisms behind these changes. We observed that MD alterations paralleled increases in NDI within the putamen, suggesting augmented neurite density. Meanwhile, other subcortical areas exhibited decreased FWF over the learning session, suggesting different processes, such as glial reorganization, beyond simple neurite proliferation. Differences between groups were evidenced in the DLPFC the morning after TMR manipulation, with both groups displaying opposite MD/FWF evolution, indicating contrasting patterns in tissue density/fraction between the groups. Three days later, differences in ODI became apparent in the right cuneus, and a divergence in ODI/FWF was observed in the right pallidum. These findings indicate distinct neural reorganization processes, potentially resulting from the TMR manipulation, or alternatively from the specific learning strategies employed. The rapid motor practice-related changes persisted in the longer term, even if with reduced magnitude. This suggests an extensive reorganization within motor-related neural networks.

## Acknowledgements

W.S. is supported by the Fonds de la Recherche Scientifique (FRS-FNRS., Aspirant Research Fellowship). The study and postdoctoral fellows A.L. and M.G. were supported by the FNRS and the Fonds Wetenschappelijk Onderzoek – Vlaanderen (FWO) under the Excellence of Science (EOS) Project (MEMODYN, No. 30446199 to P.P. and H.Z.). The authors thank Dimitri Voisin and Louis-Jacques Etaix for their help in data acquisition.

## Data availability statement

The data underlying this article are available upon reasonable request to the corresponding author.

## Bibliography

[1] M. P. Walker, “A refined model of sleep and the time course of memory formation,” Behav. Brain Sci., vol. 28, no. 1, pp. 51–64, Feb. 2005, doi: 10.1017/S0140525X05000026.

[2] Y. Chang, “Reorganization and plastic changes of the human brain associated with skill learning and expertise,” Front. Hum. Neurosci., vol. 8, 2014, doi: 10.3389/fnhum.2014.00035.

[3] B. Kolb, R. Gibb, and T. E. Robinson, “Brain Plasticity and Behavior,” Curr. Dir. Psychol. Sci., vol. 12, no. 1, pp. 1–5, Feb. 2003, doi: 10.1111/1467-8721.01210.

[4] F. Jacobacci et al., “Rapid hippocampal plasticity supports motor sequence learning,” Proc. Natl. Acad. Sci. U. S. A., vol. 117, no. 38, pp. 23898–23903, Sep. 2020, doi: 10.1073/PNAS.2009576117/-/DCSUPPLEMENTAL.

[5] W. Stee, A. Legouhy, M. Guerreri, T. Villemonteix, H. Zhang, and P. Peigneux, “Microstructural dynamics of motor learning and sleep-dependent consolidation: a diffusion imaging study,” iScience, p. 108426, Nov. 2023, doi: 10.1016/J.ISCI.2023.108426.

[6] I. Tavor, R. Botvinik-Nezer, M. Bernstein-Eliav, G. Tsarfaty, and Y. Assaf, “Short-term plasticity following motor sequence learning revealed by diffusion magnetic resonance imaging,” Hum. Brain Mapp., vol. 41, no. 2, pp. 442–452, Feb. 2020, doi: 10.1002/hbm.24814.

[7] H. Zhang, T. Schneider, C. A. Wheeler-Kingshott, and D. C. Alexander, “NODDI: Practical in vivo neurite orientation dispersion and density imaging of the human brain,” NeuroImage, vol. 61, no. 4, pp. 1000–1016, Jul. 2012, doi: 10.1016/j.neuroimage.2012.03.072.

[8] B. Rasch and J. Born, “About sleep’s role in memory,” Physiol. Rev., vol. 93, no. 2, pp. 681– 766, 2013, doi: 10.1152/physrev.00032.2012.

[9] K. Debas et al., “Brain plasticity related to the consolidation of motor sequence learning and motor adaptation,” Proc. Natl. Acad. Sci. U. S. A., vol. 107, no. 41, pp. 17839–17844, Oct. 2010, doi: 10.1073/PNAS.1013176107.

[10] S. M. Fogel et al., “fMRI and sleep correlates of the age-related impairment in motor memory consolidation,” Hum. Brain Mapp., vol. 35, no. 8, pp. 3625–3645, 2014, doi: 10.1002/HBM.22426.

[11] S. Diekelmann, I. Wilhelm, and J. Born, “The whats and whens of sleep-dependent memory consolidation,” Sleep Med. Rev., vol. 13, no. 5, pp. 309–321, Oct. 2009, doi: 10.1016/J.SMRV.2008.08.002.

[12] G. Albouy, P. Ruby, C. Phillips, A. Luxen, P. Peigneux, and P. Maquet, “Implicit oculomotor sequence learning in humans: Time course of offline processing,” Brain Res., vol. 1090, no. 1, pp. 163–171, May 2006, doi: 10.1016/j.brainres.2006.03.076.

[13] G. Albouy et al., “Sleep stabilizes visuomotor adaptation memory: a functional magnetic resonance imaging study,” J. Sleep Res., vol. 22, no. 2, pp. 144–154, 2013, doi: 10.1111/J.1365-2869.2012.01059.X.

[14] S. Song, J. H. Howard, and D. V. Howard, “Implicit probabilistic sequence learning is independent of explicit awareness,” Learn. Mem., vol. 14, no. 3, p. 167, Mar. 2007, doi: 10.1101/LM.437407.

[15] B. R. King, K. Hoedlmoser, F. Hirschauer, N. Dolfen, and G. Albouy, “Sleeping on the motor engram: The multifaceted nature of sleep-related motor memory consolidation,” Neurosci. Biobehav. Rev., vol. 80, pp. 1–22, Sep. 2017, doi: 10.1016/J.NEUBIOREV.2017.04.026.

[16] J. Born, B. Rasch, and S. Gais, “Sleep to remember,” Neuroscientist, vol. 12, no. 5, pp. 410–424, Oct. 2006, doi: 10.1177/1073858406292647.

[17] M. A. Wilson and B. L. McNaughton, “Reactivation of hippocampal ensemble memories during sleep,” Science, vol. 265, no. 5172, pp. 676–679, 1994, doi: 10.1126/SCIENCE.8036517.

[18] M. G. Frank and H. C. Heller, “The Function(s) of Sleep,” Handb. Exp. Pharmacol., vol. 253, pp. 3–34, 2019, doi: 10.1007/164_2018_140.

[19] D. S. Ramanathan, T. Gulati, and K. Ganguly, “Sleep-Dependent Reactivation of Ensembles in Motor Cortex Promotes Skill Consolidation,” PLOS Biol., vol. 13, no. 9, p. e1002263, Sep. 2015, doi: 10.1371/journal.pbio.1002263.

[20] S. Vahdat, S. Fogel, H. Benali, and J. Doyon, “Network-wide reorganization of procedural memory during NREM sleep revealed by fMRI,” eLife, vol. 6, Sep. 2017, doi: 10.7554/ELIFE.24987.

[21] S. Fogel et al., “Reactivation or transformation? Motor memory consolidation associated with cerebral activation time-locked to sleep spindles,” PloS One, vol. 12, no. 4, Apr. 2017, doi: 10.1371/JOURNAL.PONE.0174755.

[22] A. Boutin, B. Pinsard, A. Boré, J. Carrier, S. M. Fogel, and J. Doyon, “Transient synchronization of hippocampo-striato-thalamo-cortical networks during sleep spindle oscillations induces motor memory consolidation,” NeuroImage, vol. 169, pp. 419–430, Apr. 2018, doi: 10.1016/J.NEUROIMAGE.2017.12.066.

[23] D. B. Rubin et al., “Learned Motor Patterns Are Replayed in Human Motor Cortex during Sleep,” J. Neurosci., vol. 42, no. 25, pp. 5007–5020, Jun. 2022, doi: 10.1523/JNEUROSCI.2074-21.2022.

[24] P. Peigneux et al., “Learned material content and acquisition level modulate cerebral reactivation during posttraining rapid-eye-movements sleep,” NeuroImage, vol. 20, no. 1, pp. 125–134, Sep. 2003, doi: 10.1016/S1053-8119(03)00278-7.

[25] P. Maquet et al., “Experience-dependent changes in cerebral activation during human REM sleep,” Nat. Neurosci., vol. 3, no. 8, pp. 831–836, Aug. 2000, doi: 10.1038/77744.

[26] S. Fogel, C. Vien, A. Karni, H. Benali, J. Carrier, and J. Doyon, “Sleep spindles: a physiological marker of age-related changes in gray matter in brain regions supporting motor skill memory consolidation,” Neurobiol. Aging, vol. 49, pp. 154–164, Jan. 2017, doi: 10.1016/J.NEUROBIOLAGING.2016.10.009.

[27] G. Piantoni et al., “Individual differences in white matter diffusion affect sleep oscillations,” J. Neurosci., vol. 33, no. 1, pp. 227–233, Jan. 2013, doi: 10.1523/JNEUROSCI.2030-12.2013.

[28] C. Vien et al., “Thalamo-Cortical White Matter Underlies Motor Memory Consolidation via Modulation of Sleep Spindles in Young and Older Adults,” Neuroscience, vol. 402, pp. 104–115, Mar. 2019, doi: 10.1016/j.neuroscience.2018.12.049.

[29] I. Wilhelm et al., “Sleep slow-wave activity reveals developmental changes in experience-dependent plasticity,” J. Neurosci., vol. 34, no. 37, pp. 12568–12575, Sep. 2014, doi: 10.1523/JNEUROSCI.0962-14.2014.

[30] V. V. Vyazovskiy et al., “Cortical firing and sleep homeostasis,” Neuron, vol. 63, no. 6, p. 865, Sep. 2009, doi: 10.1016/J.NEURON.2009.08.024.

[31] J. M. Saletin, E. van der Helm, and M. P. Walker, “Structural brain correlates of human sleep oscillations,” NeuroImage, vol. 83, pp. 658–668, Dec. 2013, doi: 10.1016/j.neuroimage.2013.06.021.

[32] S. Kurth et al., “Mapping the electrophysiological marker of sleep depth reveals skill maturation in children and adolescents,” NeuroImage, vol. 63, no. 2, pp. 959–965, Nov. 2012, doi: 10.1016/j.neuroimage.2012.03.053.

[33] D. Bendor and M. A. Wilson, “Biasing the content of hippocampal replay during sleep,” Nat. Neurosci. 2012 1510, vol. 15, no. 10, pp. 1439–1444, Sep. 2012, doi: 10.1038/nn.3203.

[34] N. Cellini and A. Capuozzo, “Shaping memory consolidation via targeted memory reactivation during sleep,” Ann. N. Y. Acad. Sci., vol. 1426, no. 1, pp. 52–71, 2018, doi: 10.1111/NYAS.13855.

[35] D. I. Schouten, S. I. R. Pereira, M. Tops, and F. M. Louzada, “State of the art on targeted memory reactivation: Sleep your way to enhanced cognition,” Sleep Med. Rev., vol. 32, pp. 123–131, Apr. 2017, doi: 10.1016/J.SMRV.2016.04.002.

[36] X. Hu, L. Y. Cheng, M. H. Chiu, and K. A. Paller, “Promoting memory consolidation during sleep: A meta-analysis of targeted memory reactivation,” Psychol. Bull., vol. 146, no. 3, pp. 218–244, Mar. 2020, doi: 10.1037/BUL0000223.

[37] S. Laventure, S. Fogel, and O. Lungu, “NREM2 and Sleep Spindles Are Instrumental to the Consolidation of Motor Sequence Memories,” pp. 1–27, 2016, doi: 10.1371/journal.pbio.1002429.

[38] S. Laventure et al., “Beyond spindles: interactions between sleep spindles and boundary frequencies during cued reactivation of motor memory representations,” Sleep, vol. 41, no. 9, pp. 1–14, Sep. 2018, doi: 10.1093/SLEEP/ZSY142.

[39] J. W. Antony, E. W. Gobel, J. K. O’Hare, P. J. Reber, and K. A. Paller, “Cued memory reactivation during sleep influences skill learning,” Nat. Neurosci. 2012 158, vol. 15, no. 8, pp. 1114–1116, Jun. 2012, doi: 10.1038/nn.3152.

[40] M. Schönauer, T. Geisler, and S. Gais, “Strengthening Procedural Memories by Reactivation in Sleep,” J. Cogn. Neurosci., vol. 26, no. 1, pp. 143–153, Jan. 2014, doi: 10.1162/JOCN_A_00471.

[41] J. N. Cousins, W. El-Deredy, L. M. Parkes, N. Hennies, and P. A. Lewis, “Cued memory reactivation during slow-wave sleep promotes explicit knowledge of a motor sequence,” J. Neurosci. Off. J. Soc. Neurosci., vol. 34, no. 48, pp. 15870–15876, Nov. 2014, doi: 10.1523/JNEUROSCI.1011-14.2014.

[42] J. N. Cousins, W. El-Deredy, L. M. Parkes, N. Hennies, and P. A. Lewis, “Cued Reactivation of Motor Learning during Sleep Leads to Overnight Changes in Functional Brain Activity and Connectivity,” PLoS Biol., vol. 14, no. 5, May 2016, doi: 10.1371/journal.pbio.1002451.

[43] M. Rakowska, M. E. A. Abdellahi, P. Bagrowska, M. Navarrete, and P. A. Lewis, “Long term effects of cueing procedural memory reactivation during NREM sleep,” NeuroImage, vol. 244, p. 118573, Dec. 2021, doi: 10.1016/J.NEUROIMAGE.2021.118573.

[44] S. Diekelmann, J. Born, and B. Rasch, “Increasing Explicit Sequence Knowledge by Odor Cueing during Sleep in Men but not Women,” Front. Behav. Neurosci., vol. 10, no. APRIL, Apr. 2016, doi: 10.3389/FNBEH.2016.00074.

[45] K. Jann et al., “Linking Brain Connectivity Across Different Time Scales with Electroencephalogram, Functional Magnetic Resonance Imaging, and Diffusion Tensor Imaging,” Brain Connect., vol. 2, no. 1, pp. 11–20, Feb. 2012, doi: 10.1089/brain.2011.0063.

[46] M. J. Nissen and P. Bullemer, “Attentional requirements of learning: Evidence from performance measures,” Cognit. Psychol., vol. 19, no. 1, pp. 1–32, Jan. 1987, doi: 10.1016/0010-0285(87)90002-8.

[47] S. Titone, J. Samogin, P. Peigneux, S. Swinnen, D. Mantini, and G. Albouy, “Connectivity in Large-Scale Resting-State Brain Networks Is Related to Motor Learning: A High-Density EEG Study,” Brain Sci., vol. 12, no. 5, p. 530, May 2022, doi: 10.3390/BRAINSCI12050530/S1.

[48] C. H. (Janice) Lin et al., “Contextual interference enhances motor learning through increased resting brain connectivity during memory consolidation,” NeuroImage, vol. 181, pp. 1–15, Nov. 2018, doi: 10.1016/J.NEUROIMAGE.2018.06.081.

[49] S. Sami, E. M. Robertson, and R. Chris Miall, “The time course of task-specific memory consolidation effects in resting state networks,” J. Neurosci. Off. J. Soc. Neurosci., vol. 34, no. 11, pp. 3982–3992, 2014, doi: 10.1523/JNEUROSCI.4341-13.2014.

[50] R. M. C. Spencer, M. Sunm, and R. B. Ivry, “Sleep-Dependent Consolidation of Contextual Learning,” Curr. Biol., vol. 16, no. 10, pp. 1001–1005, May 2006, doi: 10.1016/J.CUB.2006.03.094.

[51] A. B. Fitzroy, K. A. Kainec, J. Seo, and R. M. C. Spencer, “Encoding and consolidation of motor sequence learning in young and older adults,” Neurobiol. Learn. Mem., vol. 185, p. 107508, Nov. 2021, doi: 10.1016/J.NLM.2021.107508.

[52] J. A. Horne and O. Ostberg, “A self-assessment questionnaire to determine morningness-eveningness in human circadian rhythms - PubMed,” Int. J. Chronobiol., vol. 4, no. 2, pp. 97–110, 1976.

[53] D. J. Buysse, C. F. Reynolds, T. H. Monk, S. R. Berman, and D. J. Kupfer, “The Pittsburgh Sleep Quality Index: a new instrument for psychiatric practice and research,” Psychiatry Res., vol. 28, no. 2, pp. 193–213, 1989, doi: 10.1016/0165-1781(89)90047-4.

[54] R. C. Oldfield, “The assessment and analysis of handedness: the Edinburgh inventory,” Neuropsychologia, vol. 9, no. 1, pp. 97–113, 1971, doi: 10.1016/0028-3932(71)90067-4.

[55] S. Ocklenburg, P. Friedrich, O. Güntürkün, and E. Genç, “Voxel-wise grey matter asymmetry analysis in left-and right-handers,” Neurosci. Lett., vol. 633, pp. 210–214, Oct. 2016, doi: 10.1016/j.neulet.2016.09.046.

[56] X.-Z. Kong et al., “Mapping cortical brain asymmetry in 17,141 healthy individuals worldwide via the ENIGMA Consortium,” Proc. Natl. Acad. Sci. U. S. A., vol. 115, no. 22, pp. E5154–E5163, May 2018, doi: 10.1073/pnas.1718418115.

[57] Z. Sha et al., “Handedness and its genetic influences are associated with structural asymmetries of the cerebral cortex in 31,864 individuals,” Proc. Natl. Acad. Sci. U. S. A., vol. 118, no. 47, p. e2113095118, Nov. 2021, doi: 10.1073/pnas.2113095118.

[58] E. M. Robertson, “The Serial Reaction Time Task: Implicit Motor Skill Learning?,” J. Neurosci., vol. 27, no. 38, pp. 10073–10075, Sep. 2007, doi: 10.1523/JNEUROSCI.2747-07.2007.

[59] B. W. Ellis, M. W. Johns, R. Lancaster, P. Raptopoulos, N. Angelopoulos, and R. G. Priest, “The St. Mary’s Hospital sleep questionnaire: a study of reliability,” Sleep, vol. 4, no. 1, pp. 93–97, 1981, doi: 10.1093/SLEEP/4.1.93.

[60] L. Genzel et al., “Sex and modulatory menstrual cycle effects on sleep related memory consolidation,” Psychoneuroendocrinology, vol. 37, no. 7, pp. 987–998, Jul. 2012, doi: 10.1016/J.PSYNEUEN.2011.11.006.

[61] K. Ikarashi, D. Sato, K. Iguchi, Y. Baba, and K. Yamashiro, “Menstrual Cycle Modulates Motor Learning and Memory Consolidation in Humans,” Brain Sci. 2020 Vol 10 Page 696, vol. 10, no. 10, p. 696, Oct. 2020, doi: 10.3390/BRAINSCI10100696.

[62] C. Jiang et al., “Diurnal Microstructural Variations in Healthy Adult Brain Revealed by Diffusion Tensor Imaging,” PLoS ONE, vol. 9, no. 1, p. e84822, Jan. 2014, doi: 10.1371/journal.pone.0084822.

[63] A. Trefler, N. Sadeghi, A. G. Thomas, C. Pierpaoli, C. I. Baker, and C. Thomas, “Impact of time-of-day on brain morphometric measures derived from T1-weighted magnetic resonance imaging,” NeuroImage, vol. 133, pp. 41–52, Jun. 2016, doi: 10.1016/j.neuroimage.2016.02.034.

[64] J. Lim and D. F. Dinges, “Sleep Deprivation and Vigilant Attention,” Ann. N. Y. Acad. Sci., vol. 1129, no. 1, pp. 305–322, May 2008, doi: 10.1196/ANNALS.1417.002.

[65] M. Delacre, D. Lakens, and C. Leys, “Why psychologists should by default use welch’s t-Test instead of student’s t-Test,” Int. Rev. Soc. Psychol., vol. 30, no. 1, pp. 92–101, 2017, doi: 10.5334/IRSP.82.

[66] M. Delacre, C. Leys, Y. L. Mora, and D. Lakens, “Taking parametric assumptions seriously: Arguments for the use of welch’s f-test instead of the classical f-test in one-way ANOVA,” Int. Rev. Soc. Psychol., vol. 32, no. 1, 2020, doi: 10.5334/IRSP.198/GALLEY/161/DOWNLOAD/.

[67] M. F. Glasser et al., “The minimal preprocessing pipelines for the Human Connectome Project,” NeuroImage, vol. 80, pp. 105–124, Oct. 2013, doi: 10.1016/J.NEUROIMAGE.2013.04.127.

[68] L. Henschel, S. Conjeti, S. Estrada, K. Diers, B. Fischl, and M. Reuter, “FastSurfer - A fast and accurate deep learning based neuroimaging pipeline,” NeuroImage, vol. 219, p. 117012, Oct. 2020, doi: 10.1016/J.NEUROIMAGE.2020.117012.

[69] J. Scholz, M. C. Klein, T. E. J. Behrens, and H. Johansen-Berg, “Training induces changes in white-matter architecture,” Nat. Neurosci., vol. 12, no. 11, pp. 1370–1371, Nov. 2009, doi: 10.1038/nn.2412.

[70] R. S. Desikan et al., “An automated labeling system for subdividing the human cerebral cortex on MRI scans into gyral based regions of interest,” NeuroImage, vol. 31, no. 3, pp. 968–980, Jul. 2006, doi: 10.1016/J.NEUROIMAGE.2006.01.021.

[71] B. Fischl et al., “Whole brain segmentation: Automated labeling of neuroanatomical structures in the human brain,” Neuron, vol. 33, no. 3, pp. 341–355, Jan. 2002, doi: 10.1016/S0896-6273(02)00569-X.

[72] B. Fischl et al., “Automatically parcellating the human cerebral cortex,” Cereb. Cortex N. Y. N 1991, vol. 14, no. 1, pp. 11–22, Jan. 2004, doi: 10.1093/CERCOR/BHG087.

[73] B. Fischl, “FreeSurfer,” NeuroImage, vol. 62, no. 2, pp. 774–781, Aug. 2012, doi: 10.1016/J.NEUROIMAGE.2012.01.021.

[74] J. L. R. Andersson, S. Skare, and J. Ashburner, “How to correct susceptibility distortions in spin-echo echo-planar images: Application to diffusion tensor imaging,” NeuroImage, vol. 20, no. 2, pp. 870–888, Oct. 2003, doi: 10.1016/S1053-8119(03)00336-7.

[75] J. L. R. Andersson and S. N. Sotiropoulos, “An integrated approach to correction for off-resonance effects and subject movement in diffusion MR imaging,” NeuroImage, vol. 125, pp. 1063–1078, Jan. 2016, doi: 10.1016/J.NEUROIMAGE.2015.10.019.

[76] J. H. Jensen and J. A. Helpern, “MRI Quantification of Non-Gaussian Water Diffusion by Kurtosis Analysis,” NMR Biomed., vol. 23, no. 7, p. 698, Aug. 2010, doi: 10.1002/NBM.1518.

[77] M. Guerreri et al., “Revised NODDI model for diffusion MRI data with multiple b-tensor encodings,” Proc. Jt. Annu. Meet. ISMRM-ESMRMB Int. Soc. Magn. Reson. Med. 2018, Jun. 2018, Accessed: Apr. 15, 2023. [Online]. Available: http://archive.ismrm.org/2018/5241.html

[78] M. Guerreri, F. Szczepankiewicz, B. Lampinen, M. Palombo, M. Nilsson, and H. Zhang, “Tortuosity assumption not the cause of NODDI’s incompatibility with tensor-valued diffusion encoding,” 2020.

[79] D. N. Greve and B. Fischl, “Accurate and robust brain image alignment using boundary-based registration,” NeuroImage, vol. 48, no. 1, pp. 63–72, Oct. 2009, doi: 10.1016/J.NEUROIMAGE.2009.06.060.

[80] C. S. Parker et al., “Not all voxels are created equal: Reducing estimation bias in regional NODDI metrics using tissue-weighted means,” Neuroimage, vol. 245, p. 118749, Dec. 2021, doi: 10.1016/J.NEUROIMAGE.2021.118749.

[81] D. N. Greve and B. Fischl, “False positive rates in surface-based anatomical analysis,” NeuroImage, vol. 171, pp. 6–14, May 2018, doi: 10.1016/J.NEUROIMAGE.2017.12.072.

[82] R. B. Berry et al., “AASM Scoring Manual Updates for 2017 (Version 2.4),” J. Clin. Sleep Med. JCSM Off. Publ. Am. Acad. Sleep Med., vol. 13, no. 5, pp. 665–666, 2017, doi: 10.5664/JCSM.6576.

[83] S. Diekelmann, S. Biggel, B. Rasch, and J. Born, “Offline consolidation of memory varies with time in slow wave sleep and can be accelerated by cuing memory reactivations,” Neurobiol. Learn. Mem., vol. 98, no. 2, pp. 103–111, Sep. 2012, doi: 10.1016/J.NLM.2012.07.002.

[84] A. C. M. Koopman^1^, et al., “Targeted memory reactivation of a serial reaction time task in SWS, but not REM, preferentially benefits the non-dominant hand,” bioRxiv, p. 2020.11.17.381913, Nov. 2020, doi: 10.1101/2020.11.17.381913.

[85] B. Rasch, C. Büchel, S. Gais, and J. Born, “Odor cues during slow-wave sleep prompt declarative memory consolidation,” Science, vol. 315, no. 5817, pp. 1426–1429, Mar. 2007, doi: 10.1126/science.1138581.

[86] C. Zelano and N. Sobel, “Humans as an Animal Model for Systems-Level Organization of Olfaction,” Neuron, vol. 48, no. 3, pp. 431–454, Nov. 2005, doi: 10.1016/J.NEURON.2005.10.009.

[87] S. I. R. Pereira, F. Beijamini, F. D. Weber, R. A. Vincenzi, F. A. C. da Silva, and F. M. Louzada, “Tactile stimulation during sleep alters slow oscillation and spindle densities but not motor skill,” Physiol. Behav., vol. 169, pp. 59–68, Feb. 2017, doi: 10.1016/J.PHYSBEH.2016.11.024.

[88] M. P. Veldman, N. Dolfen, M. A. Gann, J. Carrier, B. R. King, and G. Albouy, “Somatosensory Targeted Memory Reactivation Modulates Oscillatory Brain Activity but not Motor Memory Consolidation,” Neuroscience, vol. 465, pp. 203–218, Jun. 2021, doi: 10.1016/J.NEUROSCIENCE.2021.03.027.

[89] D. Denis and S. A. Cairney, “Neural reactivation during human sleep,” Emerg. Top. Life Sci., vol. 7, no. 5, pp. 487–498, Dec. 2023, doi: 10.1042/ETLS20230109.

[90] R. Cox, D. S. Mylonas, D. S. Manoach, and R. Stickgold, “Large-scale structure and individual fingerprints of locally coupled sleep oscillations,” Sleep, vol. 41, no. 12, pp. 1–15, Dec. 2018, doi: 10.1093/SLEEP/ZSY175.

[91] R. Cox, J. Van Driel, M. De Boer, and L. M. Talamini, “Slow Oscillations during Sleep Coordinate Interregional Communication in Cortical Networks,” J. Neurosci., vol. 34, no. 50, pp. 16890–16901, Dec. 2014, doi: 10.1523/JNEUROSCI.1953-14.2014.

[92] M. A. Hahn, K. Bothe, D. Heib, M. Schabus, R. F. Helfrich, and K. Hoedlmoser, “Slow oscillation–spindle coupling strength predicts real-life gross-motor learning in adolescents and adults,” eLife, vol. 11, Feb. 2022, doi: 10.7554/ELIFE.66761.

[93] C. Mikutta et al., “Phase-amplitude coupling of sleep slow oscillatory and spindle activity correlates with overnight memory consolidation,” J. Sleep Res., vol. 28, no. 6, p. e12835, Dec. 2019, doi: 10.1111/JSR.12835.

[94] J. W. Antony, L. Piloto, M. Wang, P. Pacheco, K. A. Norman, and K. A. Paller, “Sleep Spindle Refractoriness Segregates Periods of Memory Reactivation,” Curr. Biol., vol. 28, no. 11, pp. 1736–1743.e4, Jun. 2018, doi: 10.1016/j.cub.2018.04.020.

[95] J. W. Antony, M. Schönauer, B. P. Staresina, and S. A. Cairney, “Sleep Spindles and Memory Reprocessing,” Trends Neurosci., vol. 42, no. 1, pp. 1–3, Jan. 2019, doi: 10.1016/j.tins.2018.09.012.

[96] T. Schreiner and B. Rasch, “Boosting Vocabulary Learning by Verbal Cueing During Sleep,” Cereb. Cortex N. Y. N 1991, vol. 25, no. 11, pp. 4169–4179, Nov. 2015, doi: 10.1093/CERCOR/BHU139.

[97] J. Farthouat, M. Gilson, and P. Peigneux, “New evidence for the necessity of a silent plastic period during sleep for a memory benefit of targeted memory reactivation,” Sleep Spindl. Cortical States, vol. 1, no. 1, pp. 14–26, Mar. 2017, doi: 10.1556/2053.1.2016.002.

[98] M. Geva-Sagiv et al., “Augmenting hippocampal–prefrontal neuronal synchrony during sleep enhances memory consolidation in humans,” Nat. Neurosci. 2023 266, vol. 26, no. 6, pp. 1100–1110, Jun. 2023, doi: 10.1038/s41593-023-01324-5.

[99] T. Schreiner, M. Petzka, T. Staudigl, and B. P. Staresina, “Endogenous memory reactivation during sleep in humans is clocked by slow oscillation-spindle complexes,” Nat. Commun. 2021 121, vol. 12, no. 1, pp. 1–10, May 2021, doi: 10.1038/s41467-021-23520-2.

[100] L. J. Batterink, J. D. Creery, and K. A. Paller, “Phase of Spontaneous Slow Oscillations during Sleep Influences Memory-Related Processing of Auditory Cues,” J. Neurosci., vol. 36, no. 4, pp. 1401–1409, Jan. 2016, doi: 10.1523/JNEUROSCI.3175-15.2016.

[101] M. Göldi, E. A. M. van Poppel, B. Rasch, and T. Schreiner, “Increased neuronal signatures of targeted memory reactivation during slow-wave up states,” Sci. Rep. 2019 91, vol. 9, no. 1, pp. 1–10, Feb. 2019, doi: 10.1038/s41598-019-39178-2.

[102] G. Albouy, B. R. King, P. Maquet, and J. Doyon, “Hippocampus and striatum: Dynamics and interaction during acquisition and sleep-related motor sequence memory consolidation,” Hippocampus, vol. 23, no. 11, pp. 985–1004, Nov. 2013, doi: 10.1002/hipo.22183.

[103] K. Debas et al., “Off-line consolidation of motor sequence learning results in greater integration within a cortico-striatal functional network,” NeuroImage, vol. 99, pp. 50–58, Oct. 2014, doi: 10.1016/J.NEUROIMAGE.2014.05.022.

[104] J. Viczko, V. Sergeeva, L. B. Ray, A. M. Owen, and S. M. Fogel, “Does sleep facilitate the consolidation of allocentric or egocentric representations of implicitly learned visual-motor sequence learning?,” Learn. Mem. Cold Spring Harb. N, vol. 25, no. 2, pp. 67–77, Feb. 2018, doi: 10.1101/lm.044719.116.

[105] C. Cajochen, V. Knoblauch, A. Wirz-Justice, K. Kräuchi, P. Graw, and D. Wallach, “Circadian modulation of sequence learning under high and low sleep pressure conditions,” Behav. Brain Res., vol. 151, no. 1–2, pp. 167–176, May 2004, doi: 10.1016/j.bbr.2003.08.013.

[106] D. A. Cohen, A. Pascual-Leone, D. Z. Press, and E. M. Robertson, “Off-line learning of motor skill memory: A double dissociation of goal and movement,” Proc. Natl. Acad. Sci., vol. 102, no. 50, pp. 18237–18241, Dec. 2005, doi: 10.1073/pnas.0506072102.

[107] D. B. Willingham, M. J. Nissen, and P. Bullemer, “On the Development of Procedural Knowledge,” J. Exp. Psychol. Learn. Mem. Cogn., vol. 15, no. 6, pp. 1047–1060, 1989, doi: 10.1037/0278-7393.15.6.1047.

[108] D. J. Sanchez, E. W. Gobel, and P. J. Reber, “Performing the unexplainable: Implicit task performance reveals individually reliable sequence learning without explicit knowledge,” Psychon. Bull. Rev., vol. 17, no. 6, pp. 790–796, Dec. 2010, doi: 10.3758/PBR.17.6.790/METRICS.

