## Supplementary Information for "Shaping the structural dynamics of motor learning through cueing during sleep"

### 1. Explicit sequence knowledge assessment

To determine whether sequence knowledge obtained through repeated practice on the SRTT was implicit or explicit, we used a modified version of Jacoby's Process Dissociation Procedure [1]. Participants were instructed to generate the learned sequence under two different conditions: *Inclusion* and *Exclusion*. In the *Inclusion* condition, participants were instructed to reproduce the sequence they had practiced during the SRTT. Successful performance in this condition could result from either explicit process (i.e., consciously recalling the training sequence) or implicit knowledge (i.e., having an intuitive sense of the next key press despite thinking they are guessing). Both explicit recollection and intuitive responses contribute to improved performance in the *Inclusion* condition, as both types of knowledge enhance accurate sequence generation [2]. In contrast, in the *Exclusion* condition, participants were instructed to perform the sequence in the reverse order. Here, implicit and explicit knowledge operate in opposition. Explicit knowledge allows participants to intentionally avoid the learned sequence, thus succeeding in the *Exclusion* task. However, if participants continue to produce chunks of the previously practiced sequence despite being instructed to perform another sequence (here, the sequence in reversed order), this indicates the predominant influence of implicit knowledge, which cannot be consciously controlled [2].To analyze performance in both conditions, we calculated the percentage of correct triplets throughout the 96 keypresses, considering that humans tend to chunk behavioral sequences into segments of [3], [4] up to 3 elements [5]. High accuracy in the *Inclusion* condition would indicate proficient learning, while low accuracy in the *Exclusion* condition would suggest a high explicit knowledge. The ANOVA performed on accuracy scores with as within-subject factor *Generation* (Inclusion vs. Exclusion) and between-subject factor *Condition* (RS vs. TMR) disclosed a main *Generation* ($F_{1, 58} = 148.433$, $p < 0.001$, $\eta_p^2 = 0.719$) and a main *Condition* effect ($F_{1, 58} = 6.715$, $p = 0.012$, $\eta_p^2 = 0.104$). However, no significant *Generation *Condition* interaction effect ($F_{1, 58} = 2.089$, $p = 0.154$, $\eta_p^2 = 0.035$) was found. Additionally, we computed an estimate of explicit learning by subtracting exclusion from inclusion scores (see [2]). This estimate significantly differed from 0 in both groups ($p < 0.001$), indicating the presence of explicit knowledge in both RS and TMR *Condition*s. However, explicit knowledge did not differ between both *Condition*s ($t_{57.830} = 1.445$, $p = 0.154$, Cohen's d = 0.263).

### 2. Global Performance Index (GPI) calculation

The GPI was calculated for each block with the following formula [6]:

$$GPI = e^{-speed} * e^{-accuracy}$$

with

$$speed_{block} = mean\ RT\ (in\ seconds)$$

$$accuracy_{block} = \frac{number\ of\ correct\ triplets}{94} * 100$$

94 being the maximal number of correct triplets possible as one block counts 96 keypresses. Here, *e* stands for Euler' s number, the mathematical constant (~2.7183).

### 3. Detailed Behavioural Results

#### 3.1. Learning Session – online gains

##### *3.1.1. GPI measures*

For motor learning at Day 1, the ANOVA computed on GPI performance for sequential blocks performed with within-subject factor *Block* (1:25, 27:30) and between-subject factor *Sleep* (RS vs. TMR) disclosed a main *Block* effect ($F_{7.166,\ 408,436}$ = 38.043, $p$ < 0.001, $\eta_p^2$ = 0.400) characterized by progressive increase in GPI with task practice. Also, as expected, no main *Condition* ($F_{1,\ 57}$ = 0.940, $p$ = 0.336, $\eta_p^2$ = 0.016) nor *Block*Condition* interaction ($F_{7.166,\ 408,436}$ = 1.029, $p$ = 0.410, $\eta_p^2$ = 0.018) was found as the experimental manipulation did not happen yet at this stage. Thus, both groups exhibited a similar performance increase over task practice in the initial learning session. Additionally, a separate ANOVA was conducted with within-subject factor *Block* type (sequential blocks 25 & 27 vs random block 26) and between-subject factor *Condition* (RS vs. TMR). This analysis disclosed a main *Block* effect ($F_{1.699,\ 96.817}$ = 88.311, $p$ < 0.001, $\eta_p^2$ = 0.608), and post-hoc tests showed that RT in pseudo-random block 26 was significantly slower than in sequential block 25 ($p_{corr}$ < 0.001) and 27 ($p_{corr}$ < 0.001). It indicates that participants learned the sequence and started anticipating the upcoming position in sequential blocks, and that performance improvement was not merely due to motor practice. As previously, no main effect of *Condition* ($F_{1,\ 57}$ = 0.146, $p$ = 0.703, $\eta_p^2$ = 0.003) or *Block*Condition* interaction ($F_{1.699,\ 96.817}$ = 3.055, $p$ = 0.060, $\eta_p^2$ = 0.051) was found.

#### *3.1.2. Reaction time measures*

Similar ANOVAs were calculated on mean RT measures. The ANOVA performed on mean RT for sequential blocks with within-subject factor *Block* (1:25, 27:30) and between-subject factor *Condition* (RS vs. TMR) disclosed a main *Block* effect ($F_{3.918,\ 223,327} = 86.126$, $p < 0.001$, $\eta_p^2 = 0.602$) characterized by progressive decrease in mean RT with task practice. Also, as expected, no *Block*Condition* interaction ($F_{3.918,\ 223,327} = 1.663$, $p = 0.161$, $\eta_p^2 = 0.028$) was found and only a marginal *Condition* effect ($F_{1,\ 57} = 3.748$, $p = 0.058$, $\eta_p^2 = 0.062$). Therefore, we also conducted a separate ANOVA comparing directly the means of the 2 last *Blocks* (29:30) of the learning session between both *Conditions* (RS vs. TMR). This analysis revealed a significant difference in RTs at the end of the learning between both conditions ($F_{1,\ 54.373} = 5.632$, $p = 0.021$, $\eta_p^2 = 0.090$) while this was not expected yet. Also, the additional ANOVA conducted with within-subject factor the *Block* type (sequential blocks 25 & 27 vs random block 26) and between-subject factor *Condition* (RS vs. TMR) revealed a main *Block* effect ($F_{1.731,98.674} = 175.577$, $p < 0.001$, $\eta_p^2 = 0.755$), and post-hoc tests showed that RT in pseudo-random block 26 was significantly slower than in sequential block 25 ($p_{corr} < 0.001$) and 27 ($p_{corr} < 0.001$). However, as expected, no *Condition* ($F_{1,\ 57} = 3.748$, $p = 0.058$, $\eta_p^2 = 0.062$) or *Block*Condition* interaction ($F_{1.731,98.674} = 2.531$, $p = 0.092$, $\eta_p^2 = 0.043$) was found (see Figure S1).

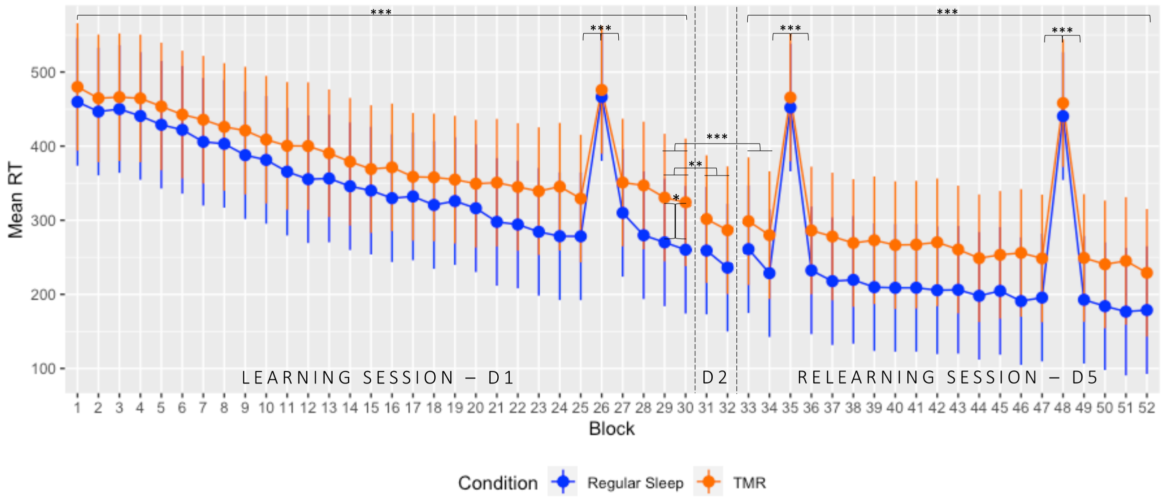


*Figure S1: SRTT : Changes in mean reaction time across the entire protocol*

Mean Reaction Time (RT) ± standard deviation plotted for all blocks performed over the three testing days. On the first day, participants completed 30 blocks in a learning session (D1; block 26 pseudo-random). Following the experimental night between Days 1 and 2, performance was reassessed on the morning of Day 2 for two blocks (D2; blocks 31-32). After three nights of sleep at home, participants returned to perform the task again for 20 blocks in a relearning session on Day 5 (D5; blocks 35 and 48 being pseudo-random). ***p < 0.001.

#### 3.1.3. *Accuracy measures*

Lastly, comparable ANOVAs were performed on accuracy measures (percentage of correct triplets throughout the 96 keypresses, as humans show a natural tendency to divide behavioural sequences in chunks [3], [4] up to 3 elements [5]). No main *Block* ($F_{13.314,\ 758.922}$ = 0.928; $p$ = 0.525, $\eta_p^2$ = 0.016), *Condition* ($F_{1,\ 57}$ = 2.441, $p$ = 0.124, $\eta_p^2$ = 0.041), or *Block*Condition* interaction ($F_{13.314,\ 758.922}$ = 1.472, $p$ = 0.120, $\eta_p^2$ = 0.025) effects were found across sequential blocks, indicating that accuracy remained stable over the learning session. However, we also performed a separate ANOVA comparing directly the means of the 2 last *Blocks* (29:30) of the learning session between both *Conditions* (RS vs. TMR) to confirm the hypothesis participants used a different strategy when performed the task by prioritizing either speed or accuracy. This analysis revealed a significant difference in accuracy at the end of the learning between both conditions ($F_{1,\ 53.831}$ = 9.172, $p$ = 0.004, $\eta_p^2$ = 0.137) with a significantly higher accuracy in the TMR group compared to the RS group. Also, no difference in accuracy was found when comparing pseudo-random block 26 with sequential blocks 25 or 27 (main *Block* effect: $F_{1.814,\ 103.389}$ = 0.048, $p$ = 0.941, $\eta_p^2$ = 8.332e$^{-4}$; main *Condition* effect $F_{1,\ 57}$ = 3.816, $p$ = 0.056, $\eta_p^2$ = 0.063; *Block*Condition* interaction ($F_{1.814,\ 103.389}$ = 1.996, $p$ = 0.145, $\eta_p^2$ = 0.034; see Figure S2).

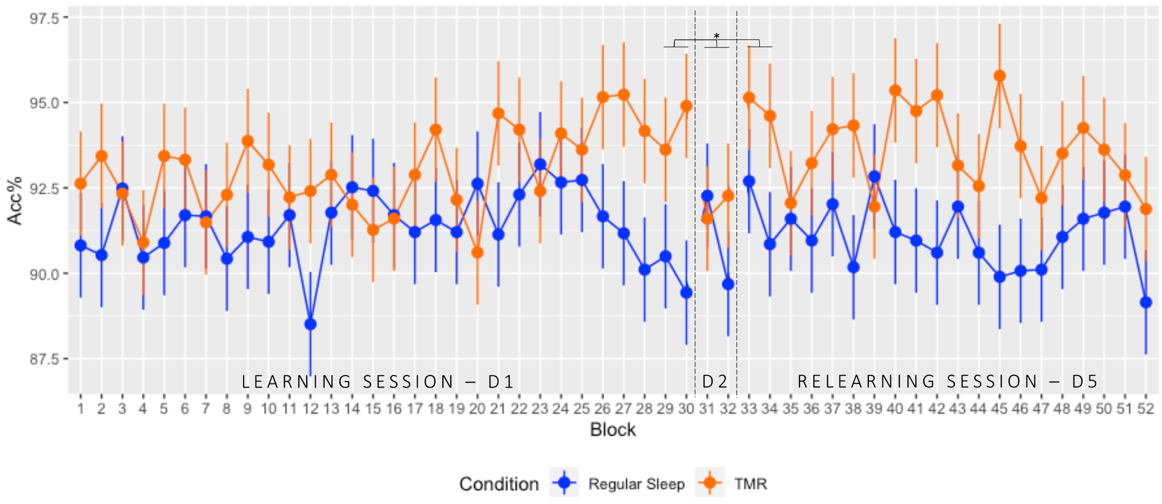


*Figure S2: Variations in accuracy throughout the entire protocol*

Accuracy (expressed as a percentage) ± standard deviation plotted for all blocks across the three testing days. On Day 1, participants completed 30 blocks during a learning session (D1; block 26 pseudo-random). After the experimental night between Days 1 and 2, accuracy was reassessed on the morning of Day 2 for two blocks (D2; blocks 31-32). After three nights of rest at home, participants returned to complete 20 blocks in a relearning session on Day 5 (D5; blocks 35 and 48 being pseudo-random). *p < 0.05.

#### 3.2. Offline gains between D1, D2 and D5

A mixed ANOVA looking at the evolution of GPI between the 2 last blocks of the learning session (LS D1; mean blocks 29:30), the 2 blocks performed during retest on Day 2 after the experimental night (RE D2; mean blocks 31:32) and the 2 first blocks of the relearning session (RL D5; mean blocks 33:34) with between-subject factor *Condition* (RS vs. TMR) disclosed a main *Day* ($F_{2, 114}$ = 11.944, $p < 0.001$, $\eta_p^2$ = 0.173) effect with no significant change in GPI between D1 and D2 ($p_{corr}$ = 0.070), but a significant increase at D5 (all $p_{corr}s$ < 0.033). However, no *Condition* effect ($F_{1, 57}$ = 0.799, $p$ = 0.375, $\eta_p^2$ = 0.014), nor the *Day*Condition* interaction was significant ($F_{2, 114}$ = 1.087, $p$ = 0.341, $\eta_p^2$ = 0.019) were significant.

A similar mixed ANOVA was performed on mean RTs with as within-subject factor *Day* (D1, D2 and D5) and with between-subject factor *Condition* (RS vs. TMR). A main *Day* ($F_{1.685, 96.022}$ = 19.349, $p < 0.001$, $\eta_p^2$ = 0.253) effect with a significant decrease in mean RT between D1 and D2 ($p_{corr} < 0.001$) or D5 ($p_{corr} < 0.001$), but no difference between D2 and D5 ($p_{corr}$ = 1.000). Additionally, the effect of *Condition* effect ($F_{1, 57}$ = 4.702, $p$ = 0.034, $\eta_p^2$ = 0.076) revealed significantly lower reaction times for the RS group compared to the TMR group. However, the *Day*Condition* interaction was not significant ($F_{1.685, 96.022}$ = 1.083, $p$ = 0.334, $\eta_p^2$ = 0.019; see Figure S1).

Concerning accuracy, a similar ANOVA revealed no significant *Day*Sleep* interaction ($F_{2, 114}$ = 2.911, $p$ = 0.058, $\eta_p^2$ = 0.049), but a main *Day* ($F_{2, 114}$ = 3.675, $p$ = 0.028, $\eta_p^2$ = 0.061) effect with a significant difference between D2 and D5 ($p_{corr}$ = 0.027), but not between D1 and D2 ($p_{corr}$ = 1.000) or D1 and D5 ($p_{corr}$ = 0.223). Also, a significant effect of *Condition* ($F_{1, 57}$ = 4.977, $p$ = 0.030, $\eta_p^2$ = 0.080) was found with a significantly lower accuracy for the RS group compared to the TMR group (see Figure S2).

#### 3.3. Relearning session – online gains

Looking at motor sequence relearning at Day 5, an ANOVA on GPI for all sequential blocks with within-subject factor *Block* (33:34, 36:47, 49:52; random blocks being numbered 35 & 48) and between-subject factor *Condition* (RS vs. TMR) disclosed a main *Block* effect ($F_{10.617, 615.762}$ = 3.929, $p$ < 0.001, $\eta_p^2$ = 0.063) with an increase in GPI over the sequential blocks. However, no main *Condition* ($F_{1, 58}$ = 1.023, $p$ = 0.316, $\eta_p^2$ = 0.017), nor *Block*Condition* interaction ($F_{10.617, 615.762}$ = 1.473, $p$ = 0.140, $\eta_p^2$ = 0.025) was found, suggesting that the presence of auditory cues during the post-learning night did not impact the behavioural time course for the practice on previously learned material. Besides, two separate ANOVAs comparing both pseudo-random blocks with their preceding and following sequential block respectively were conducted with within-subject factor *Block* (34:36 or 47:49) and between-subject factor *Condition* (RS vs. TMR). Both analyses disclosed lower GPI in pseudo-random blocks 35 and 48 than in the surrounding sequential blocks (all $p_{corr}$s < 0.001).

The ANOVA on mean reaction time (RT) for all sequential blocks with within-subject factor *Block* (33:34, 36:47, 49:52; random blocks being numbered 35 & 48) and between-subject factor *Condition* (RS vs. TMR) disclosed a main *Block* effect ($F_{8.448, 490.006}$ = 33.412, $p$ < 0.001, $\eta_p^2$ = 0.366) with a decrease in mean RT over the sequential blocks. Also, a main *Condition* ($F_{1, 58}$ = 5.291, $p$ = 0.025, $\eta_p^2$ = 0.084) effect emerged which was expected considering both groups already showed different RTs at the end of learning. However, the *Block*Condition* interaction ($F_{8.448, 490.006}$ = 1.265, $p$ = 0.257, $\eta_p^2$ = 0.021) was found, suggesting that auditory cueing did not impact the behavioural time course for the practice on previously learned material. Besides, two separate ANOVAs comparing both pseudo-random blocks with their preceding and following sequential block respectively were conducted with within-subject factor *Block* (34:36 or 47:49) and between-subject factor *Condition* (RS vs. TMR). Both analyses disclosed slower RTs in pseudo-random blocks 35 and 48 than in the surrounding sequential blocks (all $p_{corr}$s < 0.001; see Figure S1).

Regarding accuracy measures, no main *Block* ($F_{10.986, 637.216}$ = 1.189; $p$ = 0.291, $\eta_p^2$ = 0.020), or *Block*Condition* interaction ($F_{10.986, 637.216}$ = 1.139, $p$ = 0.328, $\eta_p^2$ = 0.019) effects were found across sequential blocks. However, a significant *Condition* ($F_{1, 58}$ = 7.650, $p$ = 0.008, $\eta_p^2$ = 0.117) revealed a higher accuracy in the TMR group compared to the RS group. Also, no difference in accuracy was found when comparing pseudo-random block 35 with sequential blocks 34 or

36 (all $p_{corr}$s > 0.128), neither when comparing pseudo-random block 48 with sequential blocks 47 or 49 (all $p_{corr}$s > 0.060; see Figure S2).

### 4. Additional EEG analyses on the two first sleep cycles

After all sleep recordings were manually scored following the protocol described in the Methods section, 106 out of 256 EEG channels were selected to ensure good, consistent scalp coverage. These channels were subsequently scanned by an automated artifact detection algorithm [7] to flag channels with unusually high or low standard deviations and those that were noisy, flat or contained high-amplitude peaks. We also identified bridged channels by detecting neighbouring electrodes with high correlation [8]. Only channels with artifact-free signals for at least 95% of the time across both nights were retained for further analysis. This selection process resulted in retaining between 89 and 106 consistent channels across both nights for each participant (mean number of channels for Group 2 = 98.52; for Group 3 = 98.43). A final check confirmed homogeneous scalp coverage across channels, with no significant differences observed.

After preprocessing, spindle events were detected automatically using the algorithm from [9]. Spindles were only included in further analyses if they were simultaneously present on 10-40% of the previously selected channels; spindles appearing on less than 10% of channels were considered highly localized events, while those on more than 40% were flagged as potential artifacts. Spindle density was calculated by dividing the number of spindles detected during N2 and N3 sleep by the total duration of these sleep stages. Slow oscillations were identified using the same preprocessing pipeline but automatically detected [10] with the standard parameters found in [11].

To examine the effect of TMR on spindle and SO densities, we conducted two separate 2x2 ANOVA with *Night* (habituation vs. experimental) as a within-subject factor and *Group* (RS vs. TMR) as a between-subject factor. For spindle density, results indicated no significant effect of *Night* ($F_{1, 58}$ = 1.079; $p$ = 0.303) or *Group* ($F_{1, 58}$ = 0.957; $p$ = 0.332), and no significant interaction effect ($F_{1, 58}$ = 1.596; $p$ = 0.212). In contrast, the ANOVA on SO density revealed a significant main effect of both *Night* ($F_{1, 58}$ = 8.079; $p$ = 0.006) and *Group* ($F_{1, 58}$ = 6.552; $p$ = 0.013), along with a significant interaction effect ($F_{1, 58}$ = 24.848; $p$ < 0.001). Both groups exhibited similar

SO density during the habituation night. However, on the experimental night, the RS group maintained a stable SO density as compared to the first night, while the TMR group showed a pronounced decrease.

These results indicate that while acoustic stimulation during sleep did not significantly influence spindle density, it led to a substantial reduction in SO density for the TMR group during the first two sleep cycles on the experimental night, with SO density remaining stable in the RS group.

### 5. Additional (control) Analyses

#### 5.1. Alertness before task performance

An ANOVA computed on Reciprocal Reaction Time (RRT; 1/RT) in the PVT5 [12] with within-subject factor *PVT Session* (D1, D2, D5) and between-subject factor *Condition* (RS vs. TMR) found no main effect of *Condition* ($F_{1, 58} = 1.799$, $p = 0.185$, $\eta_p^2 = 0.030$), *PVT Session* ($F_{1.364, 79.096} = 0.858$, $p = 0.390$, $\eta_p^2 = 0.015$), nor a *Condition*PVT Session* interaction ($F_{1.364, 79.096} = 0.519$, $p = 0.529$, $\eta_p^2 = 0.009$) effects, suggesting vigilance remained stable and similar for both groups throughout the entire protocol.

#### 5.2. Sleep Quality and Duration during the protocol

The ANOVA performed on sleep quality with within-subject factor *Night* (1:7) and between-subject factor *Condition* (RS vs. TMR) disclosed a main effect of *Night* ($F_{6, 342} = 24.439$, $p < 0.001$, $\eta_p^2 = 0.300$) and a *Night*Condition* interaction effect ($F_{6, 342} = 3.246$, $p = 0.004$, $\eta_p^2 = 0.054$) with a significantly lower sleep quality on the first (all $p_{corr}$s < 0.001) and fourth (all $p_{corr}$s < 0.046) night that correspond respectively to the habituation and experimental night spend at the lab under hd-EEG compared to the other nights that were spent at home. However, no main effect of *Condition* ($F_{1, 57} = 1.272e^{-4}$, $p = 0.991$, $\eta_p^2 = 2.232e^{-6}$) was highlighted, suggesting both groups experienced a similar sleep quality throughout the protocol.

A similar ANOVA conducted on sleep duration with within-subject factor *Night* (1:7) and between-subject factor *Condition* (RS vs. TMR) revealed no main *Condition* ($F_{1, 57} = 0.024$, $p = 0.876$, $\eta_p^2 = 4.296e^{-4}$), *Night* ($F_{4.296, 244.893} = 1.991$, $p = 0.091$, $\eta_p^2 = 0.034$) effect, nor *Night*Condition* interaction ($F_{4.296, 244.893} = 1.920$, $p = 0.103$, $\eta_p^2 = 0.033$).

### 6. Diffusion Weighted Imaging (DWI) Data

#### 6.1. Sleep Quality and Duration during the Motor training-related short-term structural changes (Day 1; DWI1 vs. DWI2)

##### 6.1.1. Cortical Ribbon

Significant clusters were identified based on cluster-wise p-values under the predetermined significance threshold of alpha = 0.05 (CWP < 0.05).

The surface-based statistical analysis on DTI parameters disclosed a *Learning* (Pre $DWI_1$ vs. Post $DWI_2$) effect following the learning session, with bilateral decrease in MD in the precuneus, the superior parietal, lateral occipital, paracentral, precentral and postcentral gyri, the insula, the superior frontal, inferior parietal, supramarginal, and inferior temporal gyri, the banks of the superior temporal sulcus, the lingual, rostral anterior cingulate, superior temporal, fusiform, and caudal anterior cingulate gyri, the cuneus, the transverse temporal, lateral orbitofrontal and posterior cingulate gyri. Smaller clusters were identified in the left pars opercularis and in the pericalcarine gyrus, as well as in the right caudal middle frontal and middle temporal gyri, the right pars triangularis, isthmus cingulate and rostral middle frontal gyri (see Figure S3; for detailed results and cluster size see SI, section 7, Table S3). Overall, MD decreases were observed in large occipito-parietal, temporal and frontal areas, suggesting tissue densification in response to task practice.

The surface-based analysis on NODDI parameters also disclosed a *Learning* (Pre $DWI_1$ vs. Post $DWI_2$) effect (CWP < 0.05) with increased NDI after learning in regions already exhibiting MD changes as described above. Bilaterally, NDI increased in the inferior parietal, posterior cingulate, inferior temporal and caudal middle frontal gyri, the pars opercularis, the lateral occipital gyrus, the precuneus, the banks of the superior temporal sulcus, the superior frontal, precentral paracentral, lingual, superior temporal, fusiform, supramarginal, medial orbitofrontal, rostral anterior cingulate, lateral orbito-frontal, superior parietal, rostral middle frontal, postcentral and caudal anterior cingulate gyri, the insula, the middle temporal gyrus, the pars triangularis, and the parahippocampal gyrus. Significant clusters were also identified in the right cuneus, the isthmus cingulate gyrus, the temporal pole and the pericalcarine gyrus. FWF decrease was evidenced in comparable areas exhibiting MD decrease, such as the bilateral postcentral gyrus, the precuneus, the superior parietal gyrus, the cuneus, the precentral, lateral occipital, paracentral, inferior parietal, superior frontal, lingual and posterior cingulate gyri, the insula and the supramarginal gyrus. Also, significant clusters were identified in the right superior temporal and fusiform gyri as well as in the left caudal anterior cingulate gyrus, the pars opercularis, the inferior temporal and isthmus cingulate gyri. Finally, ODI decreased in the right rostral middle frontal, superior frontal, and left posterior cingulate gyri clusters. Lastly, ODI increased in the right pericalcarine and fusiform gyri and the insula, and in the left postcentral, caudal middle frontal, isthmus cingulate and superior temporal gyri (see SI, section 7, Table S3). Altogether, the territories exhibiting large MD decreases in response to learning overlapped with NDI increases, FWF decreases and ODI changes going in both directions, indicating an increase in neurite density, accompanied by a decrease in tissue fraction and partial fibre reorganization by the end of the learning episode.

Lastly, correlation analyses did not reveal any significant between clusters showing microstructural changes over the learning session and online motor performance improvement.

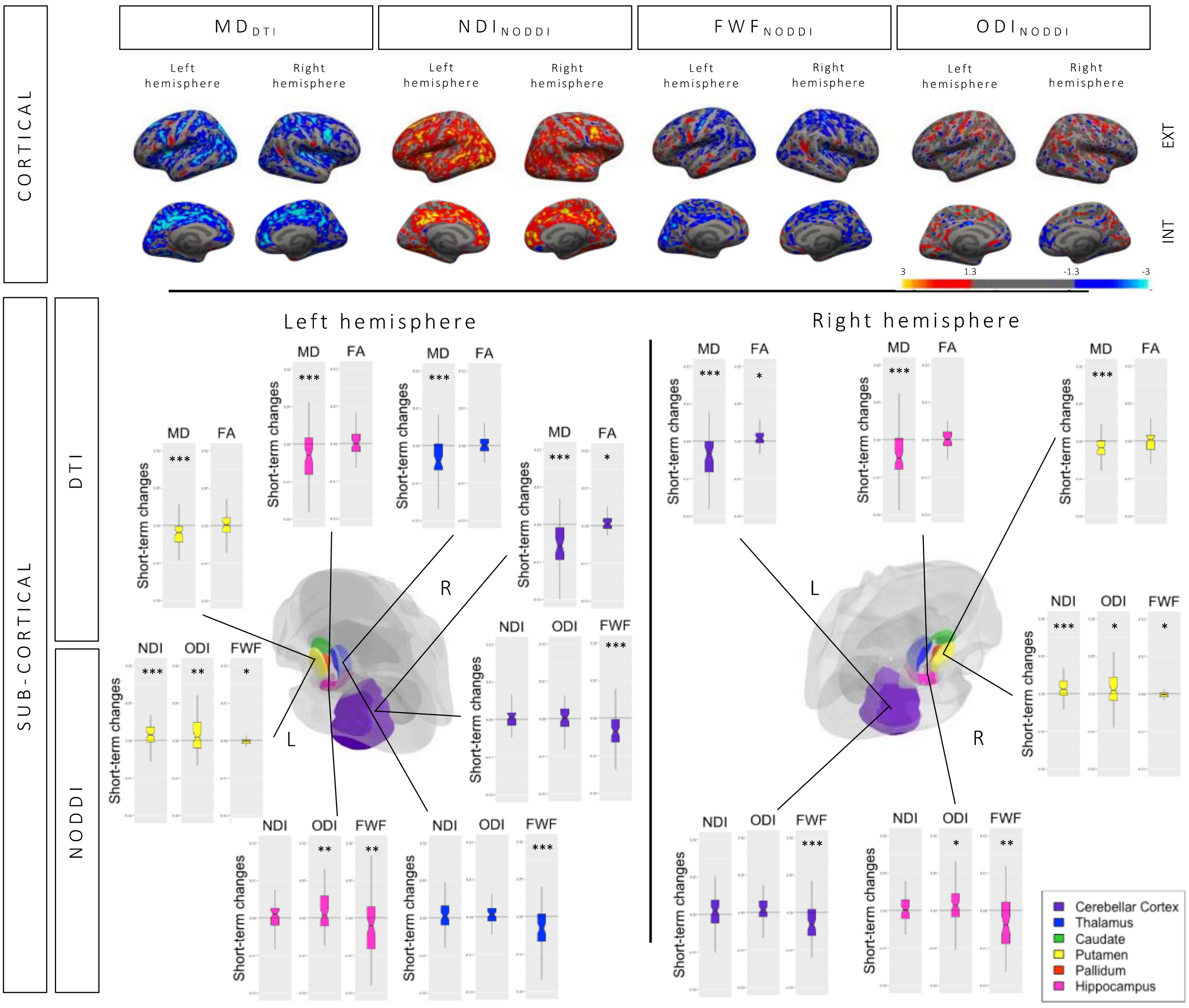


*Figure S3: Cortical and subcortical changes following the learning session ($DWI_2$-$DWI_1$)*

After comparing the post-learning scan with the baseline scan for the entire sample, noticeable changes in MD and NDI were observed across extensive occipitoparietal, frontal, and temporal regions. Additionally, changes in FWF and ODI were detected in similar regions in response to initial training. In the subcortical areas, alterations were evidenced in most of the predefined regions of interest (ROIs). No differences between *Conditions* were found at the cortical level, but a significant *Learning*Condition* interaction in the right caudate as detailed in section 3.4.1.2. DTI = Diffusion Tensor Imaging, MD = Mean Diffusivity, FA = Fractional Anisotropy, NODDI = Neurite Orientation Dispersion and Density Imaging, NDI = Neurite Density Index, FWF = Free Water Fraction, ODI = Orientation Dispersion Index. Colour-coded cortical images depict Z-scores (uncorrected threshold $p < 0.05$).

#### 6.1.2. Subcortical ROIs

MANOVAs with within-subject factor *Learning* (Pre $DWI_1$ vs. Post $DWI_2$) and between-subject factor *Condition* (TMR vs. RS) were conducted on DTI parameters for each ROI (see Figure S3). Significant main *Learning* effects were found on both sides of the cerebellar cortex, the putamen, the hippocampus and the left thalamus (all $p_{corr} < 0.001$). Post hoc analyses revealed a strong decrease in MD in all these regions (all *p*s < 0.001), suggesting increased tissue density after learning (as compared to baseline). Also, there was increased FA in the left ($p = 0.028$) and right cerebellar cortex ($p = 0.014$), reflecting increased directionality of diffusion inside of these regions.

Similar MANOVAs were performed on NODDI parameters in every ROI. Significant main *Learning* effects were observed bilaterally in the cerebellar cortex, the putamen, the hippocampus and the left thalamus (all $p_{corr} < 0.002$). Post-hoc analyses highlighted increased NDI in both parts of the putamen ($p < 0.001$), suggesting an increase in the number of neurites and increased ODI in the left ($p = 0.007$) and right putamen ($p = 0.045$) and in the left ($p = 0.008$) and right hippocampus ($p = 0.018$) suggesting increased dispersion of the fibres. Also, an important decrease in FWF was observed in all the previously mentioned ROIs such as both parts of the cerebellar cortex ($p < 0.001$), the left thalamus ($p < 0.001$), the left ($p = 0.044$) and right putamen ($p = 0.037$), and the left ($p = 0.002$) and right hippocampus ($p = 0.001$), showing a decrease in free water inside of the voxels.

ROIs displaying significant changes over the learning session did not correlate with online performance improvements (all *p*s > 0.090).

#### 6.2. Motor retraining-related structural changes and sleep-related effects (Day 5; DWI4 vs. DWI5 X RS vs. TMR)

##### 6.2.1. Cortical Ribbon

The surface-based statistical analysis on DTI parameters revealed a *Relearning* (Pre $DWI_4$ vs. Post $DWI_5$) effect (CWP < 0.05) for the whole sample, with a MD decrease bilaterally in the lateral occipital gyrus, the precuneus, the superior parietal, isthmus cingulate, precentral, fusiform, posterior cingulate, supra marginal, middle temporal gyri, the cuneus, the postcentral gyrus, the insula, the rostral middle frontal, superior frontal, lingual, caudal middle frontal, and caudal anterior cingulate gyri, as well as in the left inferior parietal and transverse temporal gyri and the banks of the right superior temporal sulcus, the right medial orbitofrontal, paracentral, lateral orbito frontal gyri, the pars opercularis and the pericalcarine gyrus (see Figure S4; for detailed results and cluster size see SI, section 7, Table S8). Hence, decreased MD after the relearning session displayed a pattern similar to the one observed after the learning episode, but with less amplitude.

The same analysis conducted on NODDI parameters revealed a *Relearning* (Pre $DWI_4$ vs. Post $DWI_5$) effect ($CWP < 0.05$) with NDI increases observed bilaterally in small clusters including the isthmus cingulate, rostral middle frontal, fusiform, precentral, caudal middle frontal, posterior cingulate, inferior parietal, middle temporal, parahippocampal, supramarginal, superior frontal and -parietal gyri, the insula, and the banks of the superior temporal sulcus. On the left hemisphere, the inferior temporal, transverse temporal, and postcentral gyri also expressed increased NDI and on the right hemisphere, clusters were found in the lateral occipital gyrus, the pars opercularis, the cuneus, precuneus, the medial and lateral-orbitofrontal, and the caudal anterior cingulate gyri. Decreased FWF was identified in bilateral cuneus, the insula, the superior parietal, postcentral, lateral occipital, pericalcarine gyri, the precuneus, the inferior parietal and supramarginal gyri as well as in the left precentral and transverse temporal gyri and the right isthmus cingulate, superior frontal gyri, the banks of the right superior temporal sulcus, and the right fusiform and entorhinal gyri. Small clusters showing ODI changes (both increase and decrease) were found in the bilateral inferior parietal and rostral middle frontal gyri, the precuneus, the banks of the superior temporal sulcus and the lateral occipital gyrus. On the left side, small clusters were also identified in the superior frontal, post- and precentral gyri, and on the right side, in the lingual gyrus, the pars triangularis, the supramarginal gyrus, the insula, the caudal middle frontal, the middle temporal, superior parietal, paracentral, medial orbitofrontal gyri. Overall, NDI increases were found in areas that overlapped the MD changes observed using DTI, indicating an increase in neurite density in response to the relearning session. FWF decreases and ODI changes going in both directions suggest a decrease in tissue proportion and fiber reorganization.

Clusters displaying significant changes did not correlate with online performance improvement during the relearning session. Also, no significant correlation was found with the different sleep parameters.

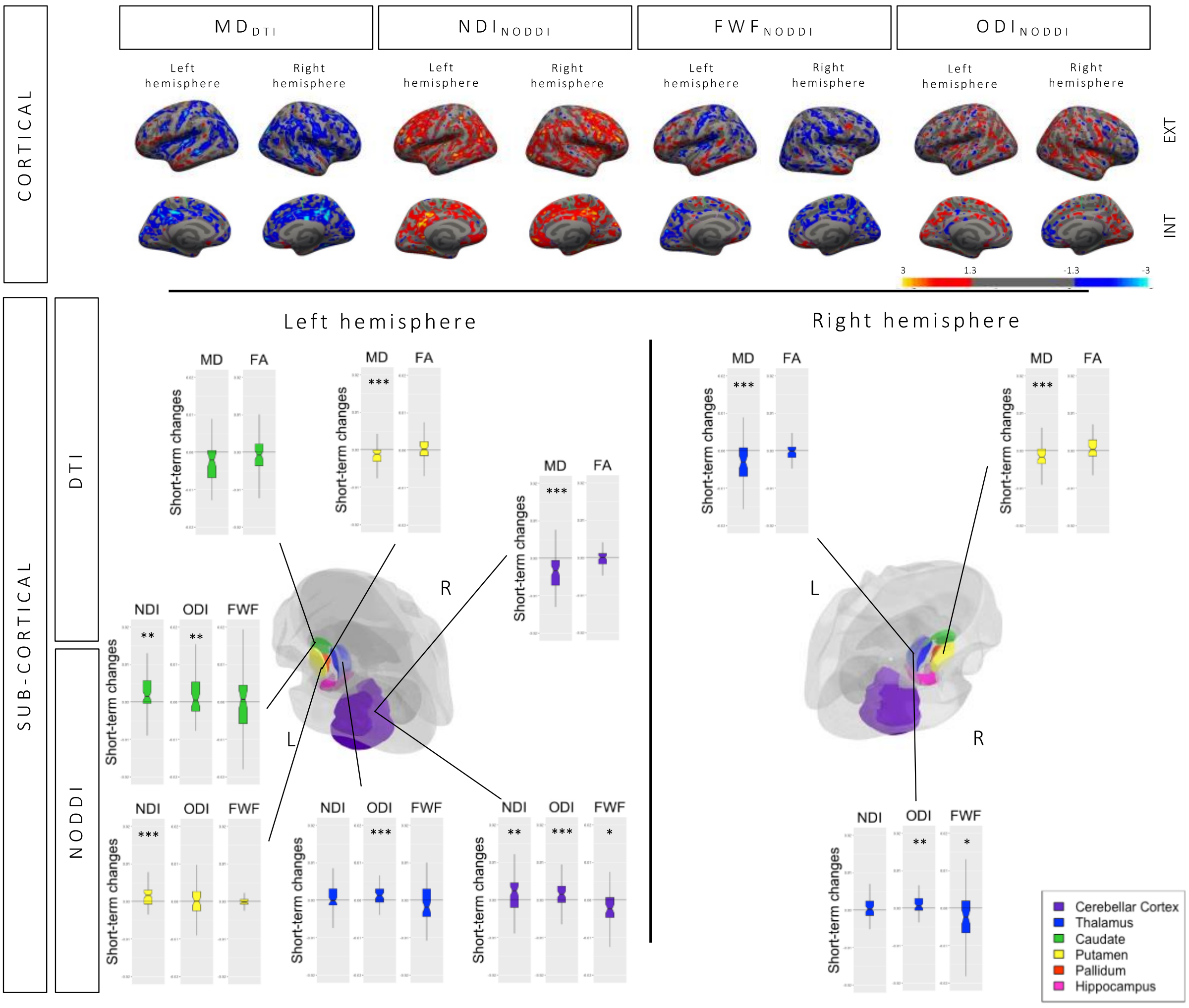


*Figure S4: Re-practice related cortical and subcortical modifications ($DWI_5$-$DWI_4$)*

After the relearning session, when comparing pre- and post-relearning scans, the entire sample exhibited structural changes across widespread cortical areas and in most predefined ROIs. Noticeably, the hippocampus did not exhibit structural changes as a result of the relearning session. However, the caudate nucleus, which was not implicated in the initial learning session, exhibited structural modifications. Cortical *Condition* differences are imaged in Figure 6. No *Relearning*Condition* interaction emerged in our ROIs as detailed in section 3.4.4.2. DTI = Diffusion Tensor Imaging, MD = Mean Diffusivity, FA = Fractional Anisotropy, NODDI = Neurite Orientation Dispersion and Density Imaging, NDI = Neurite Density Index, FWF = Free Water Fraction, ODI = Orientation Dispersion Index. Colour-coded cortical images depict Z-scores (uncorrected threshold $p < 0.05$).

#### 6.2.2. Subcortical ROIs

The MANOVA computed on DTI parameters with within-subject factor *ReLearning* (Pre $DWI_4$ vs. Post $DWI_5$) and between factor *Condition* (RS vs. TMR) highlighted a significant main *ReLearning* effect in the left ($p_{corr}$ = 0.002) and right putamen ($p_{corr}$ = 0.004), in the left cerebellar cortex ($p_{corr}$ < 0.001), the left caudate ($p_{corr}$ = 0.005) and in the right thalamus ($p_{corr}$ = 0.001). Post-hoc analyses showed that these changes were driven by decreased MD in all regions (all $p$s $< 0.001$) but the left caudate ($p = 0.838$), suggesting tissue densification (see Figure S4). No changes in FA were observed (all $p$s $> 0.059$).

The MANOVA computed on NODDI parameters disclosed a main *ReLearning* effect with changes in the left ($p_{corr} = 0.001$) and right thalamus ($p_{corr} = 0.005$), the left putamen ($p_{corr} = 0.003$), the left caudate ($p_{corr} = 0.001$) and the left cerebellar cortex ($p_{corr} = 0.001$). Post-hoc tests revealed decreased FWF in the right thalamus ($p = 0.012$) and the left cerebellar cortex ($p = 0.019$), increased NDI in the left cerebellar cortex ($p = 0.002$), the left caudate ($p = 0.005$) and the left putamen ($p < 0.001$), and increased ODI in the left cerebellar cortex ($p < 0.001$), the left thalamus ($p < 0.001$), the left caudate ($p = 0.006$) and the right thalamus ($p = 0.004$).

ROIs exhibiting significant diffusion changes did not significantly correlate with online performance improvement during the relearning session (all $p$s $> 0.107$). Regarding sleep parameters, FWF in the left cerebellar cortex positively correlated with NREM3 duration ($r = 0.308$; $p = 0.018$). No other regions exhibited significant correlations with sleep parameters (all $p$s $> 0.060$).

### 7. Tables

#### Table S1. Participant's demographics

| | Regular Sleep (RS) Group n = 30 | Targeted Memory Reactivation (TMR) Group n = 30 | Total sample n = 60 |
|---|---|---|---|
| Pittsburgh Sleep Quality Index (sleep quality) | mean score = 3.33<br>SD = 1.35<br>min = 0<br>max = 5 | mean score = 3.60<br>SD = 1.45<br>min = 1<br>max = 6 | mean score = 3.47<br>SD = 1.40<br>min = 0<br>max = 6 |
| Morningness–Eveningness Questionnaire (chronotype) | mean score = 53.417<br>SD = 7.557<br>min = 32<br>max = 68 | mean score = 55.13<br>SD = 6.63<br>min = 40<br>max = 68 | mean score = 54.28<br>SD = 7.10<br>min = 32<br>max = 68 |
| Edinburgh Inventory (laterality) | mean score = 4.27<br>SD = 6.64<br>min = -10<br>max = 10 | mean score = 5.43<br>SD = 6.61<br>min = -10<br>max = 10 | mean score = 4.85<br>SD = 6.60<br>min = −10<br>max = 10 |
| Gender | ♀ = 15<br>♂ = 15 | ♀ = 15<br>♂ = 15 | ♀ = 30<br>♂ = 30 |
| Age | mean score = 21.57<br>SD = 2.16<br>min = 18<br>max = 29 | mean score = 21.60<br>SD = 2.21<br>min = 18<br>max = 25 | mean age = 21.58<br>SD = 2.17<br>min = 18<br>max = 29 |

*Table S1.* Detailed information concerning sleep quality, chronotype, laterality, gender, and age with mean (mean score), standard deviation (SD), minimum (min) and maximum (max) for both groups separated and in the total sample.

#### Table S2. Sleep duration by stage

| | | Regular Sleep (RS) Group n = 30 | Targeted Memory Reactivation (TMR) Group n = 30 | Total sample n = 60 |
|---|---|---|---|---|
| N1 | Night 1 | mean = 1649 sec<br>SD = 760.763 sec<br>min = 390 sec<br>max = 3660 sec | mean = 1541 sec<br>SD = 577.840 sec<br>min = 630 sec<br>max = 3120 sec | mean = 1595 sec<br>SD = 671.983 sec<br>min = 390 sec<br>max = 3660 sec |
| | Night 2 | mean = 1260 sec<br>SD = 697.315 sec<br>min = 360 sec<br>max = 3810 sec | mean = 1240 sec<br>SD = 451.297 sec<br>min = 600 sec<br>max = 2370 sec | mean = 1250 sec<br>SD = 582.420 sec<br>min = 360 sec<br>max = 3810 sec |
| N2 | Night 1 | mean = 14618 sec<br>SD = 2196.586 sec<br>min = 9030 sec<br>max = 18960 sec | mean = 15141 sec<br>SD = 2390.531 sec<br>min = 9960 sec<br>max = 20730 sec | mean = 14879.5 sec<br>SD = 2291.296 sec<br>min = 9030 sec<br>max = 20730 sec |

| | | | | |
|---|---|---|---|---|
| | Night 2 | mean = 14379 sec<br>SD = 2333.707 sec<br>min = 10170 sec<br>max = 18630 sec | mean = 15154 sec<br>SD = 1783.593 sec<br>min = 11100 sec<br>max = 18510 sec | mean = 14766.5 sec<br>SD = 2096.016 sec<br>min = 10170 sec<br>max = 18630 sec |
| SWS | Night 1 | mean = 4908 sec<br>SD = 1441.736 sec<br>min = 1230 sec<br>max = 9660 sec | mean = 4423 sec<br>SD = 1378.260 sec<br>min = 1920 sec<br>max = 7260 sec | mean = 4665.5 sec<br>SD = 1419.574 sec<br>min = 1230 sec<br>max = 9660 sec |
| | Night 2 | mean = 5148 sec<br>SD = 1353.125 sec<br>min = 2130 sec<br>max = 8670 sec | mean = 4144 sec<br>SD = 1202.473 sec<br>min = 1350 sec<br>max = 6240 sec | mean = 4646 sec<br>SD = 1366.364 sec<br>min = 1350 sec<br>max = 8670 sec |
| REM | Night 1 | mean = 5381 sec<br>SD = 1893.101 sec<br>min = 150 sec<br>max = 9780 sec | mean = 5298 sec<br>SD = 1598.947 sec<br>min = 0 sec<br>max = 7590 sec | mean = 5335 sec<br>SD = 1737.914 sec<br>min = 0 sec<br>max = 9780 sec |
| | Night 2 | mean = 6083 sec<br>SD = 1311.075 sec<br>min = 3000 sec<br>max = 8520 sec | mean = 6661 sec<br>SD = 1217.344 sec<br>min = 4200 sec<br>max = 9510 sec | mean = 6372 sec<br>SD = 1287.724 sec<br>min = 3000 sec<br>max = 9510 sec |

*Table S2.* Detailed information concerning sleep duration for each stage separately during the habituation night and the experimental night with mean (mean duration expressed in **seconds**), standard deviation (SD), minimum (min) and maximum (max) for both groups separated and in the total sample.

### Table S3. Learning-related short-term structural changes (Day 1; DWI1 vs. DWI2) in the cortical ribbon

| **LEARNING $DWI_2$-$DWI_1$ for the whole sample (group RS + TMR)** | | | | |
|---|---|---|---|---|
| **Annot** | **Size(mm^2)** | **WghtVtx** | **NVtxs** | **nbClust** |
| **MD – right hemisphere** | | | | |
| paracentral | 764.01 | -7590.99 | 1876.00 | 4.00 |
| precentral | 583.39 | -6142.29 | 1534.00 | 3.00 |
| postcentral | 537.91 | -5348.23 | 1326.00 | 6.00 |
| caudalmiddlefrontal | 522.09 | -4920.94 | 1089.00 | 2.00 |
| precuneus | 403.21 | -4606.76 | 1100.00 | 3.00 |
| superiorfrontal | 366.17 | -2840.36 | 726.00 | 5.00 |
| inferiorparietal | 293.57 | -2612.39 | 618.00 | 4.00 |
| superiorparietal | 270.47 | -2467.50 | 669.00 | 5.00 |
| rostralanteriorcingulate | 258.96 | -1873.39 | 479.00 | 1.00 |
| superiortemporal | 251.48 | -2360.42 | 604.00 | 4.00 |
| fusiform | 236.16 | -1624.56 | 389.00 | 4.00 |
| caudalanteriorcingulate | 185.54 | -1844.58 | 458.00 | 1.00 |
| lateraloccipital | 180.36 | -889.10 | 256.00 | 5.00 |
| cuneus | 176.71 | -786.03 | 197.00 | 2.00 |
| transversetemporal | 161.60 | -2242.93 | 497.00 | 1.00 |

| | | | | |
|---|---|---|---|---|
| middletemporal | 113.44 | -857.03 | 237.00 | 2.00 |
| lateralorbitofrontal | 112.46 | -784.47 | 222.00 | 1.00 |
| insula | 101.71 | -994.59 | 260.00 | 3.00 |
| parstriangularis | 90.72 | -853.80 | 237.00 | 1.00 |
| supramarginal | 72.71 | -667.28 | 181.00 | 1.00 |
| lingual | 57.30 | -305.87 | 85.00 | 2.00 |
| isthmuscingulate | 27.28 | -250.67 | 75.00 | 3.00 |
| posteriorcingulate | 18.61 | -146.91 | 43.00 | 1.00 |
| bankssts | 10.47 | -94.75 | 30.00 | 1.00 |
| rostralmiddlefrontal | 4.36 | -24.72 | 8.00 | 1.00 |
| inferiortemporal | 4.35 | -21.73 | 7.00 | 1.00 |
| **MD – left hemisphere** | | | | |
| precuneus | 1490.33 | -15150.69 | 3321.00 | 3.00 |
| superiorparietal | 1221.72 | -11312.58 | 2457.00 | 7.00 |
| lateraloccipital | 972.75 | -4783.65 | 1278.00 | 9.00 |
| precentral | 575.56 | -5855.13 | 1439.00 | 6.00 |
| insula | 471.53 | -5521.23 | 1334.00 | 4.00 |
| superiorfrontal | 411.46 | -3339.73 | 802.00 | 6.00 |
| inferiorparietal | 396.07 | -3424.30 | 830.00 | 4.00 |
| postcentral | 394.59 | -3645.50 | 987.00 | 4.00 |
| supramarginal | 392.23 | -3801.17 | 1030.00 | 8.00 |
| inferiortemporal | 381.26 | -2824.09 | 662.00 | 3.00 |
| bankssts | 375.63 | -3513.14 | 885.00 | 3.00 |
| paracentral | 280.90 | -3031.27 | 784.00 | 1.00 |
| lingual | 269.79 | -1820.17 | 456.00 | 3.00 |
| caudalanteriorcingulate | 189.44 | -1699.39 | 448.00 | 1.00 |
| cuneus | 124.36 | -684.44 | 174.00 | 2.00 |
| rostralanteriorcingulate | 115.95 | -895.43 | 234.00 | 2.00 |
| fusiform | 104.21 | -890.58 | 217.00 | 2.00 |
| lateralorbitofrontal | 63.68 | -426.12 | 115.00 | 1.00 |
| superiortemporal | 58.91 | -526.34 | 149.00 | 4.00 |
| posteriorcingulate | 56.74 | -780.18 | 230.00 | 2.00 |
| transversetemporal | 50.77 | -559.52 | 145.00 | 1.00 |
| parsopercularis | 8.87 | -60.39 | 19.00 | 2.00 |
| pericalcarine | 5.65 | -30.69 | 10.00 | 1.00 |
| **NDI – right hemisphere** | | | | |
| precuneus | 431.42 | 4506.57 | 1067.00 | 4.00 |
| caudalmiddlefrontal | 428.72 | 3551.49 | 905.00 | 1.00 |
| precentral | 313.49 | 3641.10 | 888.00 | 6.00 |
| superiorfrontal | 299.55 | 1974.49 | 526.00 | 7.00 |
| paracentral | 271.90 | 2976.81 | 698.00 | 2.00 |
| superiortemporal | 251.91 | 1806.67 | 571.00 | 4.00 |
| inferiorparietal | 155.28 | 963.73 | 286.00 | 8.00 |
| lateraloccipital | 144.92 | 768.16 | 219.00 | 3.00 |
| medialorbitofrontal | 144.58 | 788.99 | 213.00 | 1.00 |
| rostralanteriorcingulate | 132.23 | 852.16 | 239.00 | 1.00 |

| | | | | |
|---|---|---|---|---|
| lateralorbitofrontal | 127.58 | 943.25 | 242.00 | 2.00 |
| superiorparietal | 117.69 | 1120.88 | 291.00 | 3.00 |
| postcentral | 94.29 | 957.01 | 252.00 | 4.00 |
| supramarginal | 82.28 | 725.54 | 213.00 | 7.00 |
| insula | 80.59 | 804.94 | 236.00 | 3.00 |
| middletemporal | 76.86 | 497.67 | 153.00 | 5.00 |
| caudalanteriorcingulate | 67.77 | 576.42 | 154.00 | 1.00 |
| fusiform | 60.03 | 429.68 | 101.00 | 2.00 |
| cuneus | 59.85 | 320.39 | 89.00 | 1.00 |
| lingual | 55.94 | 213.75 | 63.00 | 2.00 |
| isthmuscingulate | 50.82 | 461.49 | 139.00 | 2.00 |
| posteriorcingulate | 47.98 | 376.77 | 108.00 | 2.00 |
| inferiortemporal | 46.51 | 249.67 | 73.00 | 3.00 |
| rostralmiddlefrontal | 42.67 | 273.82 | 78.00 | 2.00 |
| parsopercularis | 37.10 | 308.80 | 79.00 | 2.00 |
| temporalpole | 29.31 | 164.87 | 48.00 | 1.00 |
| parstriangularis | 24.40 | 242.14 | 68.00 | 1.00 |
| parahippocampal | 15.99 | 94.29 | 30.00 | 2.00 |
| bankssts | 14.84 | 165.75 | 49.00 | 1.00 |
| pericalcarine | 4.03 | 18.40 | 6.00 | 1.00 |
| **NDI – left hemisphere** | | | | |
| inferiorparietal | 1044.55 | 8810.76 | 2151.00 | 11.00 |
| posteriorcingulate | 804.74 | 7565.56 | 1792.00 | 3.00 |
| inferiortemporal | 687.35 | 4648.96 | 1151.00 | 1.00 |
| caudalmiddlefrontal | 621.72 | 5807.70 | 1311.00 | 2.00 |
| parsopercularis | 586.90 | 5594.45 | 1387.00 | 5.00 |
| lateraloccipital | 586.75 | 2961.15 | 787.00 | 6.00 |
| bankssts | 360.80 | 3366.64 | 860.00 | 2.00 |
| superiorfrontal | 344.12 | 2284.24 | 714.00 | 7.00 |
| lingual | 252.68 | 1848.87 | 451.00 | 5.00 |
| fusiform | 198.23 | 1242.76 | 335.00 | 3.00 |
| supramarginal | 189.85 | 1780.92 | 486.00 | 5.00 |
| rostralanteriorcingulate | 137.87 | 1001.04 | 271.00 | 2.00 |
| lateralorbitofrontal | 109.94 | 828.95 | 205.00 | 2.00 |
| rostralmiddlefrontal | 99.86 | 618.85 | 184.00 | 4.00 |
| superiorparietal | 89.71 | 770.35 | 221.00 | 3.00 |
| caudalanteriorcingulate | 81.28 | 666.87 | 184.00 | 1.00 |
| insula | 80.80 | 813.82 | 235.00 | 4.00 |
| postcentral | 71.82 | 615.73 | 179.00 | 5.00 |
| superiortemporal | 70.36 | 519.34 | 155.00 | 3.00 |
| precentral | 69.73 | 524.01 | 159.00 | 3.00 |
| parstriangularis | 69.23 | 428.83 | 118.00 | 2.00 |
| middletemporal | 48.29 | 231.52 | 70.00 | 3.00 |
| medialorbitofrontal | 44.57 | 179.77 | 49.00 | 1.00 |
| paracentral | 37.48 | 409.64 | 113.00 | 2.00 |
| precuneus | 26.98 | 183.37 | 57.00 | 2.00 |

| | | | | |
|---|---|---|---|---|
| parahippocampal | 9.35 | 56.98 | 18.00 | 1.00 |
| **FWF – right hemisphere** | | | | |
| precuneus | 242.91 | -1357.34 | 334.00 | 2.00 |
| precentral | 104.87 | -896.37 | 265.00 | 2.00 |
| superiorparietal | 87.10 | -567.19 | 171.00 | 6.00 |
| postcentral | 77.45 | -467.06 | 139.00 | 3.00 |
| paracentral | 73.55 | -520.74 | 149.00 | 2.00 |
| superiortemporal | 61.99 | -530.84 | 140.00 | 1.00 |
| inferiorparietal | 32.52 | -291.92 | 75.00 | 1.00 |
| superiorfrontal | 31.56 | -185.81 | 57.00 | 2.00 |
| cuneus | 27.33 | -136.75 | 36.00 | 1.00 |
| posteriorcingulate | 24.03 | -203.29 | 58.00 | 1.00 |
| insula | 19.22 | -147.49 | 45.00 | 3.00 |
| lingual | 16.56 | -76.14 | 24.00 | 2.00 |
| supramarginal | 11.42 | -105.04 | 33.00 | 1.00 |
| fusiform | 5.82 | -27.80 | 9.00 | 1.00 |
| lateraloccipital | 3.69 | -15.20 | 5.00 | 1.00 |
| **FWF – left hemisphere** | | | | |
| postcentral | 244.72 | -1932.36 | 528.00 | 3.00 |
| superiorparietal | 213.54 | -1444.18 | 397.00 | 5.00 |
| cuneus | 157.58 | -896.33 | 224.00 | 2.00 |
| precuneus | 145.52 | -1111.52 | 312.00 | 4.00 |
| precentral | 137.99 | -1425.26 | 401.00 | 2.00 |
| lateraloccipital | 134.00 | -618.83 | 183.00 | 5.00 |
| inferiorparietal | 49.61 | -269.39 | 74.00 | 2.00 |
| caudalanteriorcingulate | 34.67 | -231.54 | 71.00 | 2.00 |
| lingual | 26.49 | -150.28 | 47.00 | 2.00 |
| insula | 21.48 | -261.82 | 75.00 | 2.00 |
| paracentral | 17.11 | -166.70 | 50.00 | 2.00 |
| parsopercularis | 10.43 | -80.61 | 25.00 | 1.00 |
| superiorfrontal | 10.40 | -79.58 | 24.00 | 1.00 |
| posteriorcingulate | 9.50 | -94.50 | 30.00 | 2.00 |
| inferiortemporal | 3.54 | -21.50 | 7.00 | 1.00 |
| isthmuscingulate | 1.29 | -9.05 | 3.00 | 2.00 |
| supramarginal | 0.29 | -3.08 | 1.00 | 1.00 |
| **ODI – right hemisphere** | | | | |
| rostralmiddlefrontal | 123.44 | -850.65 | 198.00 | 1.00 |
| pericalcarine | 9.55 | 34.81 | 11.00 | 1.00 |
| superiorfrontal | 7.50 | -31.06 | 10.00 | 1.00 |
| fusiform | 4.86 | 5.83 | 8.00 | 2.00 |
| insula | 0.89 | 6.11 | 2.00 | 1.00 |
| **ODI – left hemisphere** | | | | |
| postcentral | 39.74 | 264.14 | 68 | 1 |
| posteriorcingulate | 19.27 | -189.76 | 60 | 1 |
| caudalmiddlefrontal | 11.26 | 52.46 | 16 | 2 |
| isthmuscingulate | 5.80 | 79.07 | 22 | 1 |

| | | | | |
|---|---|---|---|---|
| superiortemporal | 3.29 | 21.47 | 7 | 1 |

*Table S3.* Clusters showing significant MD, NDI, FWF and ODI changes in both groups following learning at Day 1 after cluster-wise correction gathered by cortical region. The table represent changes for both groups. Differences between Conditions were also investigated but no significant difference emerged. Annot = the name of the regions the maximum cluster value falls into using **Desikan-Killiany atlas.** Size (mm$^2$) = sum of the cluster sizes in millimeters square. WghtVtx = sum of the cluster weights (size*intensity), a negative weight indicate decrease in intensity while a positive value points towards an increase in intensity. NVtxs = sum of the number of vertices in the clusters from the region. NbClust = number of clusters in the labelled area.

### Table S4. Delayed structural changes in Pre-learning vs. Morning scan (Day 1 DWI1 vs. Day 2 DWI3) in the cortical ribbon

| **POST-SLEEP MORNING DWI$_3$-DWI$_1$ for the whole sample (group RS + TMR)** | | | | |
|---|---|---|---|---|
| **Annot** | **Size(mm^2)** | **WghtVtx** | **NVtxs** | **nbClust** |
| **MD – right hemisphere** | | | | |
| superiorparietal | 413.75 | -3293.03 | 886.00 | 4.00 |
| inferiorparietal | 240.97 | -1800.06 | 494.00 | 6.00 |
| lateraloccipital | 186.20 | -1081.04 | 272.00 | 4.00 |
| precuneus | 129.97 | -1109.15 | 326.00 | 5.00 |
| cuneus | 77.82 | -362.80 | 96.00 | 1.00 |
| superiorfrontal | 69.92 | -419.43 | 130.00 | 7.00 |
| precentral | 63.20 | -630.30 | 174.00 | 2.00 |
| inferiortemporal | 50.64 | -197.67 | 59.00 | 1.00 |
| supramarginal | 43.94 | -312.06 | 96.00 | 6.00 |
| fusiform | 26.64 | -143.49 | 42.00 | 1.00 |
| postcentral | 20.29 | -209.29 | 58.00 | 1.00 |
| caudalanteriorcingulate | 17.21 | -149.11 | 45.00 | 1.00 |
| isthmuscingulate | 15.64 | -162.31 | 49.00 | 2.00 |
| paracentral | 14.70 | -160.59 | 48.00 | 2.00 |
| rostralmiddlefrontal | 11.31 | -63.08 | 20.00 | 1.00 |
| parsopercularis | 10.99 | -105.90 | 29.00 | 1.00 |
| parsorbitalis | 10.00 | 69.91 | 21.00 | 1.00 |
| bankssts | 7.33 | -60.61 | 19.00 | 3.00 |
| superiortemporal | 3.59 | -24.72 | 8.00 | 1.00 |
| lateralorbitofrontal | 1.64 | 9.06 | 3.00 | 1.00 |
| **MD – left hemisphere** | | | | |
| superiorparietal | 444.08 | -3465.46 | 900.00 | 4.00 |
| inferiorparietal | 442.13 | -3358.04 | 914.00 | 5.00 |
| precuneus | 301.19 | -2141.44 | 631.00 | 10.00 |
| lateraloccipital | 264.53 | -1422.44 | 365.00 | 3.00 |
| supramarginal | 224.65 | -2494.21 | 621.00 | 3.00 |
| superiorfrontal | 96.76 | -631.55 | 189.00 | 6.00 |
| posteriorcingulate | 76.77 | -726.24 | 213.00 | 3.00 |
| isthmuscingulate | 66.36 | -514.63 | 134.00 | 1.00 |

| | | | | |
|---|---|---|---|---|
| cuneus | 30.09 | -117.21 | 36.00 | 1.00 |
| lingual | 25.15 | -141.70 | 42.00 | 1.00 |
| fusiform | 18.16 | -95.88 | 31.00 | 2.00 |
| caudalanteriorcingulate | 12.40 | -95.09 | 29.00 | 1.00 |
| pericalcarine | 9.26 | -27.94 | 9.00 | 1.00 |
| middletemporal | 8.10 | 34.22 | 11.00 | 1.00 |
| precentral | 5.77 | -36.67 | 12.00 | 1.00 |
| postcentral | 3.25 | -18.44 | 6.00 | 1.00 |
| **NDI – right hemisphere** | | | | |
| inferiorparietal | 964.64 | 6797.67 | 1838.00 | 12.00 |
| precentral | 730.26 | 6161.18 | 1615.00 | 7.00 |
| superiorfrontal | 488.18 | 3186.14 | 792.00 | 7.00 |
| supramarginal | 290.82 | 2188.63 | 600.00 | 11.00 |
| caudalanteriorcingulate | 254.54 | 2137.56 | 606.00 | 4.00 |
| posteriorcingulate | 215.34 | 2179.82 | 528.00 | 4.00 |
| precuneus | 197.56 | 1466.87 | 410.00 | 9.00 |
| parsopercularis | 196.74 | 1390.88 | 298.00 | 2.00 |
| superiorparietal | 185.88 | 1675.96 | 422.00 | 1.00 |
| isthmuscingulate | 172.78 | 1610.56 | 448.00 | 3.00 |
| rostralmiddlefrontal | 161.33 | 914.55 | 269.00 | 4.00 |
| lateraloccipital | 159.75 | 877.93 | 238.00 | 4.00 |
| bankssts | 154.99 | 1333.61 | 368.00 | 2.00 |
| insula | 72.62 | 725.23 | 195.00 | 4.00 |
| medialorbitofrontal | 63.74 | 285.69 | 85.00 | 1.00 |
| parstriangularis | 61.43 | 327.19 | 95.00 | 2.00 |
| cuneus | 57.28 | 339.54 | 87.00 | 1.00 |
| paracentral | 50.10 | 520.60 | 147.00 | 3.00 |
| parsorbitalis | 49.65 | 220.80 | 62.00 | 1.00 |
| superiortemporal | 43.74 | 334.49 | 92.00 | 1.00 |
| postcentral | 29.10 | 265.72 | 83.00 | 4.00 |
| pericalcarine | 25.90 | 135.12 | 41.00 | 2.00 |
| transversetemporal | 15.51 | 141.14 | 44.00 | 1.00 |
| caudalmiddlefrontal | 4.45 | 34.07 | 11.00 | 1.00 |
| **NDI – left hemisphere** | | | | |
| superiorfrontal | 876.35 | 6079.51 | 1586.00 | 15.00 |
| superiortemporal | 854.65 | 7969.60 | 1924.00 | 3.00 |
| lateraloccipital | 666.34 | 3217.96 | 872.00 | 6.00 |
| inferiorparietal | 647.34 | 5020.45 | 1287.00 | 8.00 |
| isthmuscingulate | 405.55 | 3756.42 | 973.00 | 1.00 |
| supramarginal | 378.73 | 4021.35 | 1079.00 | 4.00 |
| superiorparietal | 296.18 | 2436.28 | 682.00 | 5.00 |
| caudalmiddlefrontal | 252.16 | 1748.07 | 482.00 | 5.00 |
| inferiortemporal | 171.26 | 1047.18 | 273.00 | 2.00 |
| insula | 168.25 | 1560.04 | 433.00 | 8.00 |
| parsopercularis | 163.57 | 1091.01 | 283.00 | 1.00 |
| precuneus | 156.34 | 1207.10 | 325.00 | 4.00 |

| | | | | |
|---|---|---|---|---|
| rostralmiddlefrontal | 150.32 | 805.11 | 238.00 | 4.00 |
| parstriangularis | 147.07 | 1116.35 | 303.00 | 2.00 |
| precentral | 137.11 | 1300.07 | 353.00 | 6.00 |
| postcentral | 127.58 | 1003.51 | 291.00 | 2.00 |
| lingual | 120.95 | 624.13 | 189.00 | 4.00 |
| posteriorcingulate | 103.79 | 1074.22 | 294.00 | 2.00 |
| fusiform | 70.04 | 289.18 | 90.00 | 4.00 |
| cuneus | 48.47 | 226.84 | 68.00 | 2.00 |
| rostralanteriorcingulate | 43.22 | 284.23 | 81.00 | 2.00 |
| middletemporal | 41.80 | 207.35 | 65.00 | 3.00 |
| bankssts | 31.93 | 241.85 | 73.00 | 2.00 |
| parahippocampal | 5.78 | 33.96 | 11.00 | 1.00 |
| pericalcarine | 3.13 | 18.31 | 6.00 | 1.00 |
| **FWF – right hemisphere** | | | | |
| lateralorbitofrontal | 106.74 | 630.61 | 159.00 | 3.00 |
| superiorparietal | 44.32 | -334.30 | 100.00 | 3.00 |
| inferiortemporal | 30.47 | 15.37 | 43.00 | 2.00 |
| superiortemporal | 23.23 | 187.64 | 57.00 | 3.00 |
| medialorbitofrontal | 15.64 | 90.45 | 29.00 | 1.00 |
| insula | 14.34 | 135.36 | 40.00 | 2.00 |
| cuneus | 11.31 | -43.34 | 14.00 | 1.00 |
| middletemporal | 4.23 | 31.23 | 10.00 | 1.00 |
| **FWF – left hemisphere** | | | | |
| lateralorbitofrontal | 177.89 | 1243.15 | 307 | 2 |
| medialorbitofrontal | 68.46 | 418.81 | 115 | 1 |
| middletemporal | 38.26 | 222.98 | 65 | 2 |
| lateraloccipital | 34.05 | -148.62 | 46 | 2 |
| superiortemporal | 31.31 | 265.38 | 74 | 2 |
| parstriangularis | 23.84 | 223.04 | 59 | 1 |
| superiorfrontal | 16.43 | 75.48 | 23 | 1 |
| bankssts | 15.36 | 155.86 | 44 | 1 |
| postcentral | 11.02 | -106.47 | 32 | 1 |
| lingual | 9.58 | -51.44 | 16 | 1 |
| paracentral | 8.88 | -90.66 | 25 | 1 |
| precuneus | 8.60 | -68.34 | 22 | 1 |
| parahippocampal | 7.71 | 48.91 | 15 | 1 |
| inferiorparietal | 5.03 | -31.20 | 10 | 1 |
| insula | 4.51 | 27.95 | 9 | 1 |
| superiorparietal | 4.17 | -25.02 | 8 | 1 |
| **ODI – right hemisphere** | | | | |
| superiortemporal | 81.19 | 501.56 | 118.00 | 1.00 |
| fusiform | 58.02 | 288.00 | 76.00 | 1.00 |
| lateralorbitofrontal | 49.23 | -194.29 | 80.00 | 2.00 |
| inferiorparietal | 40.90 | 303.78 | 89.00 | 4.00 |
| lingual | 40.62 | 172.43 | 51.00 | 3.00 |
| lateraloccipital | 31.47 | 126.42 | 37.00 | 1.00 |

| | | | | |
|---|---|---|---|---|
| inferiortemporal | 31.20 | 128.83 | 40.00 | 1.00 |
| insula | 29.80 | 298.48 | 80.00 | 1.00 |
| supramarginal | 22.39 | 142.95 | 42.00 | 2.00 |
| caudalmiddlefrontal | 18.21 | 87.74 | 26.00 | 1.00 |
| superiorfrontal | 17.97 | 38.30 | 31.00 | 2.00 |
| rostralmiddlefrontal | 16.89 | -81.51 | 26.00 | 2.00 |
| parahippocampal | 9.75 | 57.09 | 17.00 | 1.00 |
| superiorparietal | 4.17 | 25.34 | 8.00 | 1.00 |
| precentral | 0.85 | 6.11 | 2.00 | 1.00 |
| **ODI – left hemisphere** | | | | |
| superiorfrontal | 32.46 | 235.10 | 63.00 | 2.00 |
| superiorparietal | 21.19 | 108.00 | 34.00 | 2.00 |
| precuneus | 17.37 | 104.25 | 31.00 | 1.00 |
| rostralanteriorcingulate | 17.01 | -112.35 | 33.00 | 1.00 |
| inferiortemporal | 16.21 | 78.55 | 24.00 | 1.00 |
| lateraloccipital | 10.38 | 39.45 | 12.00 | 1.00 |
| caudalmiddlefrontal | 10.28 | 53.23 | 16.00 | 1.00 |
| supramarginal | 7.53 | 67.03 | 21.00 | 1.00 |
| fusiform | 6.64 | 29.12 | 9.00 | 1.00 |
| rostralmiddlefrontal | 3.95 | -15.46 | 5.00 | 1.00 |
| parstriangularis | 2.69 | -21.39 | 7.00 | 1.00 |
| lateralorbitofrontal | 2.07 | -12.31 | 4.00 | 1.00 |
| postcentral | 1.41 | 12.07 | 4.00 | 1.00 |
| precentral | 0.84 | 6.07 | 2.00 | 1.00 |

*Table S4.* Clusters exhibiting significant MD, NDI, FWF and ODI modifications at Day 2 when compared to baseline at Day 1 after cluster-wise correction brought together by region. The table represent changes for both groups. Annot = the name of the regions the maximum cluster value falls into using **Desikan-Killiany atlas**. Size ($mm^2$) = sum of the cluster sizes in millimeters square. WghtVtx = sum of the cluster weights (size*intensity), a negative weight indicate decrease in intensity while a positive value points towards an increase in intensity. NVtxs = sum of the number of vertices in the clusters from the region. NbClust = number of clusters in the labelled area.

### Table S5. Delayed structural changes and sleep-related effects in Pre-learning vs. Morning scan (Day 1 DWI1 vs. Day 2 DWI3 x RS vs. TMR) in the cortical ribbon

| **POST-SLEEP MORNING $DWI_3$-$DWI_1$; group differences (group TMR vs. RS)** | | | | |
|---|---|---|---|---|
| **Annot** | **Size(mm^2)** | **WghtVtx** | **NVtxs** | **nbClust** |
| **MD – right hemisphere** | | | | |
| rostralmiddlefrontal | 114.72 | 578.80 | 171 | 1 |
| **FWF – right hemisphere** | | | | |
| rostralmiddlefrontal | 97.10 | 469.56 | 142 | 1 |

*Table S5.* Clusters in the right rostral middle frontal gyrus exhibiting a significant difference in MD and FWF between both Conditions when comparing the morning scan on Day 2 to baseline at Day 1 after cluster-wise correction. Annot = the name of the regions the maximum cluster value falls into using **Desikan-Killiany atlas**. Size ($mm^2$) = sum of the cluster sizes in millimeters square. WghtVtx = sum of the cluster weights (size*intensity), a negative weight indicate decrease in intensity while a positive value points towards an increase in intensity. NVtxs = sum of the number of vertices in the clusters from the region. NbClust = number of clusters in the labelled area.

### Table S6. Delayed structural changes in Pre-learning vs. Pre-relearning (Day 1 DWI1 vs. Day 5 DWI4) in the cortical ribbon

| **POST-SLEEP DELAYED $DWI_4$-$DWI_1$ for the whole sample (group RS + TMR)** | | | | |
|---|---|---|---|---|
| **Annot** | **Size(mm^2)** | **WghtVtx** | **NVtxs** | **nbClust** |
| **MD – right hemisphere** | | | | |
| inferiortemporal | 10.32 | 52.89 | 17 | 1 |
| fusiform | 0.55 | -3.06 | 1 | 1 |
| precentral | 0.36 | -3.01 | 1 | 1 |
| **MD – left hemisphere** | | | | |
| lateraloccipital | 16.37 | -72.36 | 23 | 1 |
| caudalanteriorcingulate | 4.69 | -33.92 | 11 | 1 |
| rostralanteriorcingulate | 0.46 | -3.03 | 1 | 1 |
| **NDI – right hemisphere** | | | | |
| supramarginal | 11.72 | 85.12 | 26 | 1 |
| superiortemporal | 8.09 | 62.91 | 20 | 2 |
| paracentral | 4.29 | 46.24 | 15 | 1 |
| isthmuscingulate | 3.15 | -28.08 | 9 | 1 |
| insula | 1.21 | 12.31 | 4 | 1 |
| superiorfrontal | 1.02 | 6.07 | 2 | 1 |
| **NDI – left hemisphere** | | | | |
| postcentral | 7.19 | 55.57 | 18.00 | 2.00 |
| supramarginal | 6.81 | 52.08 | 17.00 | 2.00 |
| superiortemporal | 3.67 | 21.82 | 7.00 | 1.00 |
| precentral | 3.06 | 21.77 | 7.00 | 1.00 |
| superiorparietal | 1.76 | 12.08 | 4.00 | 1.00 |
| insula | 1.43 | 12.32 | 4.00 | 1.00 |
| **FWF – right hemisphere** | | | | |
| insula | 7.17 | 81.77 | 26 | 2 |
| lateraloccipital | 6.66 | -29.06 | 9 | 1 |
| precentral | 4.14 | -31.21 | 10 | 1 |
| superiorfrontal | 0.52 | 3.03 | 1 | 1 |
| **FWF – left hemisphere** | | | | |
| caudalmiddlefrontal | 28.15 | 276.69 | 69 | 1 |
| lateraloccipital | 21.08 | -95.01 | 29 | 1 |
| superiortemporal | 12.99 | -93.14 | 29 | 1 |
| insula | 5.28 | -51.83 | 16 | 1 |
| **ODI – right hemisphere** | | | | |
| lateralorbitofrontal | 38.68 | 224.64 | 64.00 | 2.00 |
| inferiorparietal | 24.34 | -148.01 | 46.00 | 2.00 |
| supramarginal | 0.33 | 3.02 | 1.00 | 1.00 |
| **ODI – left hemisphere** | | | | |

| | | | | |
|---|---|---|---|---|
| isthmuscingulate | 27.24 | 306.21 | 80.00 | 1.00 |
| inferiorparietal | 12.88 | -65.67 | 20.00 | 1.00 |
| lateraloccipital | 5.93 | 0.30 | 8.00 | 2.00 |
| parsopercularis | 5.18 | -39.72 | 13.00 | 1.00 |
| posteriorcingulate | 3.52 | -34.51 | 11.00 | 1.00 |
| paracentral | 1.28 | 12.19 | 4.00 | 1.00 |

*Table S6.* Clusters exhibiting significant MD, NDI, FWF and ODI modifications at Day 5 when compared to baseline after cluster-wise correction brought together by region. The table represent changes for both groups. Annot = the name of the regions the maximum cluster value falls into using **Desikan-Killiany atlas**. Size ($mm^2$) = sum of the cluster sizes in millimeters square. WghtVtx = sum of the cluster weights (size*intensity), a negative weight indicate decrease in intensity while a positive value points towards an increase in intensity. NVtxs = sum of the number of vertices in the clusters from the region. NbClust = number of clusters in the labelled area.

### Table S7. Delayed structural changes and sleep-related effects in Pre-learning vs. Pre-relearning (Day 1 DWI1 vs. Day 5 DWI4 x RS vs. TMR) in the cortical ribbon

| POST-SLEEP DELAYED $DWI_4$-$DWI_1$; group differences (group TMR vs. RS) | | | | |
|---|---|---|---|---|
| **Annot** | **Size(mm^2)** | **WghtVtx** | **NVtxs** | **nbClust** |
| **ODI – right hemisphere** | | | | |
| cuneus | 61.37 | 302.82 | 82 | 1 |

*Table S7.* Only one cluster located in the right cuneus exhibited a significant ODI difference between both Conditions when comparing the delayed scan on Day 5 to baseline after cluster-wise correction. Annot = the name of the regions the maximum cluster value falls into using **Desikan-Killiany atlas**. Size ($mm^2$) = sum of the cluster sizes in millimeters square. WghtVtx = sum of the cluster weights (size*intensity), a negative weight indicate decrease in intensity while a positive value points towards an increase in intensity. NVtxs = sum of the number of vertices in the clusters from the region. NbClust = number of clusters in the labelled area.

### Table S8. Post-relearning short-term structural changes (Day 5; DWI4 vs. DWI5) in the cortical ribbon.

| RELEARNING $DWI_5$-$DWI_4$ for the whole sample (group RS + TMR) | | | | |
|---|---|---|---|---|
| **Annot** | **Size(mm^2)** | **WghtVtx** | **NVtxs** | **nbClust** |
| **MD – right hemisphere** | | | | |
| lateraloccipital | 497.20 | -2240.47 | 643.00 | 6.00 |
| precuneus | 237.66 | -2292.17 | 604.00 | 2.00 |
| superiorparietal | 217.82 | -2066.96 | 467.00 | 2.00 |
| isthmuscingulate | 195.11 | -2156.33 | 528.00 | 1.00 |
| fusiform | 147.41 | -1094.17 | 299.00 | 2.00 |
| bankssts | 142.86 | -1486.55 | 407.00 | 2.00 |
| posteriorcingulate | 113.37 | -1152.40 | 288.00 | 3.00 |
| postcentral | 75.48 | -654.89 | 186.00 | 4.00 |
| insula | 73.60 | -674.03 | 192.00 | 4.00 |
| medialorbitofrontal | 61.88 | -287.96 | 82.00 | 1.00 |

| | | | | |
|---|---|---|---|---|
| supramarginal | 41.44 | -379.35 | 109.00 | 5.00 |
| lingual | 41.39 | -143.33 | 41.00 | 2.00 |
| caudalmiddlefrontal | 32.87 | -216.88 | 64.00 | 3.00 |
| superiorfrontal | 28.52 | -197.22 | 57.00 | 2.00 |
| cuneus | 25.10 | -93.88 | 30.00 | 2.00 |
| caudalanteriorcingulate | 24.72 | -171.22 | 52.00 | 1.00 |
| rostralmiddlefrontal | 24.00 | -156.23 | 48.00 | 3.00 |
| paracentral | 23.78 | -286.86 | 76.00 | 1.00 |
| lateralorbitofrontal | 13.16 | -90.49 | 28.00 | 1.00 |
| precentral | 9.09 | -87.42 | 26.00 | 1.00 |
| parsopercularis | 3.34 | -30.78 | 10.00 | 1.00 |
| pericalcarine | 2.07 | -6.08 | 2.00 | 1.00 |
| middletemporal | 0.31 | -3.06 | 1.00 | 1.00 |
| **MD – left hemisphere** | | | | |
| inferiorparietal | 323.35 | -2191.13 | 595.00 | 3.00 |
| precentral | 154.48 | -1592.66 | 424.00 | 3.00 |
| posteriorcingulate | 141.37 | -1677.40 | 384.00 | 1.00 |
| precuneus | 91.08 | -695.72 | 202.00 | 4.00 |
| supramarginal | 87.13 | -938.24 | 265.00 | 3.00 |
| middletemporal | 83.77 | -719.41 | 201.00 | 2.00 |
| transversetemporal | 80.94 | -1115.45 | 228.00 | 1.00 |
| cuneus | 80.57 | -338.63 | 94.00 | 3.00 |
| rostralmiddlefrontal | 71.61 | -421.51 | 113.00 | 1.00 |
| superiorfrontal | 70.84 | -562.84 | 155.00 | 3.00 |
| insula | 44.78 | -436.74 | 128.00 | 4.00 |
| isthmuscingulate | 43.96 | -356.48 | 100.00 | 4.00 |
| postcentral | 43.77 | -329.24 | 100.00 | 4.00 |
| fusiform | 27.02 | -171.03 | 49.00 | 1.00 |
| caudalmiddlefrontal | 18.24 | -168.93 | 49.00 | 1.00 |
| superiorparietal | 16.78 | -113.81 | 36.00 | 3.00 |
| caudalanteriorcingulate | 14.98 | -93.09 | 30.00 | 2.00 |
| lingual | 1.98 | -9.15 | 3.00 | 1.00 |
| lateraloccipital | 0.66 | -3.05 | 1.00 | 1.00 |
| **NDI – right hemisphere** | | | | |
| fusiform | 203.60 | 1533.93 | 425.00 | 5.00 |
| lateraloccipital | 155.39 | 781.37 | 213.00 | 5.00 |
| parsopercularis | 154.99 | 1103.47 | 324.00 | 3.00 |
| precentral | 149.95 | 816.49 | 239.00 | 3.00 |
| caudalmiddlefrontal | 138.31 | 982.44 | 285.00 | 4.00 |
| rostralmiddlefrontal | 133.47 | 984.68 | 258.00 | 6.00 |
| cuneus | 124.21 | 549.20 | 162.00 | 2.00 |
| precuneus | 115.81 | 1116.75 | 321.00 | 3.00 |
| medialorbitofrontal | 91.05 | 476.77 | 129.00 | 1.00 |
| posteriorcingulate | 81.92 | 713.13 | 215.00 | 4.00 |
| superiorfrontal | 58.38 | 376.40 | 115.00 | 5.00 |
| parahippocampal | 53.44 | 306.45 | 88.00 | 1.00 |

| | | | | |
|---|---|---|---|---|
| inferiorparietal | 45.01 | 379.49 | 98.00 | 1.00 |
| isthmuscingulate | 40.67 | 367.46 | 100.00 | 3.00 |
| superiorparietal | 34.84 | 316.65 | 87.00 | 1.00 |
| insula | 34.61 | 332.01 | 94.00 | 3.00 |
| lateralorbitofrontal | 19.01 | 141.04 | 40.00 | 1.00 |
| bankssts | 14.13 | 130.38 | 41.00 | 3.00 |
| caudalanteriorcingulate | 12.23 | 88.19 | 28.00 | 1.00 |
| middletemporal | 11.39 | 58.43 | 17.00 | 1.00 |
| supramarginal | 8.42 | 70.61 | 23.00 | 2.00 |
| **NDI – left hemisphere** | | | | |
| isthmuscingulate | 251.72 | 2266.81 | 604.00 | 3.00 |
| rostralmiddlefrontal | 237.01 | 1468.25 | 376.00 | 2.00 |
| posteriorcingulate | 134.78 | 1353.95 | 374.00 | 1.00 |
| inferiorparietal | 96.26 | 724.80 | 200.00 | 5.00 |
| middletemporal | 74.82 | 763.37 | 195.00 | 2.00 |
| fusiform | 72.09 | 474.73 | 129.00 | 1.00 |
| parahippocampal | 63.65 | 451.34 | 126.00 | 2.00 |
| supramarginal | 63.58 | 694.65 | 190.00 | 1.00 |
| superiorfrontal | 60.10 | 449.00 | 124.00 | 3.00 |
| caudalmiddlefrontal | 35.45 | 266.54 | 79.00 | 2.00 |
| insula | 31.29 | 281.82 | 85.00 | 4.00 |
| superiorparietal | 19.17 | 189.28 | 56.00 | 1.00 |
| inferiortemporal | 15.19 | 84.49 | 25.00 | 1.00 |
| transversetemporal | 14.46 | 154.05 | 41.00 | 1.00 |
| bankssts | 11.99 | 88.60 | 27.00 | 1.00 |
| precentral | 7.15 | 81.42 | 26.00 | 1.00 |
| postcentral | 2.69 | 18.20 | 6.00 | 1.00 |
| **FWF – right hemisphere** | | | | |
| cuneus | 181.37 | -766.23 | 209.00 | 3.00 |
| isthmuscingulate | 52.72 | -508.18 | 133.00 | 1.00 |
| insula | 48.06 | -684.32 | 169.00 | 1.00 |
| superiorparietal | 46.41 | -332.33 | 98.00 | 1.00 |
| postcentral | 45.59 | -363.98 | 103.00 | 3.00 |
| superiorfrontal | 42.55 | -272.17 | 82.00 | 3.00 |
| lateraloccipital | 35.21 | -197.36 | 53.00 | 1.00 |
| pericalcarine | 28.30 | -89.18 | 28.00 | 2.00 |
| bankssts | 16.00 | -127.58 | 39.00 | 3.00 |
| precuneus | 12.01 | -69.24 | 21.00 | 2.00 |
| supramarginal | 6.09 | -40.08 | 13.00 | 2.00 |
| inferiorparietal | 2.12 | -16.29 | 5.00 | 2.00 |
| fusiform | 1.82 | -9.24 | 3.00 | 1.00 |
| entorhinal | 0.52 | -3.00 | 1.00 | 1.00 |
| **FWF – left hemisphere** | | | | |
| precentral | 50.11 | -538.70 | 147.00 | 1.00 |
| lateraloccipital | 35.88 | -171.41 | 52.00 | 1.00 |
| cuneus | 30.76 | -157.74 | 47.00 | 1.00 |

| | | | | |
|---|---|---|---|---|
| precuneus | 27.25 | -212.63 | 59.00 | 3.00 |
| insula | 24.94 | -228.34 | 66.00 | 2.00 |
| inferiorparietal | 24.89 | -135.01 | 42.00 | 2.00 |
| postcentral | 17.56 | -130.42 | 39.00 | 1.00 |
| superiorparietal | 7.70 | -51.96 | 17.00 | 2.00 |
| transversetemporal | 4.74 | -40.45 | 13.00 | 1.00 |
| supramarginal | 4.54 | -31.73 | 10.00 | 1.00 |
| pericalcarine | 0.77 | -6.04 | 2.00 | 1.00 |
| **ODI – right hemisphere** | | | | |
| lingual | 89.03 | 321.53 | 89.00 | 3.00 |
| parstriangularis | 29.00 | -126.05 | 40.00 | 2.00 |
| inferiorparietal | 28.07 | 219.45 | 63.00 | 3.00 |
| rostralmiddlefrontal | 25.88 | -150.51 | 45.00 | 1.00 |
| precuneus | 22.43 | 110.42 | 34.00 | 1.00 |
| supramarginal | 13.94 | 97.77 | 30.00 | 1.00 |
| insula | 12.78 | 117.92 | 35.00 | 1.00 |
| caudalmiddlefrontal | 10.55 | -50.69 | 16.00 | 1.00 |
| middletemporal | 8.57 | 38.27 | 12.00 | 2.00 |
| lateraloccipital | 6.37 | 45.26 | 14.00 | 1.00 |
| bankssts | 5.10 | 33.69 | 11.00 | 1.00 |
| superiorparietal | 3.33 | 24.48 | 8.00 | 1.00 |
| paracentral | 2.41 | -18.82 | 6.00 | 1.00 |
| medialorbitofrontal | 1.01 | -6.02 | 2.00 | 2.00 |
| **ODI – left hemisphere** | | | | |
| precuneus | 19.34 | 140.47 | 42 | 1 |
| superiorfrontal | 15.39 | 108.57 | 32 | 1 |
| bankssts | 10.74 | 91.10 | 27 | 1 |
| inferiorparietal | 10.08 | 85.57 | 25 | 1 |
| rostralmiddlefrontal | 6.55 | 28.73 | 9 | 1 |
| postcentral | 2.65 | 18.71 | 6 | 1 |
| precentral | 1.73 | 12.16 | 4 | 1 |
| lateraloccipital | 0.53 | 3.04 | 1 | 1 |

*Table S8.* Clusters showing significant MD, NDI, FWF and ODI changes in both groups following relearning at Day 5 after cluster-wise correction gathered by cortical region. The table represent changes for both groups. Annot = the name of the regions the maximum cluster value falls into using Desikan-Killiany atlas. Size ($mm^2$) = sum of the cluster sizes in millimeters square. WghtVtx = sum of the cluster weights (size*intensity), a negative weight indicate decrease in intensity while a positive value points towards an increase in intensity. NVtxs = sum of the number of vertices in the clusters from the region. NbClust = number of clusters in the labelled area.

### Table S9. Post-relearning short-term structural changes and sleep-related effects (Day 5; DWI4 vs. DWI5 x RS vs. TMR) in the cortical ribbon.

| **RELEARNING $DWI_5$-$DWI_4$; group differences (group TMR vs. RS)** | | | | |
|---|---|---|---|---|
| **Annot** | **Size(mm^2)** | **WghtVtx** | **NVtxs** | **nbClust** |

| ODI – left hemisphere | | | | |
|---|---|---|---|---|
| superiorfrontal | 80.13 | -475.95 | 126 | 1 |

*Table S9.* One cluster located in the left superior frontal gyrus displayed a significant ODI difference between both Conditions when comparing the post-relearning scan to the pre-relearning scan on Day 5 after cluster-wise correction. Annot = the name of the regions the maximum cluster value falls into using **Desikan-Killiany atlas**. Size ($mm^2$) = sum of the cluster sizes in millimeters square. WghtVtx = sum of the cluster weights (size*intensity), a negative weight indicate decrease in intensity while a positive value points towards an increase in intensity. NVtxs = sum of the number of vertices in the clusters from the region. NbClust = number of clusters in the labelled area.

### Table S10. Summary of TMR studies in the motor memory domain.

| Source | Task | Cueing modality | Targeted sleep stage | Results |
|---|---|---|---|---|
| Schönauer et al., 2014 [13] | SRTT | auditory | All | Benefit (accuracy) after 3 hours of TMR (2h cueing – 1h sleep); gains = full night of regular sleep |
| Rakowska et al., 2021 [14] | SRTT | auditory | N2 & N3 | Benefit at 10 days only |
| Nicolas et al., 2024 [15] | SRTT | auditory | N2 & N3 | No benefit (older adults) |
| Antony et al., 2012 [16] | FTT | auditory | N3 | Benefit accuracy |
| Cousins et al., 2014 [17] | SRTT | auditory | N3 | Benefit (RT) & increased explicit recall |
| Cousins et al., 2016 [18] | SRTT | auditory | N3 | Benefit (RT and accuracy) |
| Koopman et al., 2020 [19] | SRTT | auditory | N3 | Benefit non-dominant hand only |
| Johnson et al., 2018 [20] | Target throwing | auditory | All | Benefit (throwing accuracy) |
| Johnson et al., 2019 [21] | Target throwing | auditory | All | Benefit (throwing accuracy) |
| Johnson et al., 2020 [22] | Target throwing | auditory | All | Benefit (throwing accuracy; older adults) |
| Cheng et al., 2021 [23] | Complex motor task | auditory | N3 | Benefit (time to target) |
| Laventure et al., 2016 [6] | FTT | olfactory | N2 | Benefit (GPI) |
| Laventure et al., 2018 [24] | FTT | olfactory | N2 | Benefit (GPI) |
| Rasch et al., 2007 [25] | FTT | olfactory | N3 | No benefit |
| Diekelmann et al., 2016 [26] | SRTT | olfactory | N3 | No benefit; but increased explicit knowledge in men (not women) |

| | | | | |
|---|---|---|---|---|
| Pereira et al., 2017 [27] | FTT | tactile | N2 | No benefit |
| Veldman et al., 2021 [28] | SRTT | tactile | N2 & N3 | No benefit |

*Table S10.* Overview of all the experiments that, to the best of our knowledge, combined Non-Rapid Eye Movement (NREM) Targeted Memory Reactivation (TMR) protocols and motor memory, ranked by cueing modality and targeted sleep stage. Main results are also summarized in the last column. SRTT = Serial Reaction Time Task; FTT = Finger Taping Task; RT = Reaction Time; GPI = Global Performance Index.

### Bibliography SI


[1] L. L. Jacoby, "A process dissociation framework: Separating automatic from intentional uses of memory," *J. Mem. Lang.*, vol. 30, no. 5, pp. 513–541, Oct. 1991, doi: 10.1016/0749-596X(91)90025-F.

[2] A. Destrebecqz *et al.*, "The neural correlates of implicit and explicit sequence learning: Interacting networks revealed by the process dissociation procedure," *Learn. Mem. Cold Spring Harb. N*, vol. 12, no. 5, pp. 480–490, Sep. 2005, doi: 10.1101/LM.95605.

[3] J. Fonollosa, E. Neftci, and M. Rabinovich, "Learning of Chunking Sequences in Cognition and Behavior," *PLOS Comput. Biol.*, vol. 11, no. 11, p. e1004592, Nov. 2015, doi: 10.1371/JOURNAL.PCBI.1004592.

[4] K. Sakai, K. Kitaguchi, and O. Hikosaka, "Chunking during human visuomotor sequence learning," *Exp. Brain Res.*, vol. 152, no. 2, pp. 229–242, Sep. 2003, doi: 10.1007/S00221-003-1548-8.

[5] A. Cleeremans and J. L. McClelland, "Learning the Structure of Event Sequences," *J. Exp. Psychol. Gen.*, vol. 120, no. 3, pp. 235–253, 1991, doi: 10.1037/0096-3445.120.3.235.

[6] S. Laventure, S. Fogel, and O. Lungu, "NREM2 and Sleep Spindles Are Instrumental to the Consolidation of Motor Sequence Memories," pp. 1–27, 2016, doi: 10.1371/journal.pbio.1002429.

[7] D. C. 't Wallant *et al.*, "Automatic artifacts and arousals detection in whole-night sleep EEG recordings," *J. Neurosci. Methods*, vol. 258, pp. 124–133, Jan. 2016, doi: 10.1016/j.jneumeth.2015.11.005.

[8] J. A. Desjardins and S. J. Segalowitz, "Deconstructing the early visual electrocortical responses to face and house stimuli," *J. Vis.*, vol. 13, no. 5, p. 22, Apr. 2013, doi: 10.1167/13.5.22.

[9] M. Mölle, L. Marshall, S. Gais, and J. Born, "Grouping of Spindle Activity during Slow Oscillations in Human Non-Rapid Eye Movement Sleep," *J. Neurosci.*, vol. 22, no. 24, pp. 10941–10947, Dec. 2002, doi: 10.1523/JNEUROSCI.22-24-10941.2002.

[10] J. Carrier *et al.*, "Sleep slow wave changes during the middle years of life," *Eur. J. Neurosci.*, vol. 33, no. 4, pp. 758–766, Feb. 2011, doi: 10.1111/j.1460-9568.2010.07543.x.

[11] M. Massimini, R. Huber, F. Ferrarelli, S. Hill, and G. Tononi, "The sleep slow oscillation as a traveling wave," *J. Neurosci. Off. J. Soc. Neurosci.*, vol. 24, no. 31, pp. 6862–6870, Aug. 2004, doi: 10.1523/JNEUROSCI.1318-04.2004.

[12] M. Basner, S. Mcguire, N. Goel, H. Rao, and D. F. Dinges, "A new likelihood ratio metric for the psychomotor vigilance test and its sensitivity to sleep loss," *J. Sleep Res.*, vol. 24, no. 6, pp. 702–713, Dec. 2015, doi: 10.1111/JSR.12322.

[13] M. Schönauer, T. Geisler, and S. Gais, "Strengthening Procedural Memories by Reactivation in Sleep," *J. Cogn. Neurosci.*, vol. 26, no. 1, pp. 143–153, Jan. 2014, doi: 10.1162/JOCN_A_00471.

[14] M. Rakowska, M. E. A. Abdellahi, P. Bagrowska, M. Navarrete, and P. A. Lewis, "Long term effects of cueing procedural memory reactivation during NREM sleep," *NeuroImage*, vol. 244, p. 118573, Dec. 2021, doi: 10.1016/J.NEUROIMAGE.2021.118573.

[15] J. Nicolas, J. Carrier, S. P. Swinnen, J. Doyon, G. Albouy, and B. R. King, "Targeted memory reactivation during post-learning sleep does not enhance motor memory consolidation in older adults," *J. Sleep Res.*, vol. 33, no. 1, p. e14027, Feb. 2024, doi: 10.1111/JSR.14027.

[16] J. W. Antony, E. W. Gobel, J. K. O'Hare, P. J. Reber, and K. A. Paller, "Cued memory reactivation during sleep influences skill learning," *Nat. Neurosci. 2012 158*, vol. 15, no. 8, pp. 1114–1116, Jun. 2012, doi: 10.1038/nn.3152.

[17] J. N. Cousins, W. El-Deredy, L. M. Parkes, N. Hennies, and P. A. Lewis, "Cued memory reactivation during slow-wave sleep promotes explicit knowledge of a motor sequence," *J. Neurosci. Off. J. Soc. Neurosci.*, vol. 34, no. 48, pp. 15870–15876, Nov. 2014, doi: 10.1523/JNEUROSCI.1011-14.2014.

[18] J. N. Cousins, W. El-Deredy, L. M. Parkes, N. Hennies, and P. A. Lewis, "Cued Reactivation of Motor Learning during Sleep Leads to Overnight Changes in Functional Brain Activity and Connectivity," *PLoS Biol.*, vol. 14, no. 5, May 2016, doi: 10.1371/journal.pbio.1002451.

[19] A. C. M. Koopman[1] *et al.*, “Targeted memory reactivation of a serial reaction time task in SWS, but not REM, preferentially benefits the non-dominant hand,” *bioRxiv*, p. 2020.11.17.381913, Nov. 2020, doi: 10.1101/2020.11.17.381913.

[20] B. P. Johnson, S. M. Scharf, and K. P. Westlake, “Targeted Memory Reactivation During Sleep, But Not Wake, Enhances Sensorimotor Skill Performance: A Pilot Study,” *J. Mot. Behav.*, vol. 50, no. 2, pp. 202–209, Mar. 2018, doi: 10.1080/00222895.2017.1327411.

[21] B. P. Johnson, S. M. Scharf, A. C. Verceles, and K. P. Westlake, “Use of targeted memory reactivation enhances skill performance during a nap and enhances declarative memory during wake in healthy young adults,” *J. Sleep Res.*, vol. 28, no. 5, 2019, doi: 10.1111/jsr.12832.

[22] B. P. Johnson, S. M. Scharf, A. C. Verceles, and K. P. Westlake, “Sensorimotor performance is improved by targeted memory reactivation during a daytime nap in healthy older adults,” *Neurosci. Lett.*, vol. 731, Jul. 2020, doi: 10.1016/j.neulet.2020.134973.

[23] L. Y. Cheng, T. Che, G. Tomic, M. W. Slutzky, and K. A. Paller, “Memory Reactivation during Sleep Improves Execution of a Challenging Motor Skill,” *J. Neurosci. Off. J. Soc. Neurosci.*, vol. 41, no. 46, pp. 9608–9616, Nov. 2021, doi: 10.1523/JNEUROSCI.0265-21.2021.

[24] S. Laventure *et al.*, “Beyond spindles: interactions between sleep spindles and boundary frequencies during cued reactivation of motor memory representations,” *Sleep*, vol. 41, no. 9, pp. 1–14, Sep. 2018, doi: 10.1093/SLEEP/ZSY142.

[25] B. Rasch, C. Büchel, S. Gais, and J. Born, “Odor cues during slow-wave sleep prompt declarative memory consolidation,” *Science*, vol. 315, no. 5817, pp. 1426–1429, Mar. 2007, doi: 10.1126/science.1138581.

[26] S. Diekelmann, J. Born, and B. Rasch, “Increasing Explicit Sequence Knowledge by Odor Cueing during Sleep in Men but not Women,” *Front. Behav. Neurosci.*, vol. 10, no. APRIL, Apr. 2016, doi: 10.3389/FNBEH.2016.00074.

[27] S. I. R. Pereira, F. Beijamini, F. D. Weber, R. A. Vincenzi, F. A. C. da Silva, and F. M. Louzada, “Tactile stimulation during sleep alters slow oscillation and spindle densities but not motor skill,” *Physiol. Behav.*, vol. 169, pp. 59–68, Feb. 2017, doi: 10.1016/J.PHYSBEH.2016.11.024.

[28] M. P. Veldman, N. Dolfen, M. A. Gann, J. Carrier, B. R. King, and G. Albouy, "Somatosensory Targeted Memory Reactivation Modulates Oscillatory Brain Activity but not Motor Memory Consolidation," *Neuroscience*, vol. 465, pp. 203–218, Jun. 2021, doi: 10.1016/J.NEUROSCIENCE.2021.03.027.
